# Investigating the versatility of cytochalasan cytochrome P450 monooxygenases using combinatorial biosynthesis reveals stereochemical restrictions

**DOI:** 10.64898/2026.02.28.708751

**Authors:** Lei Li, Tahir Ali, Jacob Goralczyk, Sameera Jayasundara, Ayan Paul, Marcelo Rodrigues de Amorim, Christine Beemelmanns, Elizabeth Skellam

**Affiliations:** Department of Chemistry, University of North Texas, 1155 Union Circle, Denton, TX 76203, USA; BioDiscovery Institute, University of North Texas, 1155 Union Circle, Denton, TX 76203, USA; Instituto de Química de São Carlos, Universidade de São Paulo, CP 780, CEP 13560-970, São Carlos, SP, Brazil; Helmholtz Institute for Pharmaceutical Research Saarland (HIPS), Helmholtz Centre for Infection Research (HZI), Campus E8.1, 66123 Saarbrücken, Germany; Pharma Science Hub (PSH), Saarland University, 66123 Saarbrücken, Germany

**Keywords:** Biocatalysis, combinatorial biosynthesis, cytochalasans, cytochrome P450 monooxygenases

## Abstract

**Background:** Cytochalasans are a large family of fungal metabolites which inhibit actin polymerization and ultimately lead to a broad range of biological effects in different assays. Investigations into the biosynthesis of cytochalasans has revealed that the cytochrome P450 monooxygenase (P450s) tailoring enzymes possess a somewhat relaxed substrate-specificity and may accept structurally-related intermediates for oxidation, partly explaining the variety of structural variations observed in this family of molecules. In this study, we investigate a broad range of P450 enzymes *via* combinatorial biosynthesis to better understand their substrate scope and potential applications as biocatalysts.

**Results:** Genome mining enabled us to identify cryptic cytochalasan biosynthetic gene clusters (BGCs) in six different species of fungi, each with at least two P450 enzymes encoded. Comparative genomics identified a cryptic thioredoxin-like enzyme encoded in cytochalasan BGCs that co-occurs with the gene encoding a Baeyer-Villiger monooxygenase. Heterologous expression of seven P450s in *Magnaporthe grisea* mutant strains, lacking P450s required for pyrichalasin H biosynthesis, enabled functional characterization of three P450s, two of which were previously cryptic. The experimental results, combined with phylogenetic analysis of the P450 sequences, reveal subtle information regarding the structures of the associated cytochalasans and begins to explain why some P450s are inactive on the substrates available to them.

**Conclusions:** The P450 enzymes involved in cytochalasan biosynthesis are known to be site-selective in their native host but also possess intrinsic promiscuity due to being able to modify structurally-related analogues. By investigating a diverse set of P450s from characterized and cryptic BGCs, we were able to identify that the stereochemistry of functional groups around the cytochalasan backbone is more restrictive than the size of the macrocycle when introducing the P450 enzyme to non-native substrates.

## Introduction

Cytochalasans are a large and diverse family of fungal natural products with a broad range of biological functions, including: inhibition of actin polymerization; cytotoxic and cytostatic effects; anti-microbial, anti-viral, and immunosuppressant activities.[1,2] Structurally, cytochalasans consist of a tricyclic core with a perhydroisoindolone moiety derived from an amino acid fused to a polyketide chain, synthesized by a polyketide synthase / non-ribosomal peptide synthetase (PKS-NRPS).[3,4] The wide range of structural diversity observed reflects the different amino acids incorporated (including alanine, leucine, (*O*-methyl)tyrosine, phenylalanine, tryptophan, and valine); the length of the polyketide chain (C_12_ – C_20_); and the modification reactions of the carbon skeleton (including hydroxylations, epoxidations, acylations, and oxygen insertions). [5] Subtle differences in structural features drastically alter the reported biological activities of this family of molecules (Figure 1).[6]

**Figure 1.**
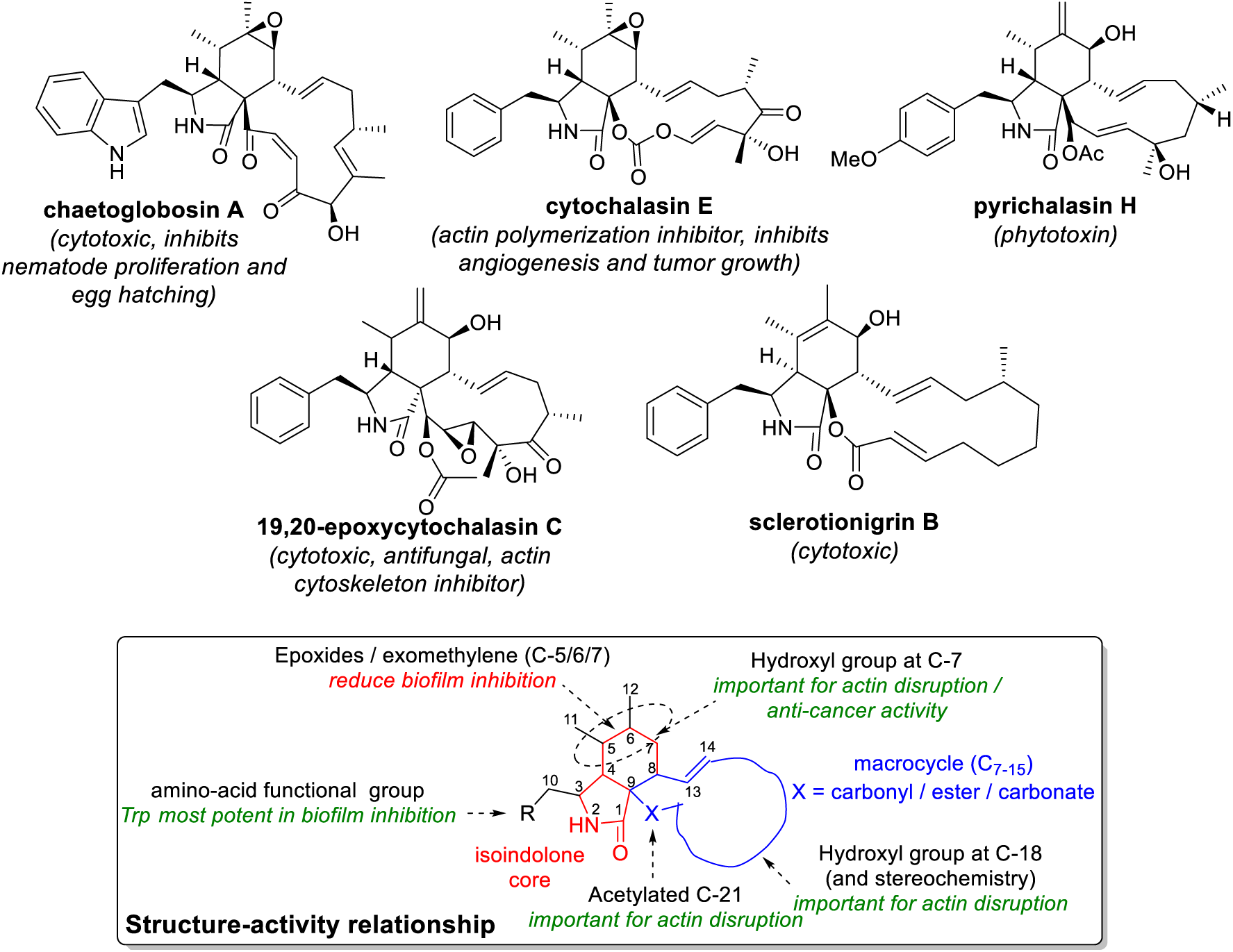
Examples of cytochalasan structures and bioactivities important to this study.

In the last 15 years, significant advances in understanding the enzymes required for the biosynthesis of cytochalasans have been made: a PKS-NRPS and *trans*-acting enoyl reductase (*trans*-ER) collaborate to synthesize a linear polyketide-peptide hybrid [7–9], released as an aldehyde [10]; an α,b-hydrolase (HYD) is involved in catalyzing the formation of the pyrrolidone core and preventing tautomerization [11]; and, a Diels-Alderase (DA) catalyzes formation of the perhydroisoindolone core [12]. After the tricyclic core has been synthesized (Scheme 1A), tailoring enzymes further modify and diversify the structure such as: cytochrome P450 monooxygenases (P450s) which epoxidize the cyclohexene moiety or hydroxylate the macrocycle [13], some of which act in an iterative fashion introducing two consecutive hydroxy groups at adjacent carbon atoms (Scheme 1B) [7]; Baeyer-Villiger monooxygenases (BVMO) which catalyze one or two oxygen insertions within the macrocycle resulting in esters or carbonates respectively [14]; oxidoreductases (OXR) which oxidize alcohols to ketones [7], reduce ketones to alcohols [15], or reduce double bonds to single bonds [16]; and methyl- and acetyltransferases [15].

**Scheme 1.**
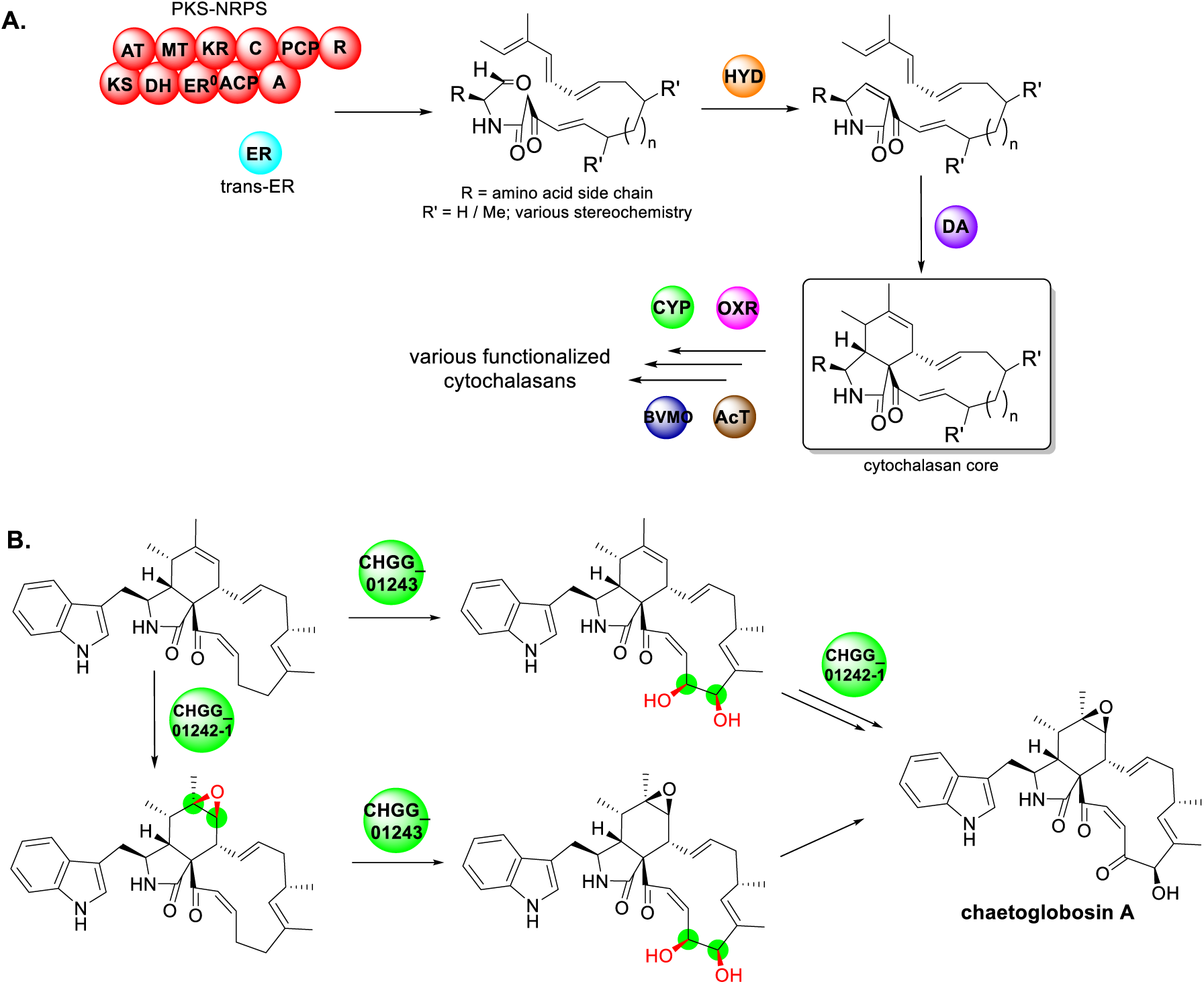
A) Overview of the biosynthesis of the cytochalasan core and subsequent tailoring reactions; B) Example of oxidative modifications performed by cytochrome P450 monooxygenases during chaetoglobosins biosynthesis, demonstrating that some P450s act iteratively and many accept more than one substrate [7]. PKS-NRPS domain abbreviations: KS = ketosynthase; AT = acyltransferase; DH = dehydratase; MT = methyltransferase; ER^0^ = enoylreductase (inactive); KR = ketoreductase; ACP = acyl carrier protein; C = condensstaion; A = adenylation; PCP = peptidyl carrier protein; R = reductase.

Furthermore, several non-enzymatic modifications have also been studied including dimerization and polymerization with non-cytochlasans producing merocytochalasans [16,17]; acid-catalyzed rearrangements [18,19]; and acid-catalyzed macrocycle cleavage and heterocycle formation [20] (Scheme S1). Finally, titers of cytochalasans have been increased through transcription factor over-expression [21, 14].

Although chemical synthesis of cytochalasans is possible, it remains challenging due to the number of steps needed, the requirement to protect and subsequently deprotect functional groups, as well as stereochemical considerations of late-stage functionalization [22,23]. Chemoenzymatic synthesis offers a route to more efficient syntheses, enabling rapid conversion of unfunctionalized molecules, provided that the tailoring enzymes have a broad enough substrate scope to be used across diverse substrates [24].

In prior work, we reported mutasynthesis as an approach for the rapid generation of novel and unnatural cytochalasans [15,25] and developed a combinatorial biosynthesis platform that enabled us to investigate the function of non-native cytochalasan P450s in a fungal host resulting in the generation of new cytochalasans, the restoration of pyrichalasin H, or no change in functionality of the cytochalasan substrates [13,26]. In this study, we extend our platform to investigate P450s from a wider range of fungal species to identify biocatalysts capable of oxidatively modifying non-native cytochalasan backbones, that possess different macrocycle sizes, functional groups, and stereocenters.

## Results

### Genome mining identifies cryptic cytochalasan BGCs, a highly conserved thioredoxin-like gene, and leads to discovery of a putative cytochalasan from *Aspergillus heteromorphus* CBS 117.55

We performed genome mining using the essential DA sequence (PyiF; NCBI accession number QCS37516.1) as a query to identify several commercially available fungi that possess a putative cytochalasan BGC, including: *Aspergillus heteromorphus* CBS 117.55, *Aspergillus sclerotioniger* CBS 115572, *Colletotrichum spinosum* CBS 515.97, *Colletotrichum sidae* CBS 518.97, *Metarhizium brunneum* ARSEF 3297, *Metarhizium robertsii* ARSEF 23. The cryptic cytochalasan BGCs were manually annotated using FGENESH and renamed *ahe*, *asc*, *csp*, *csi*, *mbr*, and *mro* respectively (Figure 2; Table S3).

**Figure 2.**
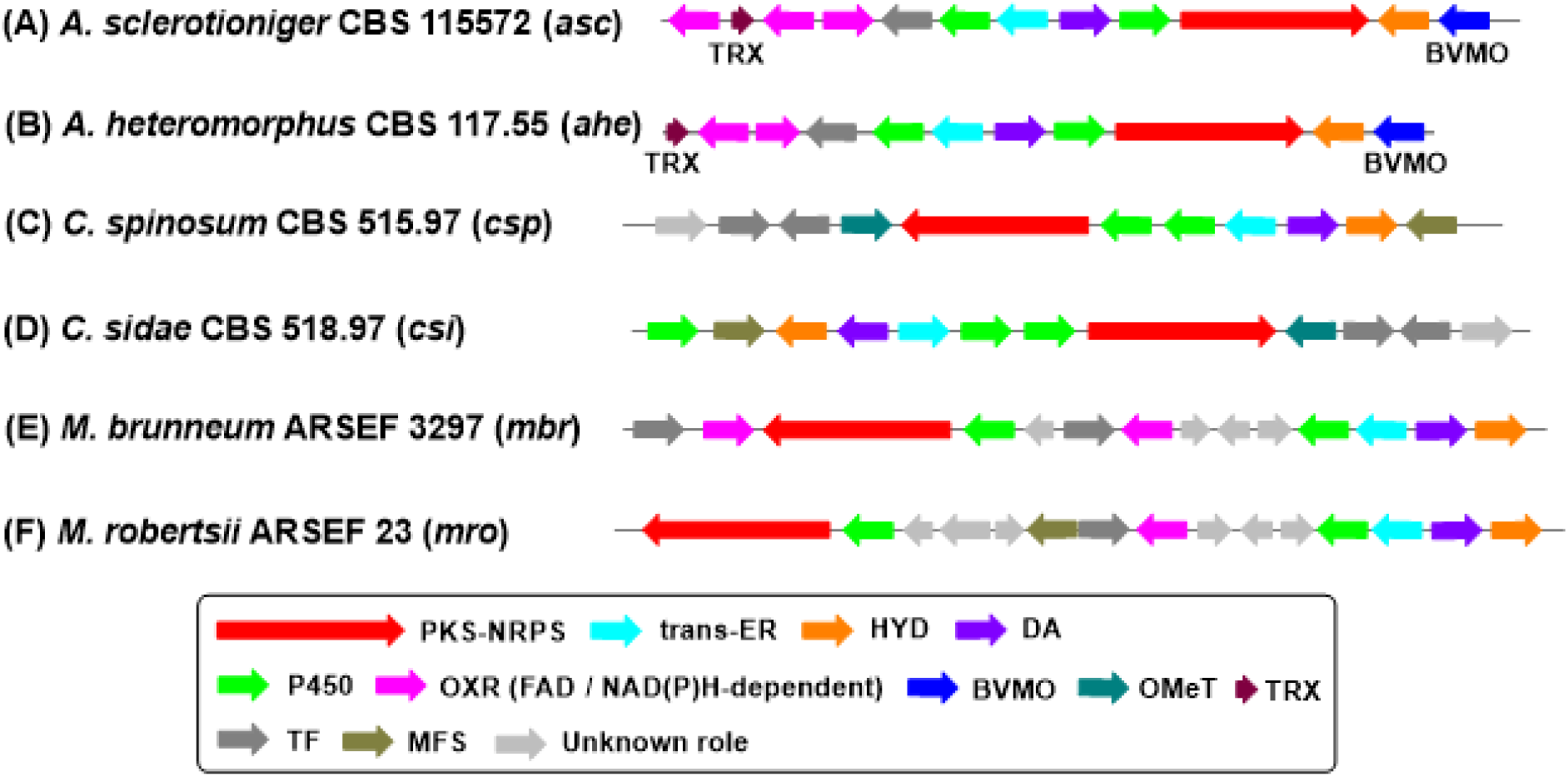
Cryptic cytochalasan BGCs identified in this study *via* genome mining. Genes are color-coded according to function, as described in the figure key.

All six BGCs encode the four core enzymes required for cytochalasan biosynthesis (PKS-NRPS, *trans*-ER, HYD and DA) [5], in addition to two P450s corresponding to macrocycle and cyclohexene regioselectivities, however, they vary considerably with the presence or absence of various tailoring enzymes, transcription factors (TF), and major facilitator superfamily (MFS) transporters (Table S3). In addition, we observed a gene encoding a thioredoxin (TRX)-like enzyme, first noticed in the *A. clavatus* NRRL1 *ccs* BGC (ACLA_078720) [3], that appears to correlate with the presence of a BVMO. Utilizing cblaster, the correlation between the TRX and BVMO was confirmed in various cryptic cytochalasan BGCs (Figure S3). Attempts to knock-out the TRX-gene (renamed to *ccsX*) in *A. clavatus* NRRL 1 to determine its function were unsuccessful based on PCR screening (Figure S6).

Other than *A. sclerotioniger* CBS 115572, which is known to produce several cytochalasans including sclerotionigrin A, sclerotionigrin B, and proxiphomin which derive from a C_18_ polyketide fused to phenylalanine (Figure S1) [27], none of the other fungi identified in this study have been reported as producing cytochalasans to the best of our knowledge. Examining the *A. sclerotioniger* CBS 115572 genome (GenBank accession number: MSFK00000000.1) *via* fungiSMASH and BLAST only one cytochalasan BGC was identified which contains the genes expected for the biosynthesis of sclerotonigrins A and B, as well as proxiphomin, linking this BGC bioinformatically to these cytochalasans.

We recently reported several natural products from *C. spinosum* CBS 515.97 and *A. heteromorphus* CBS 117.55 but did not identify any cytochalasans under the conditions used [28,29]. Due to the *ahe* BGC sharing some similarity with the *ccs* BGC in *A. clavatus* NRRL1 as well as the *asc* BGC (Figure S4), we cultivated *A. heteromorphus* CBS 117.55 under identical conditions to *A. clavatus* cytochalasin E production conditions [3], however comparison of the two extracts with a commercial standard of cytochalasan E confirmed that only *A. clavatus* NRRL1 produces this cytochalasan (Figure S15). Therefore, we cultivated *A. heteromorphus* CBS 117.55 in a broader range of media and performed RT-PCR of the PKS-NRPS and TRX genes from cDNA extracted from the resulting mycelia. Expression of the PKS-NRPS was confirmed in four out of five media used and TRX-expression was confirmed in all five media (Figure S8). In parallel, the cultures were extracted using EtOAc and subject to HR-LCMS/MS investigation (Figure S19-20). The resulting MS^2^ data was analyzed using MZmine and MS-DIAL, revealing a metabolite being detected in four of the five media with a *m/z* 496.2291 (molecular formula: C_28_H_34_NO_7_^+^, mass error: −7.85 ppm) feature in positive ion mode at 8.60 ± 0.02 min (Figure 3), reflecting the RT-PCR data. The same compound at *m/z* 494.2191 (chemical formula: C_28_H_32_NO_7_^-^, mass error: 1.42 ppm) was observed with a retention time of 8.60 ± 0.02 mins in negative ionization mode (Figure 3). Although this mass is similar to commercial cytochalasin E ([M+H]^+^ *m/z* = 496.2321), the clear retention time (9.65 mins) difference confirmed that this metabolite was not cytochalasin E.

**Figure 3.**
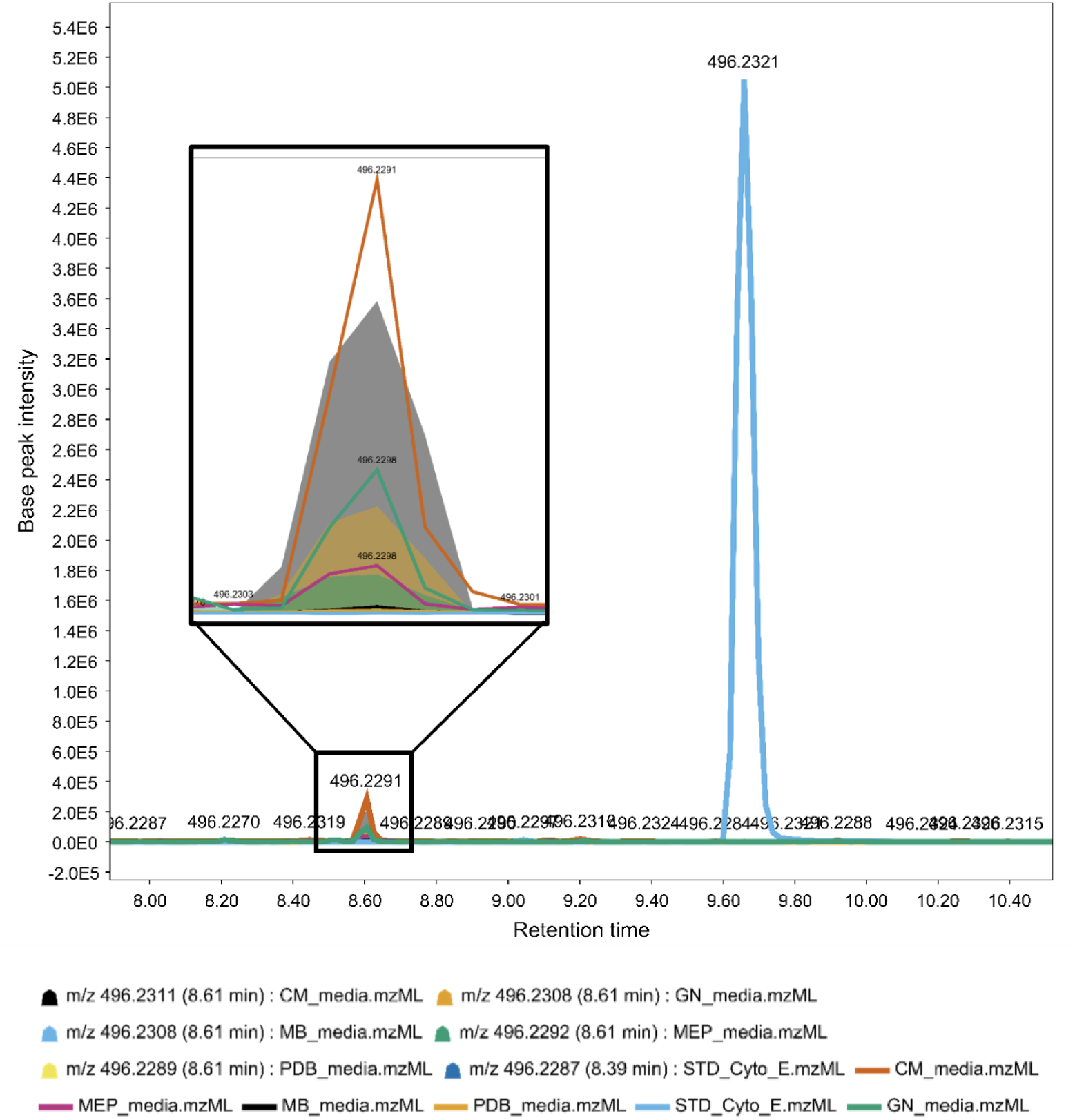
Overlay of extracted ion chromatogram for *m/z* 496.2291 showing a metabolite eluting at 8.61 min in four out of five fungal culture media with media-dependent signal intensity. Comparison with a cytochalasin E standard (0.1 mg/mL; *m/z* 496.2321; Rt 9.6 min) reveals a distinct retention time shift.

*In silico* spectral matching *via* MS-FINDER produced moderate similarity scores (6.94 and 6.90) to phenochalasin A and cytochalasin K_asp_ respectively, supporting its classification within the cytochalasan family. Notably, the highest similarity score (7.02) was obtained for cytochalasin E from the database, suggesting a closer match to this analogue. Since the *ahe* BGC shares more homology with the *asc* BGC than the *ccs* BGC, especially between the *trans*-ER and DA genes (Figure S4), this could indicate that the *ahe* cytochalasan molecule will be more similar to the proxiphomin and sclerotionigrins A/B backbones which contain two additional carbon atoms in the macrocycle but lack a pendant methyl group at C-18. Due to its low abundance, we have been unable to isolate the molecule for full structural elucidation but based on the interpretation of the observed MS^2^ fragmentation pattern propose that it could be derived from a C_18_ polyketide lacking methyl groups at the C-16 and C-18 positions (Figure S21).

### Phylogenetic analysis of P450s reveals structural information about cytochalasan structures

Multiple sequence alignments (MSA) of the cryptic P450s identified from genome mining and P450s from experimentally investigated cytochalasan BGCs identified the conserved CPG active site for co-ordinating heme and a conserved EXXR motif for structural integrity, indicating that all P450s should be functional (Figure 4A).[30] There are no major differences in primary sequences between P450s that were functional in previous heterologous expression studies and those that were not (Figure S5).[13]

**Figure 4.**
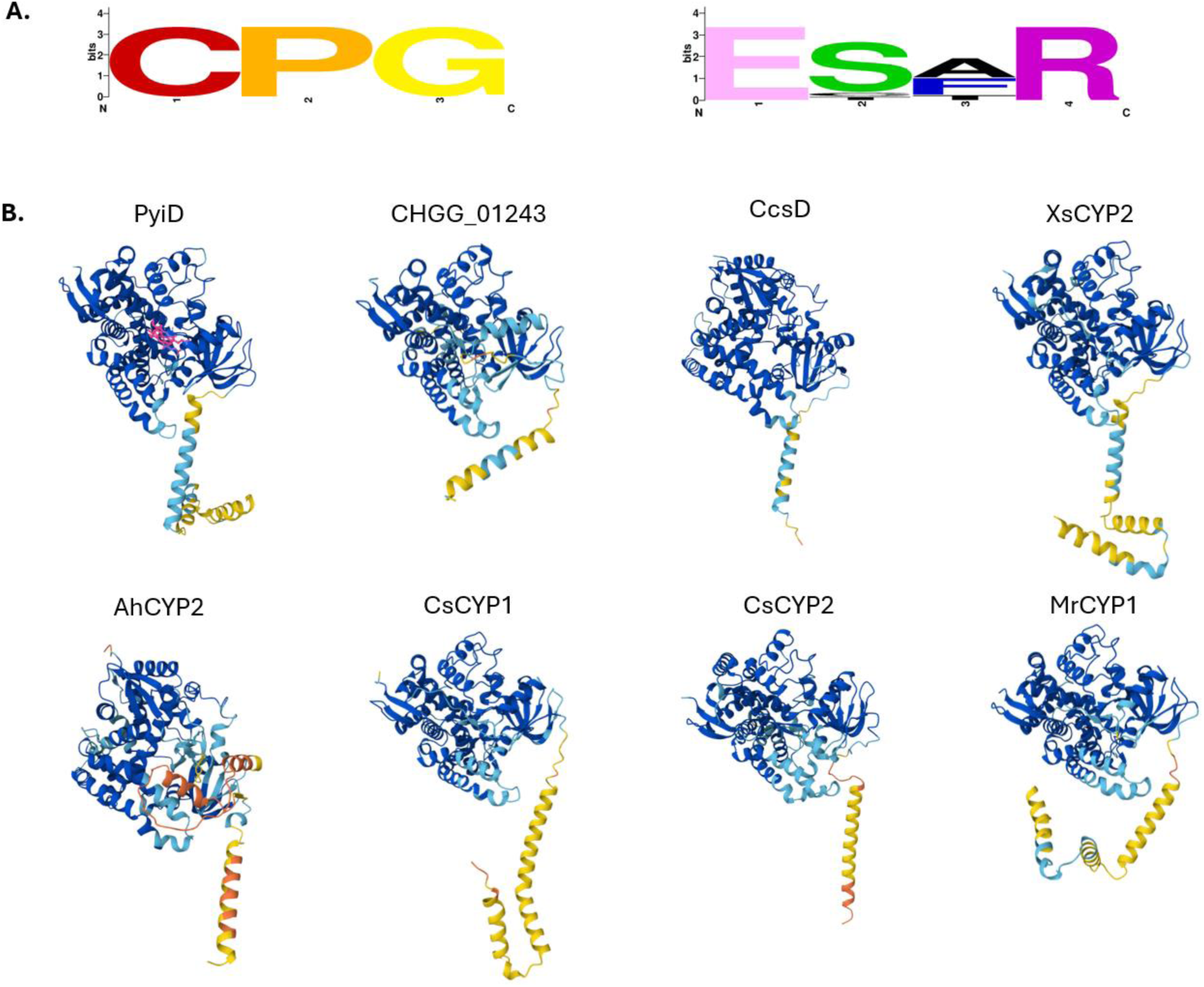
A) Analysis of conserved catalytic (CPG) and structural motifs (EXXR) within experimentally validated P450s created using WebLogo [31]; B) AlphaFold [32] models of P450s investigated in this study demonstrating variable α-helix length and structures. The heme co-factor is indicated in pink in the PyiD AlphaFold model. Note: the original truncated CcsD sequence is depicted.

Modelling the P450s using AlphaFold confirmed that the P450s under investigation have highly conserved structures with most variation observed within the length and orientation of the N-terminal α –helix (Figure 4B). In eukaryotes, P450s are typically anchored to the endoplasmic reticulum (ER) via this N-terminal α-helix. [33] Examining the MSA, most macrocycle P450s possess an extended N-terminal sequence of ∼ 20 - 60 amino acids which is not present in the cyclohexene P450 sequences (Figure S5). Noteably, CcsD also lacks this N-terminal sequence, however, re-examination of the *ccs* BGC revealed a 108 bp sequence upstream of ccsD apparently encoding the missing α-helix region (Figure S5).

The P450 sequences from the cryptic and experimentally validated BGCs [3,7,13,20] were used to construct a phylogenetic tree to determine if the different oxidative chemistries were reflected in their primary sequences (Figure 5). The P450s appear to clade according to their regiochemistry *e.g.* cyclohexene *vs.* macrocycle oxidative modifications, and the sub-clades reflect structural information regarding the chain length of the polyketide core and the type of amino acid incorporated, as well as the presence or absence of specific tailoring enzymes.

**Figure 5.**
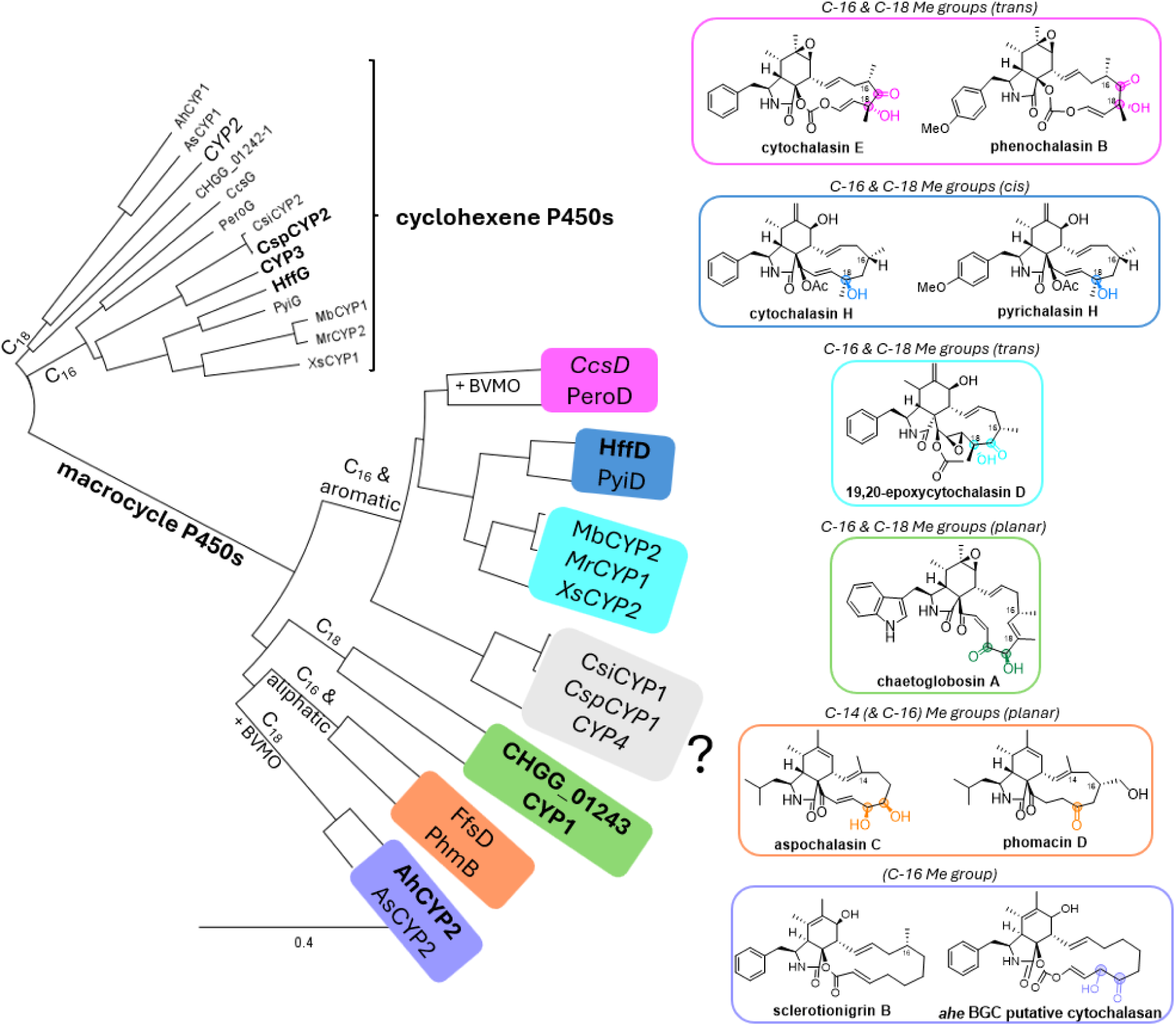
Phylogenetic tree of P450s involved in cytochalasan biosynthesis, distinguishing between cyclohexene and macrocycle associated enzymes. The oxidized C-H positions around the macrocycle are color-coded to identify the sub-clade of the corresponding P450 enzyme. P450s that have been shown to be functional in heterologous hosts are displayed in bold type; P450s that were investigated in a heterologous host but were not functional are displayed in italics. Structures for all cytochalasans identified from a given BGC are shown in Figures S1 and S2 for clarity.= No cytochalasan molecule has been isolated from BGCs associated with CsiCYP1, CspCYP1, and CYP4.

### Heterologous expression of macrocycle P450s from experimentally validated and cryptic cytochalasan BGCs

The cytochalasans chaetoglobosin A, cytochalasin E, and 19,20-epoxycytochalasin D possess oxidative modifications at two adjacent positions in the macrocycle (Figure 5) and have been experimentally or bioinformatically linked to a BGC. In chaetoglobosin A biosynthesis, CHGG_01243 is the P450 that introduces two consecutive (or iterative) hydroxylations at positions C-19 and C-20 (Scheme 1B). [7] The P450s that introduce the oxidative modifications at C-17 and C-18 during cytochalasin E/K (Figure 5, Figure S2) and 19,20-epoxycytochalasin C/D biosynthesis are currently unknown [3,34], but our phylogenetic analysis indicates that CcsD and XsCYP2 respectively may be responsible, suggesting that they may also be iterative P450s (Figure 5).

To study the substrate promiscuity of these enzymes more in detail, the gene sequences encoding these P450s were synthesized by a commercial vendor and individually expressed under the *amyB* promoter in M*agnaporthe grisea* Δ*p*yiD, a heterologous host that produces pyrichalasin H analogues lacking a hydroxy group at C-18 (Figure 5). [13,15] Transformants were selected three times on TNK-(SU)-CP agar containing the antibiotic glufosinate (basta), before being cultivated in liquid DPY medium for 7 days, and the EtOAc extracts being analyzed by LCMS. Heterologous expression of the iterative hydroxylase CHGG_01243 from *Chaetomium globosum* CBS 148.51 in *M. grisea* Δ*p*yi*D* restored pyrichalasin H production in eight out of eleven transformants (Figure 6B), as determined by LCMS comparison with a pyrichalasin H standard (Figures S28 – S34), however dihydroxylation of the pyrichalasin core structure was not observed.

**Figure 6.**
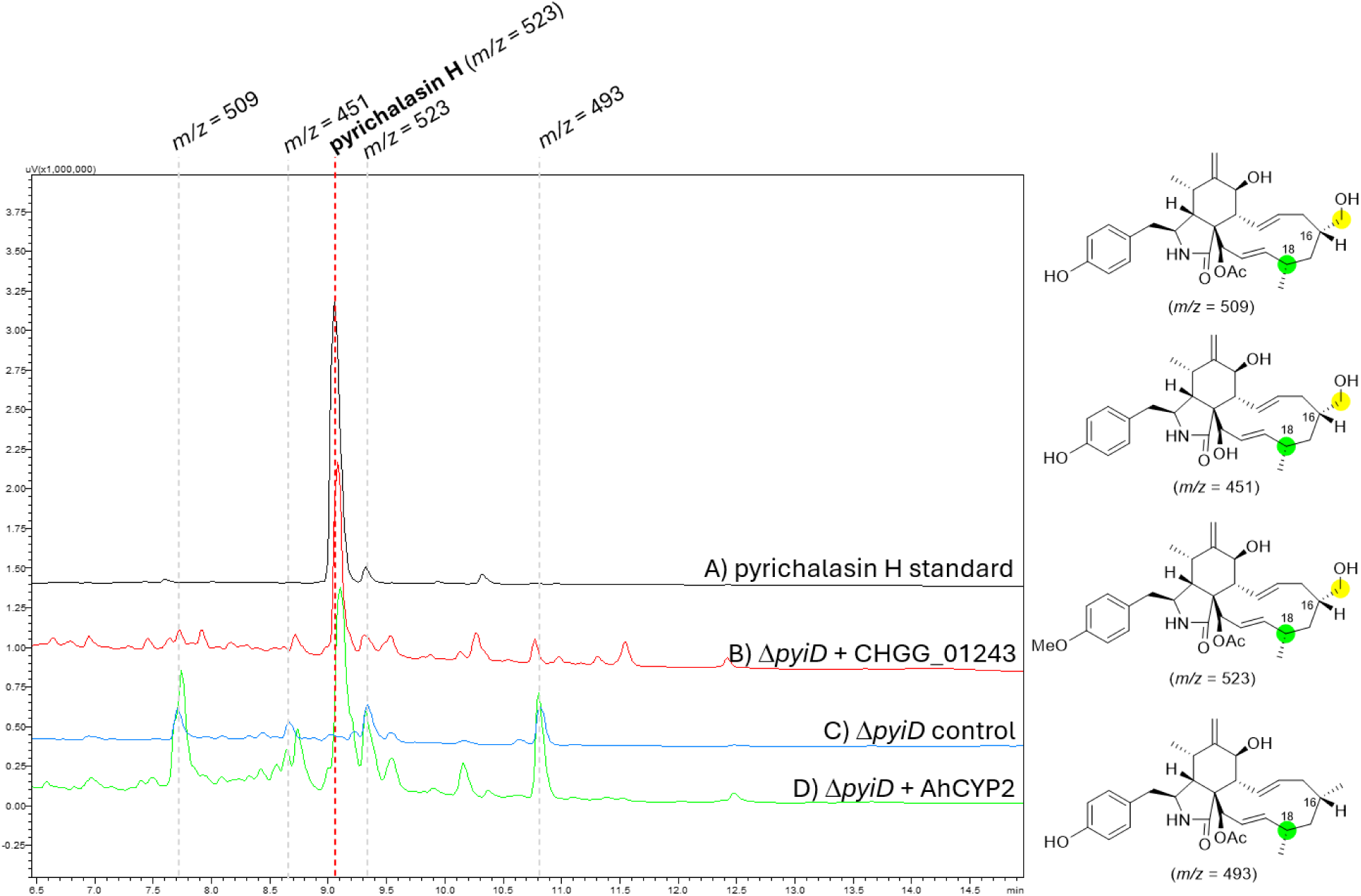
LCMS chromatograms of mycelial extracts prepared from *M. grisea* Δ*pyiD* strains monitored at 190nm. Previously isolated cytochalasans present are indicated by mass with the corresponding structures shown to the right [15]. The C-18 position that PyiD hydroxylates is highlighted in green; oxidations from unknown enzymes encoded outside of the *pyi* BGC are highlighted in yellow.

Next, we selected to heterologously express *ccsD* from *Aspergillus clavatus* NRRL1 in *M. grisea* Δ*p*yi*D.* Compared to pyrichalasin H, cytochalasin E/K has the same sized polyketide-derived macrocycle and methyl groups at the same positions, therefore is more structurally similar than chaetoglobosin A (Figure 5). Despite these structural similarities, none of the resulting transformants exhibited restored pyrichalasin H production nor produced any new metabolites, even when the extended N-terminal α-helix sequence was included (Figures S33-39). RT-PCR on selected transformants confirmed that *ccsD* was transcribed, and while the second intron was correctly spliced, it appears that *M. grisea* had difficulties processing the first intron, resulting in the intron not being spliced, being fully spliced, or partially spliced preventing translation into functional protein (Figures S9-S10).

Therefore, XsCYP2 from *Xylaria* sp. X802 was synthesized from its cDNA sequence [34] and heterologously expressed in *M. grisea* Δ*p*yi*D*. Compared to pyrichalasin H, 19,20-epoxycytochalasin D has the same sized polyketide-derived macrocycle with methyl groups at the same positions (Figure 5). Of the transformants obtained, none restored pyrichalasin H production and the LCMS traces did not reveal obvious new metabolites (Figure S40). RT-PCR confirmed that the gene sequence was transcribed with no errors (Figures S9 and S11), therefore the lack of function of this P450 must be due to another, as yet, unknown reason.

Finally, we turned our attention to the cryptic P450s identified from genome mining. Due to the very close similarity between the BGCs from *A. heteromorphus* CBS 117.55 (*ahe*) and *A. sclerotioniger* CBS 115572 (*asc*); *C. spinosum* CBS 515.97 (*csp*) and *C. sidae* CBS 518.97 (*csi*); and *M. robertsii* ARSEF 23 (*mro*) and *M. brunneum* ARSEF 3297 (*mbr*) respectively (Figure 2, Table S3) we opted to experimentally investigate the predicted macrocycle-associated P450s from one of each pair of BGCs. Although several cytochalasans have been isolated from *A. sclerotioniger*, none include oxidative modifications around the macrocycle (Figure S1B) [27], therefore in case AsCYP2 was non-functional we chose to work with the closely related AhCYP2 from *A. heteromorphus* CBS 117.55 since a putative cytochalasan with oxidative modifications around the macrocycle had been detected by HR-LCMS/MS analysis. Heterologous expression of AhCYP2 into *M. grisea* Δ*pyiD* successfully restored pyrichalasin H production (Figure 6D) in 3 out of 6 transformants. No additional peaks were observed in the chromatogram, indicating that AhCYP2 is not iterative under the conditions tested (Figures S41 – S46). This experiment therefore confirms that AhCYP2 is functional, introns are spliced correctly, and that its designation as a macrocycle-associated P450 is correct.

Heterologous expression of MrCYP1 from *Metarhizium robertsii* ARSEF 23 in *M. grisea* Δ*pyiD* did not restore pyrichalasin H production or lead to any new metabolites being detected in the eight transformants obtained (Figure S47). cDNA was prepared from a transformant which confirmed that MrCYP1 was expressed (Figure S12). MrCYP1 is in the same clade as XsCYP2 and therefore assumed to oxidatively modify cytochalasans derived from C_16_ polyketides, yet, similarly to CcsD and XsCYP2, is also non-functional in this system.

CspCYP1 from *C. spinosum* CBS 515.97 appears related to P450s that act on cytochalasans derived from a C_16_ backbone based on it forming a clade with CYP4, which we previously investigated using this system and did not find to be functional [13]. Similarly, heterologous expression in of CspCYP1 in *M. grisea* Δ*pyiD* did not restore production of pyrichalasin H and no new peaks were observed in the LCMS chromatogram (Figures S48-51). As a control, we also heterologously expressed CspCYP2 in *M. grisea* Δ*pyiG* – a mutant which produces pyrichalasin H analogues lacking a hydroxy group at C-7 – and were able to restore pyrichalasin H production (Figure 7C, Figures S57 - 62), confirming the prediction that CspCYP2 is a cyclohexene-associated P450.

**Figure 7.**
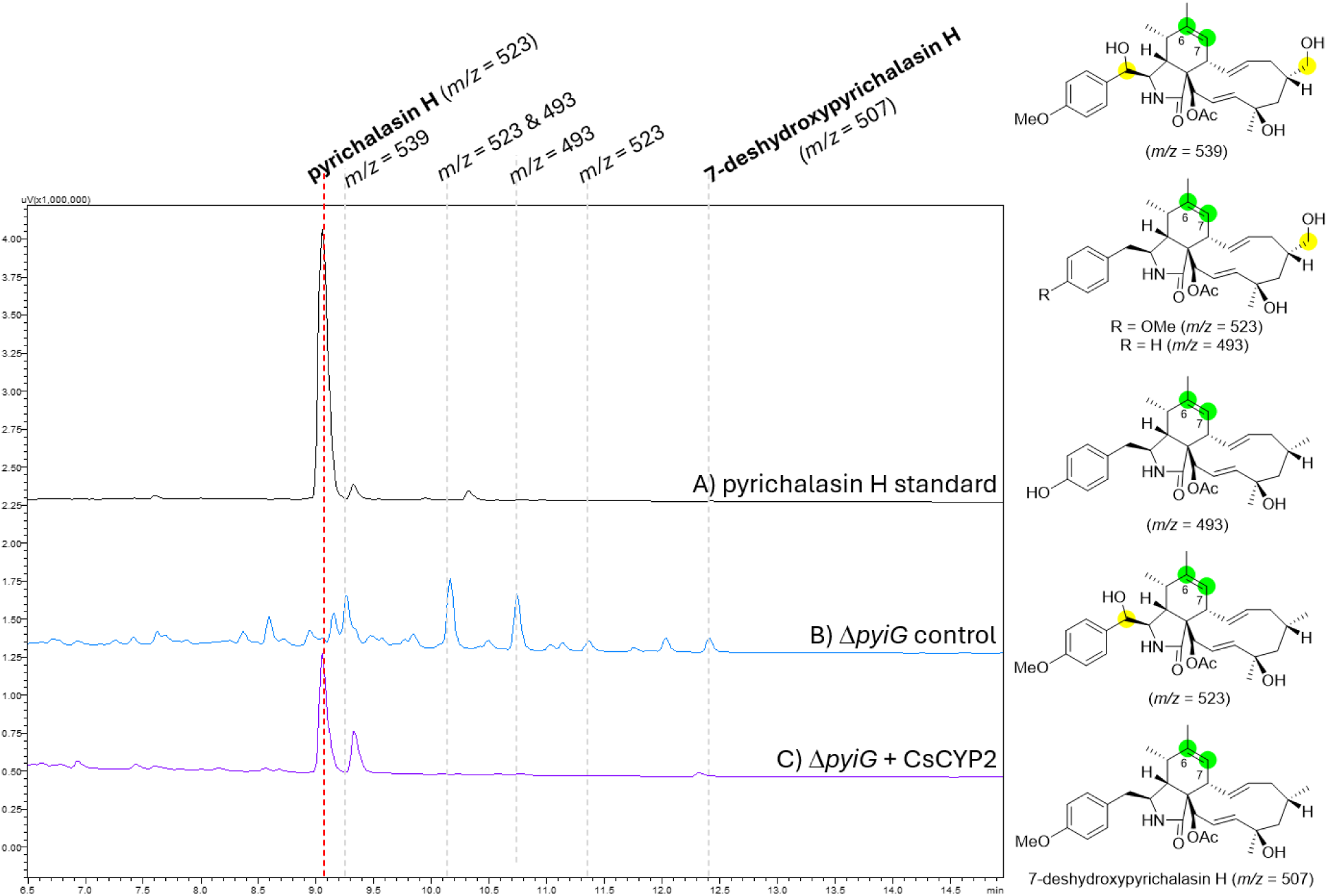
LCMS chromatograms of mycelial extracts prepared from *M. grisea* Δ*pyiG* strains monitored at 190nm. Previously isolated cytochalasans present are indicated by mass with the corresponding structures shown to the right [13]. The C-6/7 positions that PyiG epoxidizes is highlighted in green; oxidations from unknown enzymes outside of the *pyi* BGC are highlighted in yellow. Note: Due to difficulties with synthesizing the genomic CspCYP2 sequence due to repetitive sequences, the putative cDNA sequence was used.

### Biotransformation as an alternative method for functionalizing cytochalasans

Recognizing that combinatorial biosynthesis is quite a laborious procedure, we set out to determine whether biotransformation is possible for exogenously fed cytochalasans. The major intermediate 7-deshydroxypyrichalasin H was purified from *M. grisea* Δ*pyiG* which possesses intact cyclohexene functionality (Figure S63 – S65). 5 mg of 7-deshydroxypyrichalasin H in DMSO was pulse fed to *M. grisea* Δ*pyiD* - which possesses the functional cyclohexene P450 PyiG – from day 4 to 6 of fermentation. After 7 days the culture broth was extracted with EtOAc, concentrated, and analyzed by LCMS, confirming that pyrichalasin H production was restored again (Figure 8; Figures S59 - 60). This result indicates that exogenously added cytochalasans are able to enter the cell and can be modified *in situ* offering a rapid way for selectively functionalizing cytochalasan skeletons.

**Figure 8.**
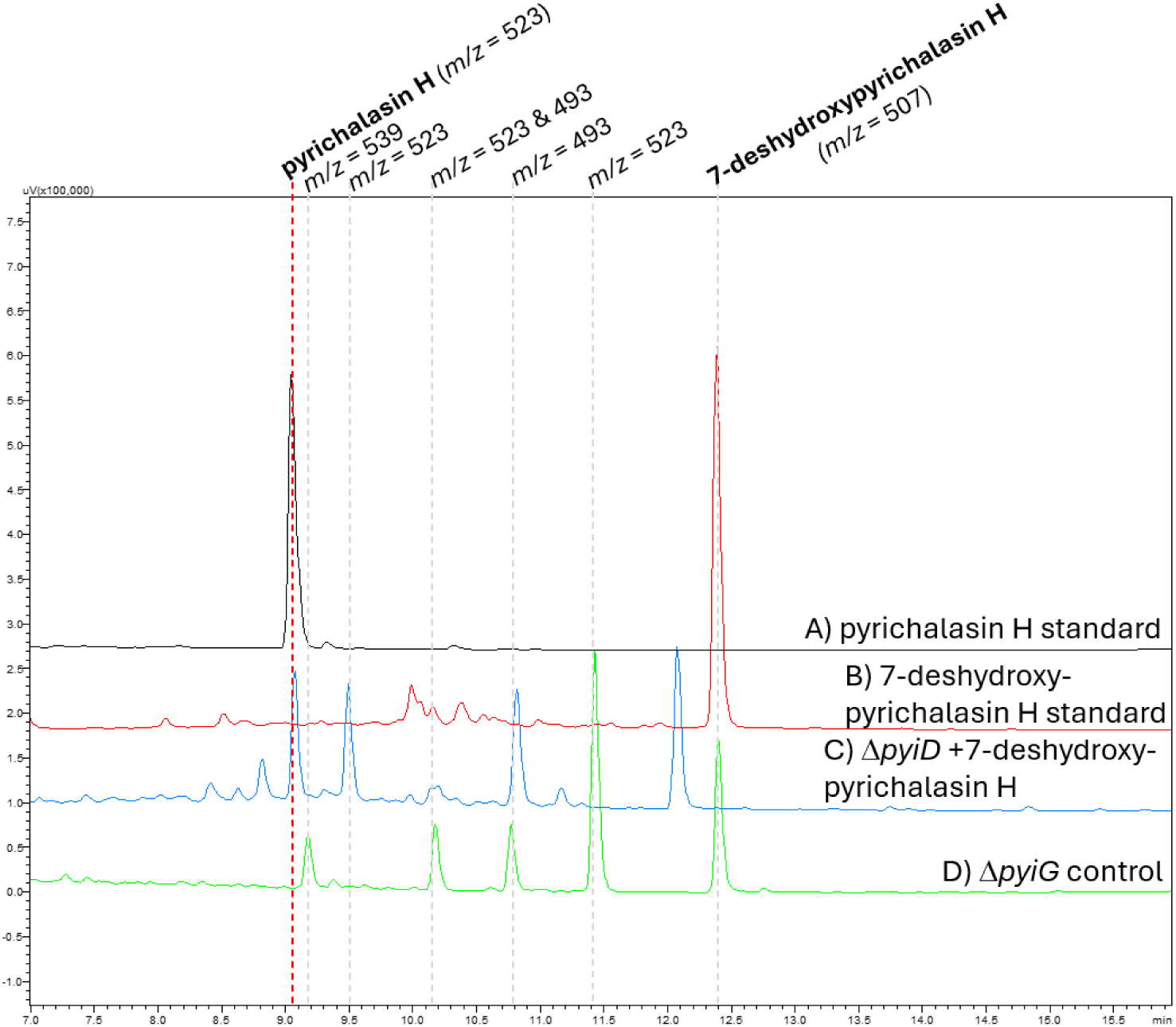
LCMS chromatograms of mycelial extracts prepared from *M. grisea* Δ*pyiD/G* strains monitored at 275nm. Previously isolated cytochalasans present are indicated by mass with the corresponding structures shown in Figures 5 and 6.

## Discussion

Biosynthetic investigations of different cytochalasan enzymes have revealed a high level of substrate promiscuity, [7,13,15] which may explain the large number of cytochalasan molecules isolated from various fungal sources demonstrating only slight structural differences [1,2,5]. Expanding on our prior work, where cytochalasan P450s from either closely related species or those which act upon nearly structurally identical molecules were investigated *via* combinatorial biosynthesis [13], here we focused on macrocycle P450s that are either known, or suspected to be, iterative and a broader array of cryptic P450s, to understand the scope of these P450s as biocatalysts.

Genome mining enabled us to identify three pairs of closely related cytochalasan BGCs from *Aspergillus*, *Colletotrichum*, and *Metarhizium* sp. (Figure 2). Surprisingly, a TRX-like gene was identified in BGCs that also encode a BVMO, consistent with the TRX-like genes also observed in the cytochalasan BGCs from *A. clavatus* NRRL1 and *Peroneutypa* sp. M16 [3,20]. Performing cblaster and clinker analyses, identified a TRX-like gene in over a dozen highly diverse fungal species (Figure S3). Since the TRX-like gene is relatively small (< 750bp) we manually searched the pyrichalasin H, chaetoglobosin A, and 19,20-epoxycytochalasin D BGCs in case it had been overlooked by gene calling programs, but could not identify a similar gene sequence. Attempts to knock-out the TRX-like gene in *A. clavatus* NRRL1 were unsuccessful, most likely due to the competing non-homologous end-joining (NHEJ) pathway which is predominant in fungi [35]. Although genes have been knocked out in *A. clavatus* NRRL1, also using the *trpC* promoter and basta selection marker, the host strain had the NHEJ protein XXX already disrupted [14], likely leading to much higher knock-out efficiency. Since the TRX-like gene is only observed in cytochalasan BGCs that also encode BVMOs we speculate that the TRX-like enzyme may be involved in scavenging peroxide radicals released from the BVMO in a process known as uncoupling [36], however we plan to confirm this in a follow-up study.

Since no cytochalasans have been reported from any of the strains except for *A. sclerotioniger* CBS 115572 (Figure S1), we grew *A. heteromorphus* CBS 117.55 on a media panel and screened for expression of the PKS-NRPS gene. In four of five media tested, the PKS-NRPS gene was expressed, whereas in all five media the TRX-like was expressed (Figure S8). Examination of the organic extracts’ HRMS data with MZMine and MS-DIAL revealed production of a putative cytochalasan with chemical formula C_28_H_33_NO_7_ in four out of five media also, albeit in minor amounts (Figure 3). Combined with the *ahe* / *asc* / *ccs* BGC comparison and phylogenic analysis, we propose that this cytochalasan derives from a C_18_ polyketide lacking pendant methyl groups; analysis of the MS^2^ fragmentation supports our putative assignment (Figure S21).

Phylogenetic analysis clearly differentiates between P450s that act on the macrocycle *vs*. the cyclohexene, with more divergence observed for macrocycle-associated P450s (Figure 5). This is not necessarily surprising as we recently showed that epoxide opening was a non-enzymatic process [20] and therefore it is likely that all cyclohexene associated P450s introduce epoxide functionality, which rearrange under acidic or basic conditions to give the observed alcohol and alkene functional groups. The clades formed by the macrocycle-associated P450s appear to reflect structural features of the final cytochalasan molecule with respect to the type of amino acid incorporated, polyketide chain length, and number of methyl groups (Figure 5).

Based on the small number of experimentally characterized cytochalasan P450s, it is not yet possible to determine which P450s are iterative and those which are not, based on protein sequence alone. Therefore, CHGG_01243, an experimentally validated iterative P450 [7], and CcsD and XsCYP2, two P450s that are suspected to be iterative based on the oxidative modifications at the adjacent C-17 and C-18 positions (Figure 5), and absence of a third P450 enzyme encoded within the BGC, were heterologously expressed in *M. grisea ΔpyiD*. While CHGG_01243 could restore pyrichalasin H production, no additional cytochalasans were identified, indicating that this known iterative P450 does not act iteratively on a non-native substrate with a smaller sized macrocycle (Table 1). Neither CcsD nor XsCYP2 restored pyrichalasin H production or caused a change in the metabolite profile of the *M. grisea ΔpyiD* extracts (Table 1). While intron processing issues, the truncated N-terminal α-helix, and the structural differences *e.g.* the lack of carbonate functional group in pyrichalasin H intermediates may explain why CcsD is non-functional, the lack of activity by XsCYP2 is more challenging to explain since it was cloned from a cDNA sequence, the N-terminal sequence is 70% identical to the native PyiD N-terminal sequence, and the cytochalasan backbone and functional groups are almost identical to pyrichalasin H (Table 1).

**Table 1:**
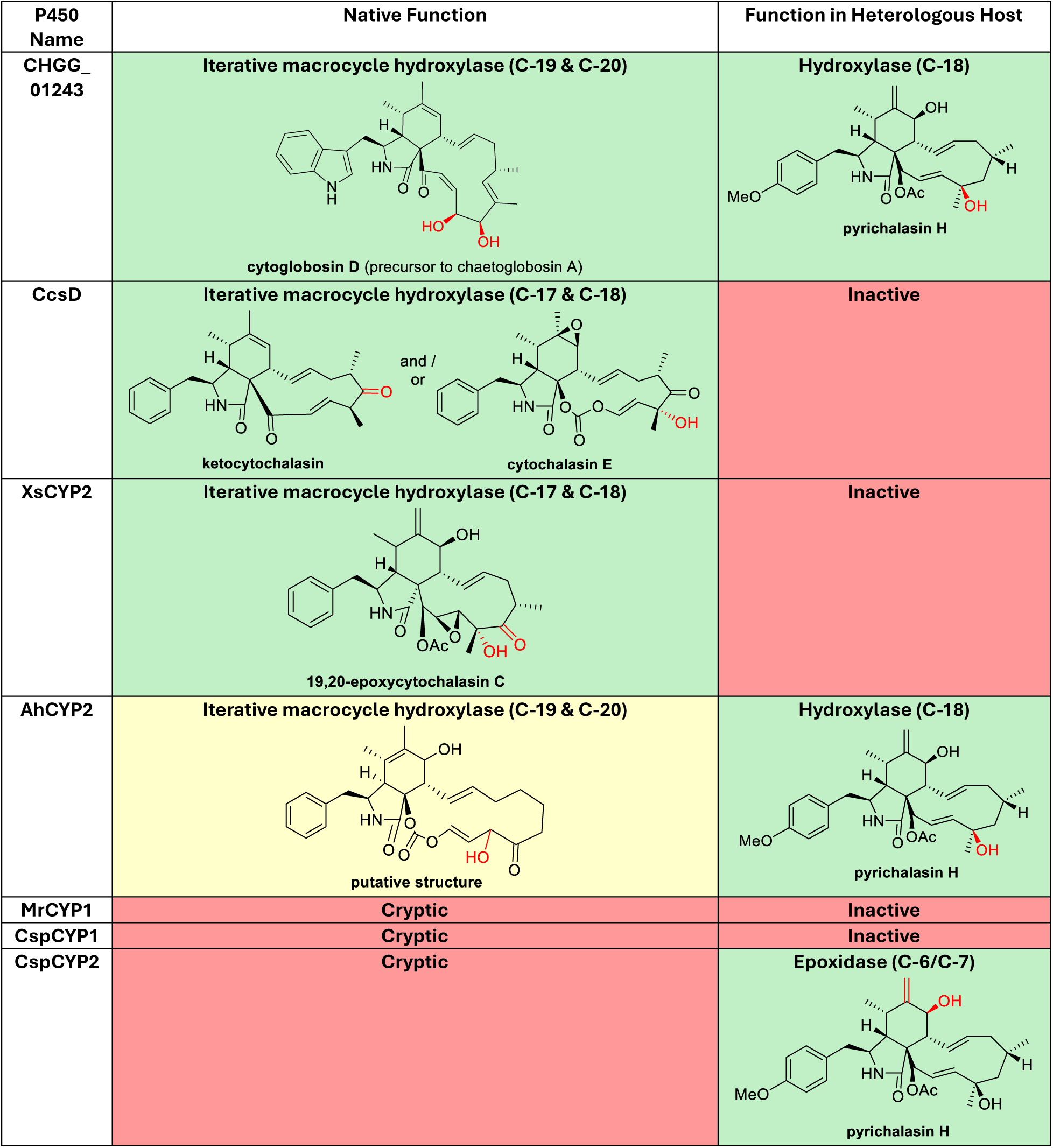
Summary of heterologous expression results and the deduced functions of the cryptic P450s. The oxygens introduced by the P450s and subsequent modifications are highlighted in red.

| P450 Name | Native Function | Function in Heterologous Host |
| --- | --- | --- |
| CHGG_01243 | **Iterative macrocycle hydroxylase (C-19 & C-20)**<br>cytoglobosin D (precursor to chaetoglobosin A) | **Hydroxylase (C-18)**<br>pyrichalasin H |
| CcsD | **Iterative macrocycle hydroxylase (C-17 & C-18)**<br>ketocytochalasin and / or cytochalasin E | Inactive |
| XsCYP2 | **Iterative macrocycle hydroxylase (C-17 & C-18)**<br>19,20-epoxycytochalasin C | Inactive |
| AhCYP2 | **Iterative macrocycle hydroxylase (C-19 & C-20)**<br>putative structure | **Hydroxylase (C-18)**<br>pyrichalasin H |
| MrCYP1 | Cryptic | Inactive |
| CspCYP1 | Cryptic | Inactive |
| CspCYP2 | Cryptic | **Epoxidase (C-6/C-7)**<br>pyrichalasin H |

A subtle structural difference between pyrichalasin H and 19,20-epoxycytochalasin D, is the stereochemistry of the methyl groups around the macrocycle (Table 1). For example, the methyl groups at C-16 and C-18 in pyrichalasin H have a *cis* relationship; in contrast, the C-16 and C-18 positions in 18,19-epoxycytochalasin D - and cytochalasin E - have a *trans* relationship, with the C-18 methyl group having the inverse stereochemistry to the C-18 position in pyrichalasin H. In contrast, chaetoglobosin A and the putative cytochalasin from *A. heteromorphus* CBS 117.55, lack methyl groups where the macrocycle P450 acts, and the corresponding P450s were shown to be active in our platform. Therefore, the stereochemistry of methyl groups combined with the flexible conformation of the macrocycle (Figure S14) may restrict which P450s can be used successfully with non-native substrates.

The mechanism of hydrocarbon hydroxylation performed by P450s typically proceeds with a retention of stereochemistry [37], indicating that the methyl groups’ stereochemistry around the macrocycle must be established during biosynthesis of the polyketide chain by the PKS-NRPS enzyme. Oikawa and colleagues have established a method for predicting the absolute stereochemistry of methyl groups in fungal polyketides arising from highly reduced polyketide synthases (HRPKS) in predominantly linear polyketides [38]. Due to cytochalasans having a large number of post-PKS modifications obscuring these stereochemical rules, we instead analyzed 385 cytochalasan structures recently reported (Sheet S1) [2]. 57% of cytochalasans derive from a C_16_ polyketide chain (octaketides), 40% derive from a C_18_ polyketide chain (nonaketides), with the remainder being derived from C_12_, C_14_, or C_20_ polyketide chains. The relative stereochemistry of the methyl groups at the C-16 and C-18 positions are shown in Figure 9, demonstrating high variability, which is also reflected in our phylogenetic analysis of a much smaller subset of P450s (Figure 5).

**Figure 9:**
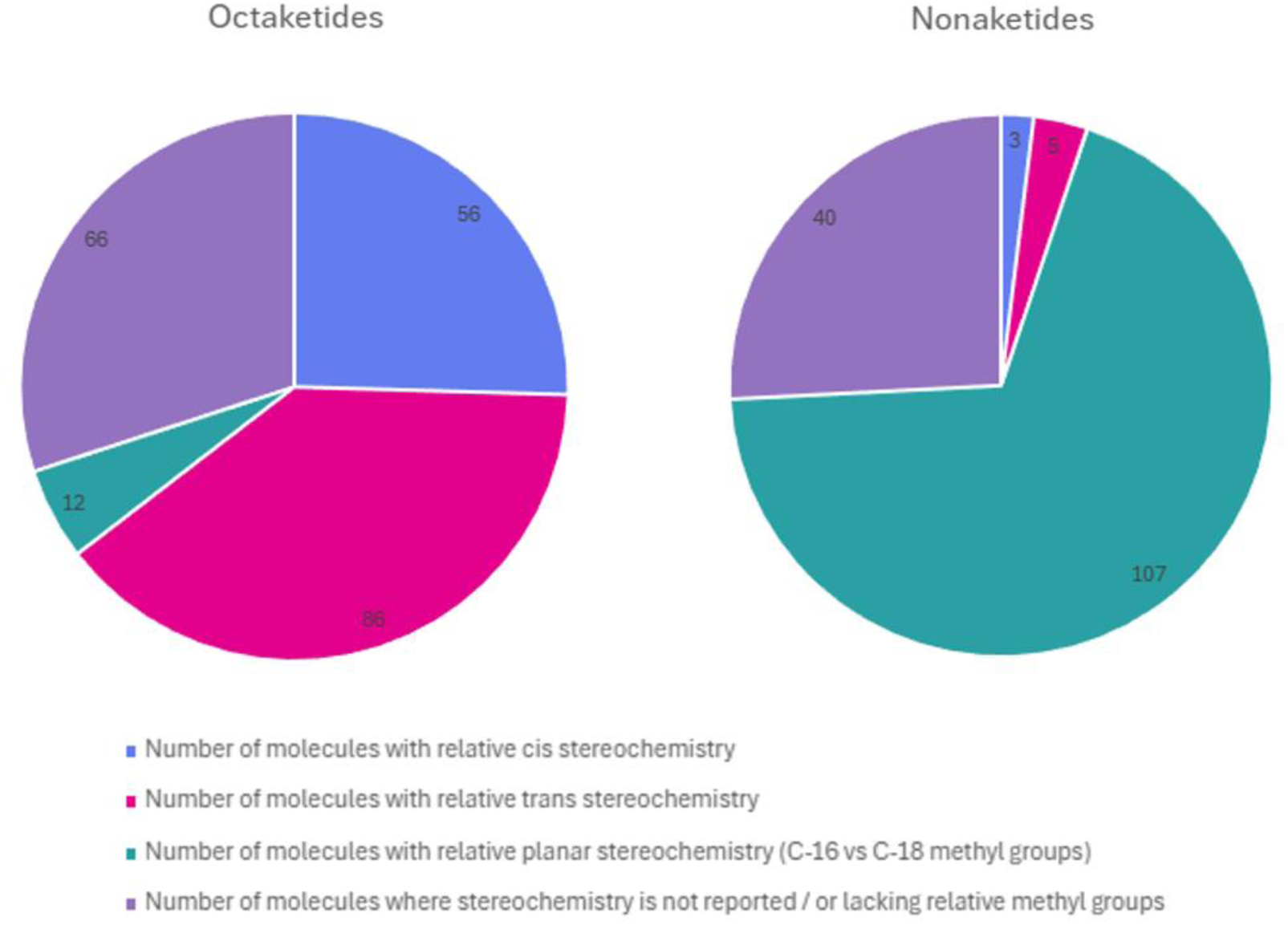
Structural analysis of recently discovered cytochalasans categorized according to the number of carbons in the polyketide chain and the number and relative stereochemistry of methyl groups around the macrocycle.

Since no additional cytochalasans were identified, MrCYP1 and CspCYP1 are presumed inactive on the available pyrichalasin H intermediates as RT-PCR data confirms transcription of the genes (Figure S9). MrCYP1 forms a clade with XsCYP2 (Figure 5) indicating that the stereochemistry of the C-18 methyl group may prevent hydrogen abstraction by the P450 enzyme. CspCYP1 forms a clade with CYP4 (Figure 5) which is a P450 presumed to oxidatively modify the cryptic SYN2/RAP2 cytochalasan produced by *Pyricularia oryzae* Guy11 [39]. When we previously investigated the function of CYP4 it appeared to be inactive in our platform. [13] Larsen and co-workers had heterologously expressed *SYN2* and *RAP2* in *Aspergillus nidulans* resulting in the decalin product niduporthin as the hydrolase and Diels-Alderase enzymes were not included [40]. Niduporthin derives from a C_16_ polyketide with just one pendant methyl group fused to tryptophan [40]. Although the position of this methyl group corresponds to the C-18 position if the molecule was drawn as cytochalasan, it is not possible to determine its absolute or relative stereochemistry.

As an alternative to combinatorial biosynthesis, which relies on the correct annotation of fungal genes, introns from non-native sequences to be correctly spliced, and laborious rounds of screening of transformants, we investigated whether exogenously added cytochalasans could be biotransformed. By purifying the major cytochalasan intermediate 7-deshydroxypyrichalasin H from *M. grisea* Δ*pyiG* and feeding it to *M. grisea* Δ*pyiD* we could detect restoration of pyrichalasin H biosynthesis, indicating that exogenously added cytochalasans can penetrate the cell wall and be biotransformed by native enzymes.

In conclusion, we demonstrate that while cytochalasan P450s possess intrinsic substrate promiscuity, this is restricted by the stereochemistry of existing functional groups decorating the macrocycle at the oxidation site. The size of the macrocycle is of lesser significance, provided that the atomistic composition of the macrocycle is not hindered. These observations may enable more efficient production of novel and unnatural cytochalasans with enhanced bioactivities by correctly pairing substrates with P450 enzymes. Finally, our work reveals significant variation in the number, positions, and stereochemistry of methyl groups introduced into cytochalasan backbones by the PKS-NRPS enzymes, which may also be a major determinant in the biological activities observed.

## Supporting information

Supplementary Sheet 1

## List of abbreviations

A: adenylation domain
ACP: acyl carrier protein domain
AcT: acyl transferase
AT: acyl transferase domain
BVMO: Baeyer-Villiger monooxygenase
C: condensation domain
CYP: cytochrome P450 monooxygenase
DA: Diels-Alderase
DH: dehydratase domain
ER_0_: inactive enoyl reductase domain
EtOAc: ethyl acetate
HYD: hydrolase
KR: ketoreductase domain
KS: ketosynthase domain
LCMS: liquid chromatography mass spectrometry
MFS: major facilitator superfamily transporter
MT: methyl transferase domain
NMR: nuclear magnetic resonance
OXR: oxidoreductase
P450: cytochrome P450 monooxygenase
PCP: peptidyl carrier domain
PKS-NRPS: polyketide synthase / non-ribosomal peptide synthetase
RT-PCR: reverse transcription polymerase chain reaction
Trans-ER: trans-enoyl reductase domain
TRX: thioredoxin

## Declarations

### Ethics approval and consent to participate

Not applicable

### Consent for publication

Not applicable

### Availability of data and materials

All data generated or analysed during this study are included in this published article [and its supplementary information files].

### Authors’ contributions

LL constructed plasmids, performed transformations, extractions, LCMS analysis, purifications and NMR analysis.

TA performed HR-LCMS/MS, MZMine, and MS-DIAL analysis.

JG purified pyrichalasin H from *M. grisea* NI980.

SJ prepared RNA and performed RT-PCR.

AP performed extractions and LCMS analysis.

MRA performed NMR analysis and contributed to funding acquisition (FAPESP).

CB contributed to study design and funding acquisition (DFG).

ES was responsible for overall study design, supervision of LL, TA, JG, SJ, AP and MRA, funding acquisition (NSF), and performed genome mining / bioinformatics analysis, plasmid construction, transformation, extractions, and LCMS analysis.

## Acknowledgements

We would like to thank Dr. Katherine Williams from UWE Bristol, UK for the pE-YA and pTYGSxxx fungal / yeast / *E. coli* cloning and expression plasmids, as well as Prof. Dr. Russell J. Cox from LUH, Germany for the *M. grisea* NI980 strains. We also want to recognize Dr. Hongjun Pan for the acquisition of NMR spectral data and Jean Christophe Cocuron from the Bioanalytical Facilities (BAF) for acquiring HRMS data and useful discussions.

