## Supplementary Information for "Investigating the versatility of cytochalasan cytochrome P450 monooxygenases using combinatorial biosynthesis reveals stereochemical restrictions"

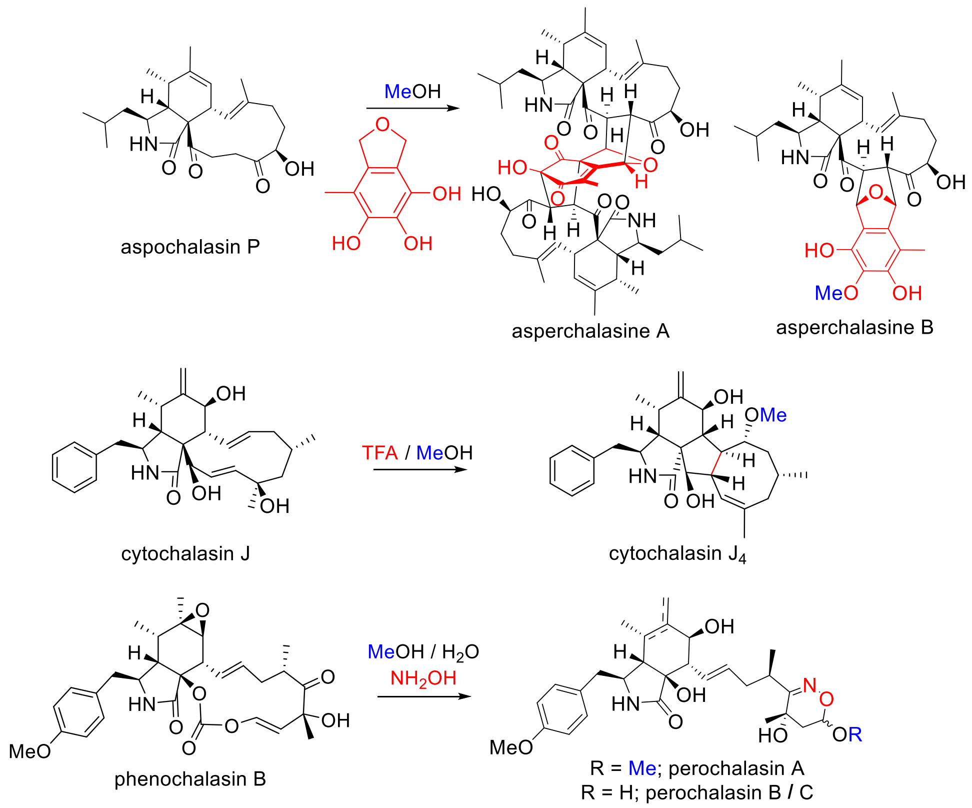


**Scheme S1**: Examples of non-enzymatic modifications during cytochalasan biosynthesis, *via* interactions with solvents *e.g.* MeOH; small molecules *e.g.* TFA and $NH_2OH$; and natural products also synthesized by the fungal strain *e.g.* epicoccine. [1– 3]

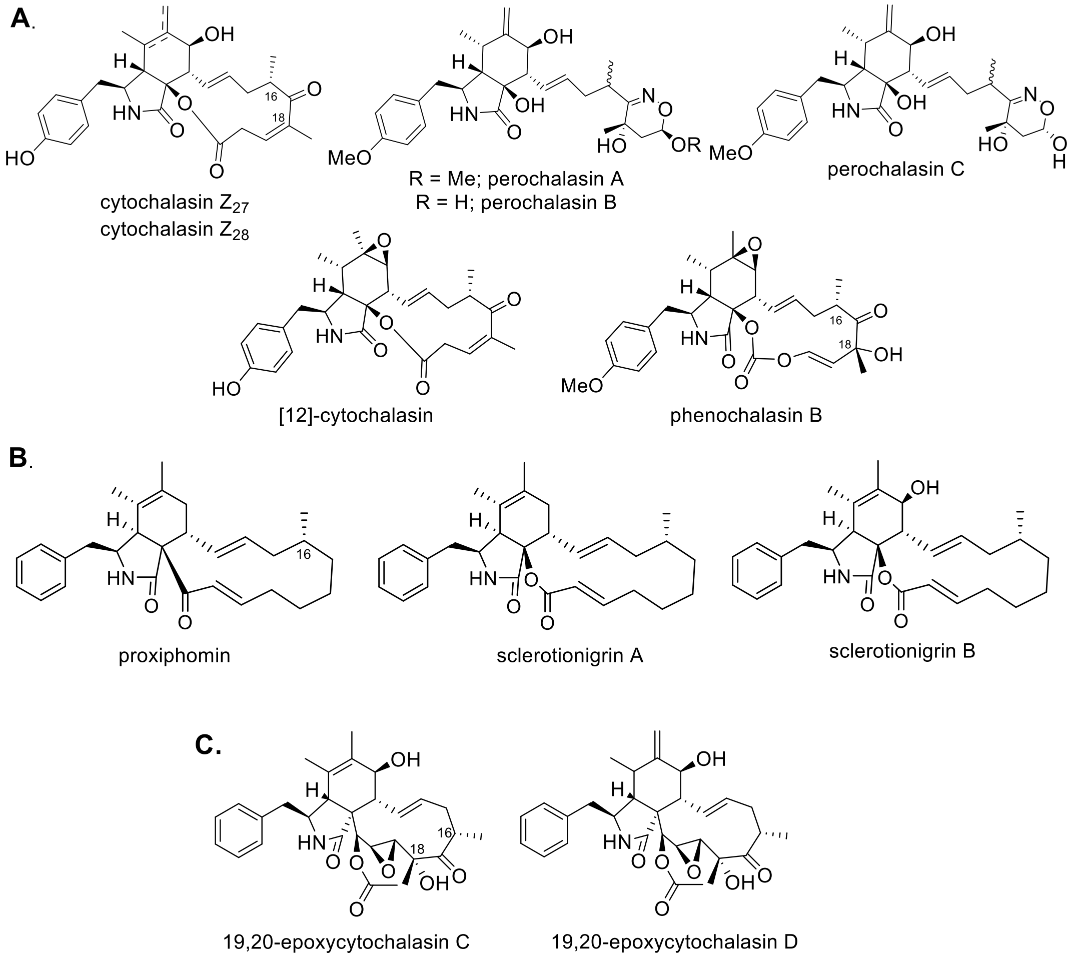


**Figure S1**: Structures of cytochalasans bioinformatically linked to a BGC, relevant to this study, grouped according to which fungal strain they were isolated from. A) *Peroneutypa* sp. M16 [3]; B) *Aspergillus sclerotioniger* CBS 115572 [4]; C) *Pseudoxylaria* sp. X802 [5].

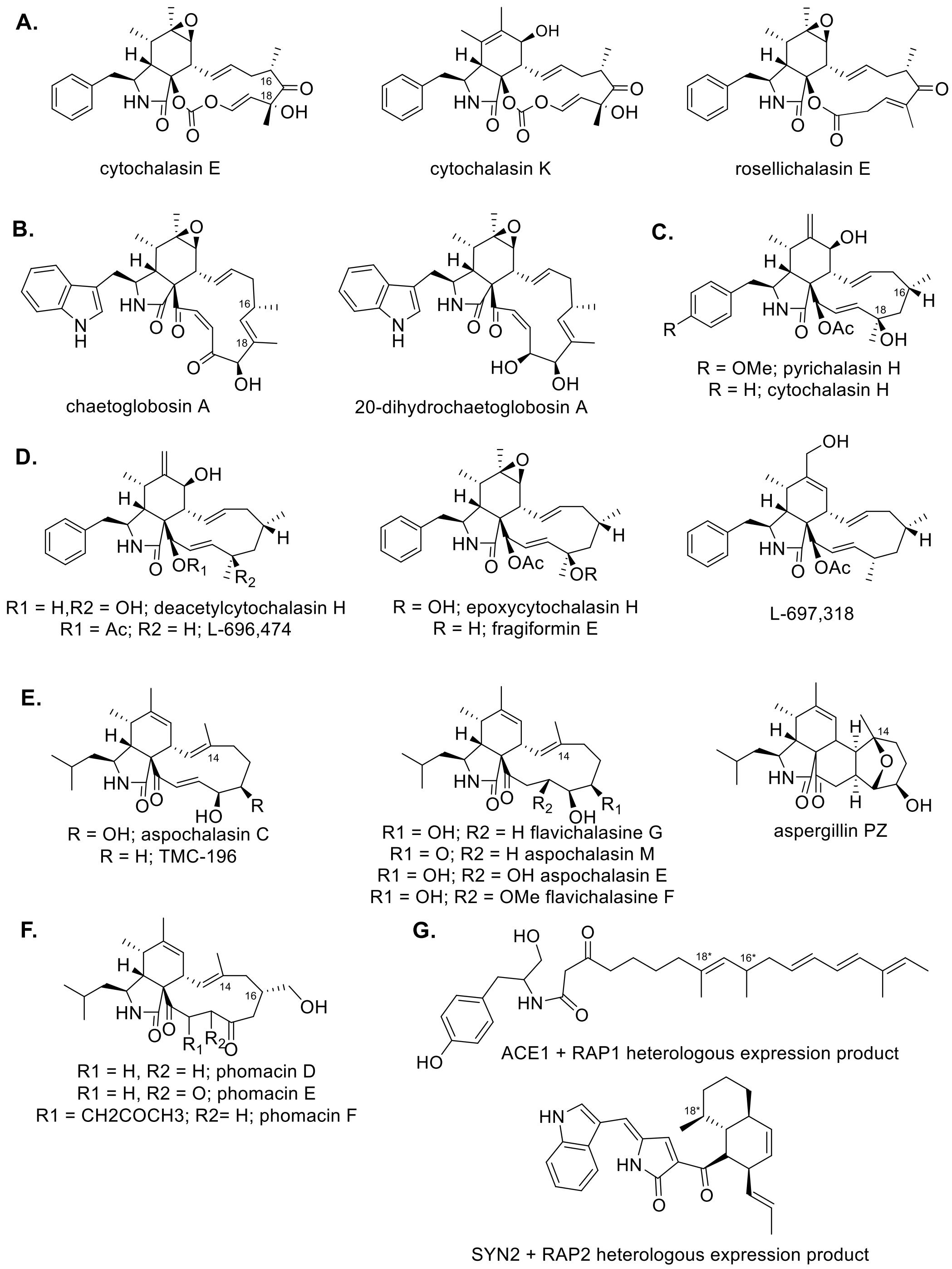


**Figure S2**: Structures of cytochalasans experimentally linked to a BGC, relevant to this study, grouped according to which fungal strain they were isolated from. A) *Aspergillus clavatus* NRRL1 [6]; B) *Chaetomium globosum* CBS 148.51 [7]; C) *Magnaporthe grisea* NI980 [8]; D) *Hypoxylon fragiforme* MUCL 51264 [9]; E) *Aspergillus flavipes* CNL-338 [10]; F) *Parastogonospora nodorum* [11]; G) *Pyricularia oryzae* Guy 11 ACE1/SYN2 heterologous expression products [12,13].

1. **General experimental procedures**

**1.1 Reagents**

Analytical grade chemicals, LCMS grade solvents, and reagents were purchased from Sigma Alrich and Thermo Fisher Scientific. General molecular biology procedures were performed as standard; molecular biology kits (QIAGEN, Invitrogen, New England Biolabs, Sigma Aldrich, Zymo Research) were used according to the manufacturer's protocols unless stated otherwise. PCR was performed using Q5 polymerase and cDNA synthesis used the LunaScript RT SuperMix Kit (NEB). Restriction endonucleases were purchased from NEB or Thermofisher Scientific. The chemical standard of cytochalasin E was purchased from Cayman Chemicals.

Oligonucleotides used in this work (Table S1) were purchased from Integrated DNA Technologies (IDT). Synthetic genes and gene fragments were purchased from Twist Bioscience or BioCat GmbH. PCR products were sequenced by Eurofins Genomics or Plasmidsaurus; plasmids generated in this study were sequenced by Plasmidsaurus.

| **Primers used in this study** | **Sequence** | **Description** |
|---|---|---|
| LL X802 F | TCGAGCCTTTTCTGGTGCTGCG | For confirming pTYGS bar + XsCYP2 |
| LL X802 F | AAGGACTCGCGGAGGAAGCTGT | |
| CcsD F | ATGGTGAACGAGATAAGCCC | For confirming expression of CcsD |
| CcsD R | TTAGTTTCGAGACTTGAACA | |
| X802 CYP2 F | ATGTTGTCAAGCATCCAAAC | For confirming expression of XsCYP2 |
| X802 CYP2 R | CTAACATCTGGCCCTAATAC | |
| MrP450-1 F | GTCTGAATGCTCGACAACAT | For confirming expression of MrCYP1 |
| MrP450-1 R | CTAAGATCTGGACCTAATGC | |
| CsP450-1 F | ATGTCATTCTCCATCGAATC | For confirming expression of CspCYP1 |
| CsP450-1 R | TCACTTGCCAGCCCTCGACT | |
| Ah PKS-NRPS F | TGCAGAGTCAGATCCCACATG | For confirming expression of the PKS-NRPS gene present in the *ahe* putative cytochalasan BGC |
| Ah PKS-NRPS R | TTGATGCCAGTCTCCCGAATG | |
| Ahet TRX F | ATGACCCTCATCAAGATCTC | For confirming expression of the TRX-like gene in the *ahe* putative cytochalasan BGC |
| Ahet TRX R | CTAGCATGTATTCCCCAAGG | |
| 5TRX-Hyg F | TTGCGAACCTGCTTGTCAAC | For cloning *ccsX* TRX KO cassette from synthetic fragments |
| 5TRX-Hyg R | GGCGACCTCGTATTGGGAAT | |
| Hyg-3TRX F | TCAGCGAGAGCCTGACCTAT | |
| Hyg-3TRX R | CACAACCGAGCTATGCATCC | |

**Table S1:** Primers used in this study

#### 1.2 Bioinformatics methods

Experimentally characterized cytochalasan BGCs were identified using MiBIG or NCBI (Table S2). The PyiF protein sequence was used with BLAST to identify publicly available genomes containing cytochalasan BGCs. Additional genes required for cytochalasan biosynthesis were identified using contig walking *via* NCBI Gene (Table S3). For *Pseudoxylaria* sp. X802 the whole genome sequence (WGS; JAJFDH000000000) was uploaded to fungiSMASH to identify the putative cytochalasan BGC (Table S4). Once a putative cytochalasan BGC was identified, the region was downloaded and reannotated using FGENESH to further refine gene boundaries, introns, and resulting protein sequence. [14] Sequence alignments and phylogenetic trees were generated using Geneious™ keeping the default settings. Coding sequences of P450s (Table S5) were investigated using BLAST, multiple sequence alignments (Figure S4), and AlphaFold.

The BMVO (CcsB) and TRX (CcsX) sequences from the *A. clavatus* NRRL1 *ccs* BGC were uploaded to cblaster to identify other fungal BGCs encoding these proteins. The resulting BGCs were visualized using cblaster using the default parameters (Figure S5). [15]

| **Fungal Strain** | **Cytochalasan biosynthetic proteins identified** | **Corresponding NCBI / MiBIG accession numbers** |
|---|---|---|
| *Magnaporthe grisea* NI980 | PyiA<br>PyiE<br>PyiT<br>PyiG<br>PyiC<br>**PyiF**<br>PyiD<br>PyiR<br>PyiB<br>PyiH<br>PyiS | BGC0001881 |
| *Aspergillus clavatus* NRRL 1 | ACLA_078610<br>ACLA_078620<br>ACLA_078630<br>ACLA_078640<br>ACLA_078650<br>ACLA_078660<br>ACLA_078670<br>ACLA_078680<br>**ACLA_078690**<br>ACLA_078700<br>ACLA_078710<br>ACLA_078720<br>ACLA_078730 | BGC0000983 |
| *Chaetomium globosum* CBS 148.51 | TF<br>Transposase<br>PKS-NRPS<br>Trans-ER<br>**DA** | CHGG_01237<br>CHGG_01238<br>CHGG_01239<br>CHGG_01240<br>CHGG_01241 |

| | | |
|---|---|---|
| | Fused P450/OXR<br>P450<br>Unknown<br>OXR<br>HYD<br>Deleted record<br>MFS | CHGG_01242<br>CHGG_01243<br>CHGG_01244<br>CHGG_01245<br>CHGG_01246<br>CHGG_01247<br>CHGG_01248 |
| *Aspergillus flavipes* CNL-338 | FfsC<br>FfsR<br>FfsD<br>FfsA<br>FfsH<br>**FfsI**<br>QOG08947.1<br>FfsF<br>FfsJ<br>Ffs-ORF1<br>Ffs_ORF2 | BGC0002204 |
| *Parastagonospora nodorum* SN15 | SNOG_00306<br>SNOG_00307<br>SNOG_00308<br>SNOG_00309<br>**SNOG_00310**<br>SNOG_00311<br>SNOG_00312<br>SNOG_00313<br>SNOG_00314 | BGC0002205 |
| *Pyricularia oryzae* 70-15 | PKS-NRPS<br>Trans-ER<br>HYD<br>OXR<br>Hypothetical<br>P450<br>TF<br>OXR<br>P450<br>MFS<br>**DA**<br>Trans-ER<br>P450<br>P450<br>Hypothetical<br>PKS-NRPS<br>OMeT | MGG_12447<br>MGG_08391<br>MGG_08390<br>MGG_08389<br>MGG_15906<br>MGG_08387<br>MGG_08386<br>MGG_15927<br>MGG_15928<br>MGG_08384<br>MGG_08381<br>MGG_08380<br>MGG_08379<br>MGG_08378<br>MGG_15929<br>MGG_15097<br>MGG_08377 |
| *Hypoxylon fragiforme* MUCL 51264 | HffH<br>HffB<br>HffR<br>HffE<br>HffT<br>HffS<br>HffD<br>**HffF** | QFX78098.1<br>QFX78099.1<br>QFX78100.1<br>QFX78101.1<br>QFX78102.1<br>QFX78103.1<br>QFX78104.1<br>QFX78105.1 |

| | | |
|---|---|---|
| | HffC<br>HffG | QFX78106.1<br>QFX78107.1 |
| *Peroneutypa* sp. M16 | PeroB<br>PeroT<br>PeroR<br>PeroA<br>PeroX<br>PeroG<br>PeroC<br>**PeroF**<br>PeroE<br>PeroD<br>PeroS | XFF06364.1<br>XFF06365.1<br>XFF06366.1<br>XFF06361.1<br>XFF06360.1<br>XFF06362.1<br>XFF06363.1<br>XFF06367.1<br>XFF06368.1<br>XFF06369.1<br>XFF06370.1 |

**Table S2**: Experimentally validated and putative cytochalasan BGCs identified by genome mining. The core cytochalasan genes are highlighted in red and the DA query protein is also bolded. The MiBIG or NCBI accession numbers are provided for each BGC / individual protein respectively.

| **Deduced function** | ***pyi* cluster homologs** | ***asc* cluster proteins (% id*)** | ***ahe* cluster proteins (% id*)** | ***csp* cluster proteins (% id*)** | ***csi* cluster proteins (% id*)** | ***mbr* cluster proteins (% id*)** | ***mro* cluster proteins (% id*)** |
|---|---|---|---|---|---|---|---|
| PKS-NRPS | PyiS (QCS37521.1) | XP_025464061.1 (46%) | XP_025399161.1 (45%) | TDZ28122.1 (53%) | TEA13957.1 (53%) | KID72760.1 (69%) | XP_007816617.1 (69%) |
| *Trans*-ER | PyiC (QCS37515.1) | XP_025464058.1 (46%) | XP_025399158.1 (45%) | TDZ28142.1 (63%) | TEA13956.1 (66%) | KID72751.1 (71%) | XP_007816628.1 (73%) |
| HYD | PyiE (QCS37512.1) | XP_025464062.1 (51%) | XP_025399162.1 (47%) | TDZ28131.1 (61%) | TEA13959.1 (61%) | KID72749.1 (64%) | XP_007816630.1 (64%) |
| DA | PyiF (QCS37516.1) | XP_025464059.1 (48%)$ | XP_025399159.1 (48%) | TDZ28108.1 (61%) | TEA13960.1 (62%) | KID72750.1 (79%) | XP_007816629.1 (74 %) |
| OMeT | PyiA (QCS37511.1) | N/A | N/A | TDZ28140.1 (46%) | TEA13952.1 (48%) | N/A | N/A |
| P450m | PyiD (QCS37517.1) | XP_025464060.1 (43% | XP_025399160.1 (53%) | TDZ28124.1 (51%) | TEA13954.1 (51%) | KID72759.1 (69%) | XP_007816618.2 (68 %) |
| P450c | PyiG (QCS37514.1) | XP_025464057.1 (37%) | XP_025399157.1 (38% - fused to OXR) | TDZ28141.1 (48% | TEA13958.1 (48%) | KID72752.1 (53%) | XP_007816627.1 (69 %) |
| NAD(P)H-OXR | PyiH (QCS37520.1) | XP_025464055.1 (N/A) | N/A | N/A | N/A | KID72761.1 (68%) | N/A |
| OAcT | PyiB (QCS37519.1) | N/A | N/A | N/A | N/A | N/A | N/A |
| MFS | PyiT (QCS37513.1) | N/A | N/A | TDZ28132.1 (52%) | TEA13965.1 (52%) | N/A | N/A |
| TF | PyiR (QCS37518.1) | XP_025464056.1 (26%) | N/A | TDZ28137.1 (36%) | TEA13955.1 (35%) | KID72762.1 (29%) | N/A |
| TF2 | N/A | N/A | N/A | TDZ28123.1 (N/A) | N/A | KID72758.1 (N/A) | XP_007816622.2 (N/A) |
| FAD-OXR | N/A | XP_025464054.1 (N/A) | XP_025399157.1 (fused to P450c) | N/A | N/A | N/A | N/A |
| BVMO | N/A | XP_025464063.1 (60%^) | XP_025399163.1 (57%^) | N/A | N/A | N/A | N/A |
| TRX | N/A | XP_025464053.1 (35%^) | XP_025399156.1 (34% ^) | N/A | N/A | N/A | N/A |

**Table S3**: Comparative analysis of putative cytochalasan BGCs identified by genome mining. Abbreviations: PKS-NRPS = polyketide synthase / non-ribosomal peptide synthetase; *trans*-ER = *trans*-acting enoyl reductase; HYD = hydrolase; DA = Diels-Alderase; P450m = macrocycle P450; P450c = cyclohexene P450; NAD(P)H-OXR = NAD(P)H-dependent oxidoreductase; OAcT = acetyltransferase; MFS = multifacilitator superfamily transporter; TF = transcription factor; FAD-OXR = FAD-dependent oxidoreductase; BVMO = Baeyer-Villiger monooxygenase; TRX = thioredoxin. Note: *%id refers to percentage identity compared to *pyi* BGC proteins; ^%id refers to percentage identity compared to *ccs* BGC proteins; N/A refers to no homologous proteins in the *pyi / ccs* BGCs.

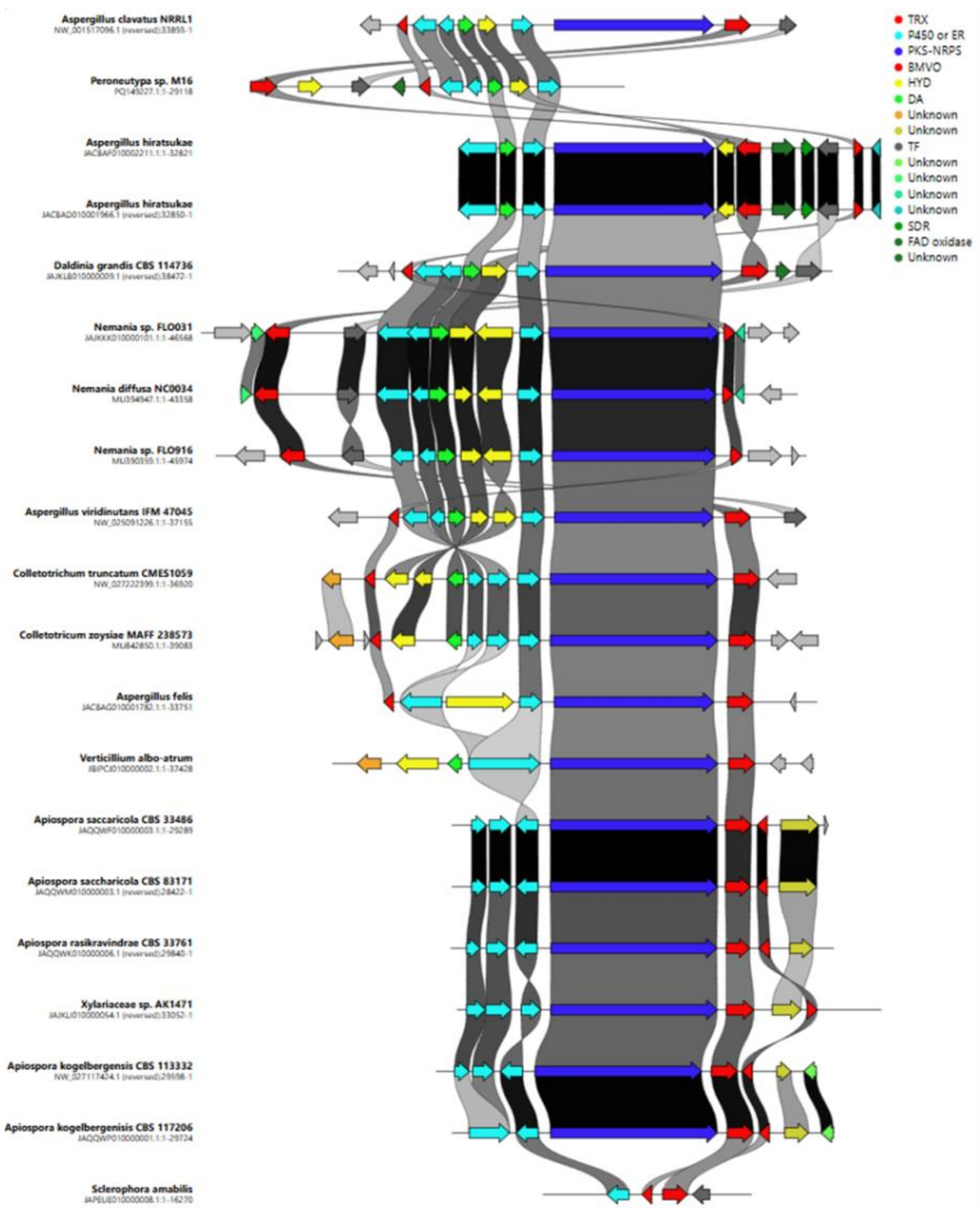


**Figure S3**: Clinker synteny analysis of putative cytochalasan BGCs demonstrating highly conserved BVMO and TRX-like genes (highlighted in red) from diverse fungal species.

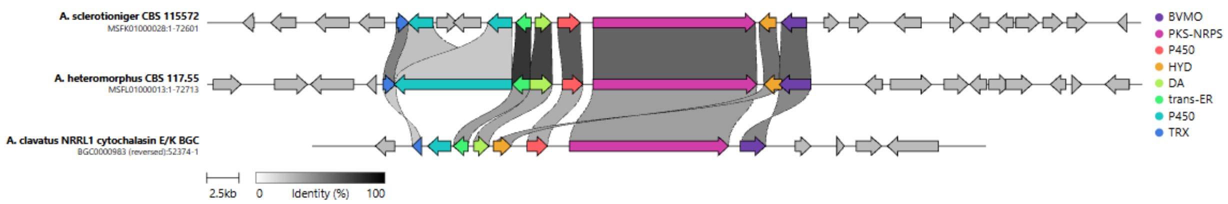


**Figure S4**: Comparison of the *asc*, *ahe*, and *ccs* BGCs from *Aspergillus* sp. using clinker, indicating higher sequency identity between the *ahe* and *asc* BGCs than *ahe* and *ccs* BGCs.

| Protein Name | Amino acid sequence | DNA sequence |
|---|---|---|
| XsCYP1 | MQQLSVKMPSPASIALAIVAVLCSL<br>FYSLVYSGSLTAKKRALAHIPELRFE<br>NDDTPERYRRETRSLLRIGYQKYLQ<br>YGVPFQMHNSVGELGNQVLLPMK<br>YLDEVKRAPSSLYSFEAFSEKLFLLS<br>YFNSPRQTDAVTHAIRLDMNRNLD<br>NVLNALWAEANQLFKETVPATGWK<br>TVPVGELACSIVSRMVSYIIVGPVLC<br>RNPEWQQIAIEATFVIVETALDLRAK<br>YTRNWRWLARWQNGTAQKLGEIR<br>KRASELIKPLYEERRRAMNQVDGR<br>SFYDTIYWVLKRGKADQSLRELVD<br>QQLFLTLTSIHTTGGTLQSILFDWLD<br>HPEYHADIRAEINEALADVESSGGK<br>WTLPRIAAMKKLDSFMKESTRVNPI<br>GFTTIQRYALKSHTFKDGFILPAGTV<br>FQFAMDALHHDPNIYPDPDKFDAY<br>RFLHLREKTDANQFHYAFVSDMTL<br>NWGAGQHACPGRFLATLVLKLTLIL<br>LVTRCEVKFSDGAAENRQDVCVD<br>NSRRVDPTVRLDIKARS | ATGCGAGTCTCGGTCATGCAATTAAAGGGGCAACGCTGTATGGAGTACCAGTACAGTGGGACA<br>AGGCAGCTGGCTCCTAGTCGTCAGCTCTCGCATCTCAGCATCACGTCGACGTACCGAACAGC<br>ACTTGCCTACCTACATTACACATCCTCCTCCTCTTCTTCTGTCTCTTACAATCTACCGAGACATCA<br>AACAACAAACCGCGTTCCTAGTACAACCCTTCGAGCACAATTCCATAACTACCACCCATTCAAA<br>CTACGCCAGCATGCCTGGCTTCGACTTCTCGAACTACAACCGCAATGCAGCGCTGCACGCAC<br>GAGGCGTGCCCTTGCCTAAAGCTACCAGCACAGGAACCACCATTGTTGGGTGTATATACGATG<br>GCGGCGTGGTGGTAAGTTATCCCAGACTCAAACCTAGATCACTTTATCTCCGGCAGGCTCAATC<br>TGATGGAGTTGCTTAGATCGCTGCCGACACGCGAGCTACGAGCGGCCCTATCGTCGCCGACA<br>AGAACTGCGAGAAGCTTCATTATATCTCCCCCCAGATCTGGTGTGCTGGCGCCGGTACAGCAG<br>CCGATACCGAGTTTACCACAGCCCTTATCTCCTCACAGCTCGAGCTACACTCCCTCTCTACCG<br>GCCGTAAACCCCGCGTCGTCACCTGCATGACCCTCCTCAAACAGCACCTATTCCGTTACCAG<br>GGCCACATCGGAGCCTATCTGGTCGTTGCCGGCGTCGATCCTACTGGCACTCATCTCTTTACT<br>GTTCACGCTCACGGTAGCACCGACAAGCTTCCCTATGTCACCATGGGTTCCGGCTCATTAGCC<br>GCCATGTCCGTTTTCGAGTCCCAGTGGAAGCCTAACTGCACCAAGGAGGATGCCATGAATCTC<br>TGCTCCCAGGCTATCCAAGCCGGTATCTTCAACGACCTAGGTTCTGGATCTAATGTTGATCTGGC<br>CGTCATCACAGCCGACAAGACCACACTACACCGCGGATTTGTAAAACCAAACGAACGTAGCC<br>AGAAATTGAAGAACTACAAGTTCCAGCGTGGTACAACGGCAGTGCTGAATGAGAAGATCATCAC<br>CAAGGATGATATCGGGAAATACGTTACTGTTCACAAGCTAGAAGCAACAGGCGAGGAAAAGATG<br>GACATCGACCCATAA |
| XsER | MPISSAVSLPTTRTAIVEGNDHRLRI<br>EHNAPLPTLRLGEVLVHVKAVAINP<br>CDYKMHERFPCAGAVDGCDFSGV<br>IVALGSQVANLSIGDRVCGAVHGS<br>NPIRPESGAFADYIVSESEFTLKVPA<br>EMSFVEAAGLGVTGLATLGMALFKT<br>LALPGTLEEPVKKPRTVLVHGGSSS<br>VGTMAMQLLRLAGHIPITTCSPKNF<br>ALARRFGAEEVFDYNSPDCAGNIK<br>AYTKNTLSYVLDPFTDAKSIALCYGA<br>MGRAGGRYSCLEMYPDYILERKSIK | ATGCCAATCTCGTCCGCTGTTTCGCTGCCGACGACCCGTACTGCCATCGTCGAGGGCAATGAT<br>CACCGCCTCCGTATCGAGCACAATGCCCCACTCCCCACCCTCCGATTAGGTGAAGTATTGGT<br>GCATGTGAAAGCCGTTGCTATCAATCCATGTGATTACAAGATGCACGAGCGCTTCCCCTGTGCT<br>GGGGCAGTCGACGGTTGCGATTTTTCGGGGGTGATTGTCGCGCTCGGGTCGCAAGTGGCAAA<br>TCTTAGCATCGGAGACCGAGTCTGCGGGGCAGTCCACGGCTCGAATCCCATTCGGCCCGAAT<br>CGGGCGCCTTCGCCGATTACATAGTGTCCGAGTCCGAATTTACGCTGAAAGTTCCGGCCGAAA<br>TGTCATTTGTCGAAGCTGCCGGGCTAGGAGTGACAGGTCTGGCGACACTGGGTATGGCGTTGT<br>TCAAGACGCTGGCTTTGCCAGGGACCCTGGAGGAGCCGGTGAAAAAGCCGCGGACGGTGC<br>TTGTCCACGGAGGTAGCTCATCCGTAGGAACAATGGCGATGCAGTTGTTGAGACTGTGAGTACT<br>ATACAGATCTACCGCAACACGACTCATGTAGCTAACAGAACTGGGGCGGAAGGGCCGGACAT<br>ATACCCATAACTACCTGCTCTCCTAAAAACTTTGCCTTGGCCAGACGTTTCGGCGCGGAGGAA |

| | | |
|---|---|---|
| | VGFVMGPALLGHRLALSDGYERDE<br>DPEMRAFGVQWYQGVQKMLDQG<br>KLRPHPLRVLGTSFEAVLEGVEMLK<br>RKAVSGEKLVVTLDCN | GTCTTTGATTATAATTCCCCAGACTGCGCCGGAAATATTAAGGCCTACACCAAAAACACGCTCTC<br>CTACGTGCTAGACCCTTTTACCGATGCCAAGAGCATCGCCCTCTGCTATGGGGCCATGGGTCG<br>CGCTGGAGGCCGCTATTCCTGCCTGGAGATGTACCCAGACTACATTCTCGAGCGCAAATCGAT<br>CAAGGTGGGCTTCGTCATGGGTCCCGCTCTGTTGGGCCACCGGCTCGCGTTGAGCGACGGT<br>TACGAGAGAGACGAGGATCCTGAGATGCGCGCGTTCGGCGTGCAGTGGTACCAAGGTGTTCA<br>GAAGATGCTGGACCAAGGGAAATTGAGACCACATCCGCTGAGGGTTCTGGGCACGAGCTTCG<br>AAGCAGTGCTGGAAGGCGTTGAAATGCTAAAGCGAAAGGCAGTATCGGGTGAGAAATTAGTTGT<br>GACCCTTGATTGTAA |
| XsDA | MLIYLISFTGLLAGTFAKTHGWPEP<br>WPSHWRQPNSDLFPGQQAPLVIE<br>DTDANADCRLSNITAFEMAKGRKIV<br>DFTTKAMGSLEEPKIRPLNSSGGE<br>QWEFDGVSEDGMQSFVFGFYRDA<br>NYAILGTGNFRLSIEFAFADRTRFYE<br>VYYPERSVIETCSQGTRGLWIDEKS<br>GYQFSFLVNADMTEAVITLDSDTVK<br>GKASIYSRALPLAADGSVWPAENA<br>STAPIPYYHWSQPIPAGTVDVDVKIK<br>GKHVQWTGMGGHERFWSAFSWF<br>TCMRNLQAVRAMLGPYVLSYFSFT<br>SNLEQDLTHQSVVLFKDGIPVFRST<br>VGTASETEDYALVTKTYGGAVTGTLK<br>DKVTGFQLELVSPANMRHYTFFVE<br>HLNLGFEYILGEGVGGSGFSGHSR<br>GGHVGLSQYEGIALTEALTFPKNSP<br>LFKSNYVA | ATGCTAATCTATCTCATATCTTTCACTGGTCTGCTGGCCGGTACCTTTGCTAAAACCCATGGCTGG<br>CCGGAACCTTGGCCGTCCCACTGGCGTCAGCCAAACAGCGATCTCTTTCCTGGCCAGCAAG<br>CCCCCCTCGTGATCGAGGACACTGACGCCAATGCGGACTGCCGGCTAAGCAACATAACGGC<br>GTTCGAGATGGCCAAAGGCAGGAAAATTGTGGATTTCACAACTAAGGCTATGGGTAGCCTTGAA<br>GAGCCAAAGATCCGTCCACTTAACTCGTCCGGCGGGGAGCAGTGGGAATTTGATGGAGTCTCT<br>GAGGATGGCATGCAGTCCTTTGTTTTTGGCTTCTACCGAGATGCGAACTACGCCATCCTGGGCA<br>CCGGAAACTTCCGTCTGTCCATCGAGTTTGCATTTGCAGACCGCACGCGGTTTTATGAGGTGTA<br>TTACCCGGAACGGTCCGTCATCGAGACATGCTCGCAAGGGACCCGCGGCCTGTGGATCGAC<br>GAGAAGAGTGGCTACCAGTTTTCTTTCCTGGTCAACGCAGATATGACGGAAGCTGTTATCACACT<br>AGATTCCGATACTGTCAAAGGCAAGGCTTCCATTTACTCACGGGCTTTGCCCCTGGCTGCGGA<br>TGGGAGCGTGTGGCCGGCCGAGAACGCGTCCACAGCACCGATTCCCTACTATCATTGGTCAC<br>AACCAATCCCAGCGGGAACGGTGGACGTCGATGTGAAAATTAAGGGCAAGCATGTCCAATGG<br>ACCGGCATGGGGGGGCATGAGCGGTTCTGGTCCGCTTTCAGGTCAGTCGCGTCTAATGCTCT<br>GTGGCACATCGTAATATCTCAGCCTATTAACCCACTGCTTCTTATATAGTTGGTTTACTTGTATGAG<br>AAATCTCCAAGCAGTGCGGGCCATGTTGGGTCCTTATGTGCTGAGCTATTTCTCCTTTACTTCGA<br>ATCTCGAACAAGATCTCACGCACCAGTCCGTTGTTCTTTTCAAGGACGGTATTCCCGTCTTCCG<br>AAGCACGGTCGGAACTGCCTCCGAAACAGAAGACTACGCTCTCGTCACAAAGACATACGGTG<br>GGGCCGTCACAGGCACTCTCAAGGATAAAGTTACGGGCTTCCAGCTGGAGCTCGTGTCACCA<br>GCAAACATGCGTCACTACACCTTTTTCGTGGAGCATCTTAACCTCGGGTTTGAATACATCCTCGG<br>AGAGGGCGTCGGAGGTAGCGGGTTCTCTGGCCACTCGAGAGGCGGCCATGTAGGCCTCAGT<br>CAGTATGAGGGAATCGCGCTGACCGAGGCGCTGACATTCCCCAAGAATTCACCCCTTTTTAAG<br>AGCAACTATGTAGCATAA |
| XsCYP2 | MLSSIQTNIVEPFLVLRQSVAPLKLS<br>RWQLTKLMIRTALNGLPDGSLFFLL<br>AALAATIAVYYLIFNNTNTRVQVPPG<br>LAVVKRDDMHFLDIIDEGRKLYPG | ATGTTGTCAAGCATCCAAACGAATATTGTCGAGCCTTTTCTGGTGCTGCGCCAGAGTGTGGCAC<br>CTTTGAAACTGTCCAGATGGCAGCTCACGAAGCTTATGATCCGCACTGCACTCAATGGGCTGC<br>CCGACGGAAGCCTCTTTTTCCTCCTAGCGGCCTTGGCAGCGACTATTGCTGTGTACTACCTTAT<br>ATTCAACAATACAAACACACGCGTACAAGTGCCTCCCGGGCTCGCCGTGGTGAAGAGGGATG |

| | | |
|---|---|---|
| | QPFLAVNRRHSFVIYPPQCFDEIKR<br>LPEHTASARAFFHATNYGHWSHVG<br>TETPELIKSVIADLTRSLPARVLARQE<br>DCQNAFDGVLGRRRDWKEFPLM<br>MTTFEIVTQINACSFVGRKLGTSRG<br>WVKSVMMSPIFIHVAVTLLDACPFI<br>LRPLMAPIYFFPTMKNRWDMKRLL<br>TPILEEDIKDFYAATDKKEILRPRPD<br>GKIPFTGFLLSRYRTAEASIRQLISDY<br>ILISFDSTPSTASAFFHALCELALHP<br>EAADILREELDEYVVDGNLPGTHL<br>QELRKMDSFLRESFRLHPIGIFSLQ<br>RVVEKPIKLSVGPTIPPGTIIAVDGQ<br>AINRSPDLWPNPDSFDMDRFYKLR<br>QKPGNENRFHFLTTGSDSPGWGD<br>GTQACPGRFFATSTLKIAMAHFLRN<br>YDIEIKPECLPLKHTPLSNGSWKPD<br>DTAIARIRARC | ACATGCATTTCCTTGATATCATCGATGAAGGACGGAAGCTGGTATGTACAACTTCATAATCCGCCT<br>TCCACCTACTGGGAAAACATTTGTTGGCTCAAGTTCTGACACATCGCAGTACCCAGGTCAACCC<br>TTCTTGGCGGTAAACAGGCGGCATAGTTTTGTCATATACCCGCCTCAATGCTTTGATGAGATCAA<br>GCGTCTTCCGGAGCACACTGCGTCCGCGAGGGCATTCTTCCACGCTACCAACTATGGCCACT<br>GGAGCCACGTTGGCACTGAGACACCCGAACTCATCAAATCGGTCATTGCGGATTTGACCCGTT<br>CGCTCCCTGCTCGAGTGCTCGCTCGTCAAGAAGACTGCCAGAACGCATTCGACGGGGTCCTT<br>GGGCGCCGGCGCGATTGGAAGGAATTCCCTCTGATGATGACTACATTCGAAATCGTCACCCAG<br>ATCAATGCGTGTTCTTTTGTCGGAAGAAAACTAGGCACTAGCCGCGGCTGGGTCAAGTCTGTCA<br>TGATGTCGCCTATCTTCATTCACGTTGCTGTCACGCTCCTGGATGCCTGCCCATTCATCCTCCG<br>GCCGCTCATGGCCCCGATTTACTTTTTCCCTACTATGAAGAACCGATGGGATATGAAGCGACTG<br>CTCACGCCTATTCTGGAGGAGGACATAAAAGATTTCTATGCAGCGACGGACAAGAAAGAAATCT<br>TGCGACCCCGGCCAGACGGCAAGATTCCCTTCACTGGGTTCCTCCTCTCACGCTACCGGAC<br>GGCCGAAGCAAGCATCAGACAGCTGATCTCAGACTACATCCTCATCAGCTTCGACTCAACCCC<br>ATCCACCGCATCGGCATTTTTCCATGCTCTCTGTGAGTTGGCATTGCATCCGGAAGCCGCCGA<br>CATTCTGCGAGAAGAACTAGACGAATATGTCGTTGACGGAAACCTCCCTGGGACTCATCTTCAG<br>GAGCTAAGGAAAATGGACAGCTTCCTCCGCGAGTCCTTCCGGTTGCATCCTATTGGCATATGTA<br>AGTCATGTCCAGCGCCTGGAACAAGCTCGAAGATGCGGTATGTCTTTCTTTAGGAACCGTTATCT<br>AACTTGTCGTTGCAGTCAGCCTCCAACGCGTCGTTGAAAAGCCGATAAAACTCTCGGTGGGCC<br>CGACTATCCCGCCTGGCACAATCATCGCTGTCGATGGACAAGCTATCAACCGGTCCCCGGATT<br>TGTGGCCAAATCCGGATAGTTTCGACATGGACCGATTCTATAAGTTGCGGCAAAAACCCGGCA<br>ACGAAAATCGGTTTCATTTCCTGACCACTGGCTCCGACTCGCCGGGTTGGGGCGACGGACA<br>CAGGCCTGCCCGGGTCGGTTCTTCGCGACAAGCACGTTAAAGATCGCGATGGCGCATTTTCT<br>CAGGAATTATGACATCGAGATAAAGCCAGAGTGTCTGCCGCTGAAGCACACGCCGCTTTCAAA<br>TGGATCTTGGAAGCCCGACGATACGGCAATCGCACGTATTAGGGCCAGATGTTAG |
| XsPKS-NRPS | METIREPIAIIGTGCRFPGQCDTPSK<br>LWDLLRKPRDLLKEIPEDRFSTDAF<br>YHPKNHHHGTTNVRHSYLLDEDL<br>RGFDAQFFGINPVEANSVDPQQRL<br>LLETVYESLEAAGLSAKQLQGSDTA<br>VYVGVMSADFTDMIGRDTEMFPTY<br>FATGTARSILSNRLSYFFDWHGPS<br>MTIDTACSSSLIAMHQAVQTLRAGE<br>SSVAVVAGSNLILGPEQYIAESKLQ<br>MLSPTGRSRMWDADVDGYARGEG | ATGGAGACCATTCGTGAGCCTATCGCTATCATTGGCACCGGGTGCCGGTTTCCTGGCCAGTGC<br>GATACTCCATCTAAGTTATGGGATCTCCTGCGTAAGCCGAGAGATCTTCTAAAGGAGATCCCAGA<br>GGACAGATTCAGCACAGATGCCTTCTACCATCCTAAGAACCACCACCATGGCACGACCAACGT<br>GCGGCACTCCTATCTTCTGGATGAAGACCTGCGCGGATTTGACGCTCAGTTCTTTGGGATCAAT<br>CCGGTCGAAGCCAACTCTGTCGATCCCCAACAGCGACTGCTTCTCGAAACCGTGTACGAGAG<br>TCTCGAAGCTGCCGGCCTCTCAGCGAAGCAGCTACAAGGGTCCGACACGGCCGTCTACGTA<br>GGCGTCATGAGCGCCGACTTTACCGATATGATTGGACGTGACACTGAAATGTTCCCAACATATTT<br>TGCTACGGGGACTGCAAGGAGCATTCTCAGTAACAGGCTGTCCTACTTTTTTGACTGGCATGGC<br>CCGTCCATGACCATCGATACCGCCTGCTCATCAAGCCTCATCGCTATGCACCAAGCTGTGCAG<br>ACTCTCCGTGCGGGGGAATCGTCCGTCGCCGTTGTAAGTACATGTTCCTGTCTAGGAGTATTCC |

| | | |
|---|---|---|
| | VAAIVLKKLSQAIADGDHIECIIRETG | ATTACGTTTCTACTCCGAAGGTGATCATGGGGGCCGCGGGAACGCTACTAACCACATTGTACA |
| | LNQDGRTPGITMPSATAQEALIRTT | GGTCGCCGGGTCCAACCTGATATTAGGCCCAGAGCAATACATCGCCGAAAGTAAACTACAGAT |
| | YAKAGLDISKRADRPQFFEAHGTGT | GCTTTCGCCTACGGGCCGGAGCCGTATGTGGGACGCTGATGTAGATGGCTACGCTCGTGGCG |
| | PAGDPVEAQAVYNAFFGPGSHFKP | AAGGCGTTGCGGCCATTGTCCTCAAGAAACTAAGTCAAGCAATTGCAGACGGCGATCACATCG |
| | SNPDDTLFVGSIKTVIGHTEGTAGL | AGTGCATTATCCGCGAGACCGGTCTGAACCAAGACGGCAGGACACCAGGCATCACTATGCCA |
| | AAVIKASLALQAGVVPPNMLLNKLN | AGCGCTACAGCGCAAGAAGCCTTAATACGAACTACATACGCCAAGGCTGGTTTGGACATTAGC |
| | PKIEPFYGDVRILSAAQKWPKLAEG | AAGCGTGCTGATCGTCCTCAGTTCTTCGAGGCGCATGGCACAGGTACATAAATCTAAAACGCTG |
| | AVRRVSVNSFGFGGANCHAILESF | ACTATTCATTTTCCTCCTGTAGTCATGGAACGTTTTGGCTGATTTGATCTATTAAGGTACTCCTGCTG |
| | EPNETSHKHPSNKVTTCFTPFVFSA | GTGACCCAGTTGAAGCGCAGGCTGTTTATAACGCTTTCTTCGGCCCGGGGTCCCATTTCAAAC |
| | ASESALTARLERYREYLAGSKADAT | CCAGTAATCCAGACGATACCTTGTTCGTTGGCTCTATCAAGACTGTCATCGGTCATACAGAAGGA |
| | VRLRDLSWTLSNRRSTLPWRAVVP | ACGGCAGGTCTTGCGGCTGTTATAAAAGCGTCGCTGGCGCTGCAAGCTGGTGTCGTCCCTCC |
| | ATNDAEDLIRKLDDCTEFTSESSSS | GAACATGTTACTAAACAAACTGAACCCCAAAATCGAGCCTTTTTACGGGGATGTACGGATTCTTT |
| | SSVKTKGSKPRILGIFTGQGAQWPR | CCGCCGCTCAGAAATGGCCCAAGCTTGCGGAGGGTGCTGTGCGAAGAGTGAGCGTCAATAG |
| | MGAELIEKSPAASKILARLEKSLRSL | CTTTGGCTTTGGCGGTGCTAATTGCCATGCCATTCTCGAAAGCTTCGAGCCCAACGAGACATC |
| | PHRDRPTWSLREELLAGAESSSVG | CCACAAACATCCGTCGAATAAAGTTACCACGTGCTTTACCCCATTCGTCTTCTCAGCGGCTTCT |
| | KASLSQPLCTAVQIILVDMLRAAGIE | GAGAGTGCCCTCACCGCTAGGCTGGAAAGGTACCGAGAGTACCTTGCCGGTAGTAAAGCGGA |
| | FSAVVGHSSGEISAAYAAGYLSSED | TGCCACAGTGAGACTCCGAGATTTATCATGGACACTGAGCAATCGCCGATCGACGTTGCCTTG |
| | AIRVAYYRGLHMRSLTQNGAMLAV | GCGGGCCGTCGTACCAGCCACTAATGATGCTGAGGACCTTATTAGAAAGCTCGATGATTGCAC |
| | GTSYEDAKELCELPAFEGRVCVAAS | CGAATTCACGAGCGAGTCTTCTAGCTCCTCTTCAGTCAAGACTAAGGGGTCGAAACCTCGGATT |
| | NSPSSVTLSGDAEAIDEIKTVLDEEK | CTCGGTATTTTCACTGGTCAAGGTGCCCAGTGGCCACGGATGGGCGCTGAATTGATTGAGAAG |
| | KFTRHLKVDRAYHSHHMAACSEPY | TCACCGGCCGCAAGTAAGATTCTTGCACGCTTGGAAAAGAGCCTCCGGTCACTACCACATCG |
| | VSSLQQCGIQQLSPTGARKCRWV | GGATCGGCCAACATGGTCGCTTCGCGAGGAACTCCTCGCCGGTGCCGAGTCGTCATCCGTA |
| | SSVFTCDITDIPVTDGLQGKYWALN | GGCAAAGCTTCCCTGTCGCAGCCTCTATGCACAGCCGTGCAGATTATTCTCGTCGATATGCTCA |
| | LTKPVMFAEALQILLSGGQDGEVY | GAGCGGCAGGTATCGAATTTTCCGCGGTAGTGGGTCACTCGTCCGGTGAGATCAGCGCAGCG |
| | DMAIEVGPHPALKGPARQTIEGCL | TATGCGGCGGGATATCTTTCATCTGAGGATGCCATTCGCGTGGCATACTATCGCGGCCTGCATA |
| | DGQSIPYTGVLSRDKDGIESFSQGL | TGAGGTCCCTCACCCAAAACGGGGCCATGCTCGCAGTCGGAACGTCGTACGAAGACGCCAA |
| | GYIWRTWGEGAVDFASYSRFMND | AGAGTTGTGTGAGCTCCCAGCCTTCGAGGGCCGTGTCTGTGTAGCGGCGAGTAACTCTCCGTC |
| | EQIETGMPQLVPMKDLPPYPWEH | CAGCGTGACACTGTCTGGAGATGCCGAGGCGATTGACGAAATCAAGACAGTCTTAGATGAGGA |
| | NRKFWHESRLSRAFRTGKDQPNE | GAAAAAATTCACTCGCCACCTCAAAGTAGATCGCGCTTATCATTCACATCACATGGCAGCCTGC |
| | LLGRRILDGAPDQLRWRNVLKRNE | TCAGAACCGTACGTATCTTCTTTACAGCAATGCGGGATTCAACAGCTTTCTCCGACGGGGGCTA |
| | IDWLDGHQVQRQTVFPCAGYLSA | GAAAGTGCCGATGGGTATCCAGCGTTTTCACCTGCGATATCACGGACATTCCTGTAACAGACGG |
| | CVEASLKIRRDTNVQSIELQNFVVG | TCTTCAAGGGAAGTACTGGGCGCTGAACCTGACAAAGCCGGTCATGTTTGCCGAGGCCCTAC |
| | QAVAFNDDDSGIETLIVLDNIKEFEE | AAATCCTCCTTAGCGGGGGTCAGGATGGGGAGGTTTATGACATGGCTATTGAGGTCGGACCAC |
| | QGKKTVSAKFAFYSSLNNEILDMTS | ACCCAGCACTTAAGGGTCCCGCAAGACAGACAATCGAGGGATGCCTTGACGGACAGTCCATC |

| | | |
|---|---|---|
| | HASCDVRVTYGDSAANLLPHKMH<br>DIDEDSMLDVESDRFYNVLEQLGF<br>GYSGPFRALTSLKRKLGKAMGHIQ<br>NPESSQLFQKPLLIHPATLDAGIQSI<br>MLAYCYPGDSMMRSIYLPTGIRRLII<br>NPEHCRAFAGQETKVLFDSSASVD<br>TSRSLSGDVSIYSPEGFACKAIQLE<br>GLQTQPLFHPTESNDLNIFTELVW<br>DVDRPDSGEIVGKINVQELNAELLF<br>SLERVAYFYLRFLDKTIPHSERTNLE<br>WHYTRLFAYVDHVLSKVARGANRF<br>AKKEWQHDTKEVILGIFDRFPDNID<br>LRLMRAVGENMPAVVRGEITMLEP<br>MLQDNMLNDFYVVAHGMPRYTAY<br>LAALASQIGHRYPHMHVLEIGAGT<br>GGATKSFLGALGDKFSTYTFTDISG<br>GFFEKAKHVFASHSSKMNFKVLDIE<br>KDIEGQGFAAGSYDVIIASLVLHATR<br>DLAQTLRNVRRLLKPGGYLLLLEITE<br>NDQMRFGLLFGGLQGWWLGYDD<br>GRALSPCIGLEEWSTYLKQTGFSGI<br>DTSMPHDENLPVPLSVIVSQATDD<br>RVELLKQPLREKNTTAIVVPQLTIIG<br>GAALATDVRRILGSYCGSVTFIESLN<br>DLGPDDLPVGGSVLCLSDIEEPVFK<br>SMDRDKLRGFQTIFKQSTSVLWVT<br>QGVRSGDPFSRMVVGFGRTIVLEM<br>LHLRLQFLDLDTDAPADPTEIAESLL<br>RFQMAGNWESDGAENSPLLHSIE<br>PELYLEKSGRVYIPRFKLNRRPNDR<br>YNSARRNITEQISLREKTVELVQRET<br>QDGSWYLVEGKHMPEQEEAVEID<br>VLHSVNRTVEVSRGTFLFPILGRNR<br>DTGETVLALSPKQASRVRIPPAFVIP | CCTTACACGGGCGTGTTATCACGCGACAAAGACGGTATCGAGTCATTCTCACAAGGACTCGGA<br>TATATCTGGCGAACGTGGGGGGAAGGAGCTGTGGATTTTGCCTCCTACAGTCGCTTTATGAATG<br>ACGAGCAAATAGAAACGGGCATGCCACAACTCGTTCCTATGAAGGATCTCCCTCCTTACCCGT<br>GGGAACACAACCGGAAGTTTTGGCACGAGTCTCGCCTCTCTCGTGCTTTCAGAACCGGCAAA<br>GACCAACCCAATGAGCTCCTCGGGCGTCGAATCCTCGACGGCGCTCCCGACCAGCTCCGTT<br>GGCGCAACGTTCTCAAGCGCAACGAGATAGATTGGCTCGATGGTCACCAAGTCCAGCGACAG<br>ACTGTTTTCCCTTGTGCCGGCTATTTGTCCGCTTGCGTGGAAGCCTCGCTGAAGATTCGTAGAG<br>ACACTAACGTACAATCTATTGAACTCCAAAACTTCGTGGTTGGTCAGGCTGTGGCGTTTAATGATG<br>ATGACTCGGGGATTGAGACGTTGATCGTACTAGATAACATCAAGGAATTCGAAGAACAAGGCAA<br>AAAAACGGTGTCTGCGAAGTTTGCCTTCTACTCGTCCCTTAACAATGAGATTTTGGACATGACCA<br>GTCACGCTAGCTGTGACGTCCGTGTCACGTACGGGGACAGCGCCGCGAATCTTCTCCCACAC<br>AAAATGCACGATATTGACGAGGATTCCATGTTGGATGTTGAGTCCGATCGGTTTTATAACGTCCTC<br>GAACAGCTTGGATTTGGATACTCTGGTCCATTCCGGGCGTTAACAAGCCTGAAACGAAAGTTGG<br>GCAAGGCGATGGGTCATATTCAGAACCCGGAAAGCAGTCAGCTTTTCCAAAAGCCTCTCCTCA<br>TCCACCCAGCCACTTTAGATGCAGGAATCCAGTCCATCATGCTAGCATATTGCTACCCGGGAG<br>ACAGCATGATGCGCTCCATATATCTGCCCACCGGTATACGCCGACTCATCATCAACCCAGAGC<br>ATTGTCGAGCCTTTGCCGGTCAGGAGACAAAAGTTCTGTTCGACTCGTCAGCGTCGGTCGACA<br>CCTCTCGGAGTCTATCAGGCGACGTGAGTATATACTCCCCGGAGGGCTTCGCCTGCAAGGCC<br>ATTCAGCTCGAGGGACTCCAGACGCAACCTTTGTTTCACCCGACCGAGTCGAACGATCTAAAC<br>ATCTTCACCGAACTTGTGTGGGACGTTGATCGACCCGACAGCGGGGAAATTGTCGGCAAAATA<br>AATGTCCAGGAGCTCAATGCGGAATTGTTATTTTCCCTCGAGAGAGTGGCCTACTTCTACCTCCG<br>ATTTTTGGACAAAACCATTCCTCATAGTGAGCGTACTAACCTCGAATGGCATTACACCCGCCTTT<br>TTGCCTATGTAGATCATGTCCTCTCCAAAGTGGCCCGCGGTGCGAATCGCTTTGCCAAAAAGGA<br>ATGGCAGCACGATACCAAGGAAGTTATACTCGGTATCTTTGACAGATTCCCTGACAACATTGACC<br>TGAGGCTCATGCGCGCTGTGGGTGAGAACATGCCTGCCGTTGTCCGTGGCGAGATCACTATG<br>CTAGAGCCTATGCTCCAGGACAATATGCTCAACGACTTCTACGTAGTCGCACATGGCATGCCTC<br>GTTATACTGCGTATCTTGCCGCTCTTGCCAGCCAGATTGGCCACAGGTACCCTCATATGCATGT<br>CCTCGAGATTGGTGCCGGTACAGGCGGCGCGACGAAATCGTTCCTCGGAGCGCTTGGTGATA<br>AGTTCTCAACCTACACCTTCACCGACATTTCTGGTGGCTTTTTCGAGAAAGCAAAGCATGTGTTT<br>GCATCACATAGCTCAAAAATGAACTTCAAAGTGCTTGACATCGAGAAGGATATTGAAGGCCAGG<br>GATTCGCCGCGGGCTCCTACGATGTCATCATCGCCTCCTTGGTTCTCCATGCCACGCGCGACT<br>TGGCTCAAACCCTTCGCAATGTACGTCGTTTGCTGAAGCCCGGTGGATACCTGCTACTTCTCGA<br>GATTACAGAAAACGATCAGATGAGGTTTGGACTACTTTTTGGTGGTCTCCAGGGCTGGTGGCTC<br>GGCTATGATGATGGGCGGGCCCTGTCTCCTTGTATTGGATTGGAGGAATGGTCGACATATCTGA |

| | | |
|---|---|---|
| | TQKSAGYLQLFYTELLTRAAIRDVSA<br>GTLIIVLHPSKMLVRAMDRIAPDKG<br>ARVFYLAPEPGSEWDYIHPNATKA<br>DIQNLITSKIGTIPPSSVLLLDMGAD<br>RPLSAGLLECLPAEVAQVKVGTLLT<br>SSIARIAPGHLEQEIRSILVEIKYSLW<br>PAQQAMNTCEGPDELEVVTLEDLT<br>TQDYVAGTDARVVSWPPESNTVPV<br>HVEPVDTRVKFRKDRTYWLVGLTG<br>GLGLSLCEWMAKQGARYIVISSRN<br>PKVDRRWVHKMKTLGVNVEVISN<br>NICDRESVRSVYKKICQEMPPIAGV<br>AQGAMVLHDTIFSELDMERVNKVT<br>QPKVNGSSYLEEIFHDINLDFFVFF<br>SSMACVTGNPGQSAYAAANMFMS<br>GLAAQRRRRGLNASAVHIGAIFGN<br>GYVTRELTLAQQEFLRKVGNMWLS<br>EQDFRQLFAEAIYAGHPGNGASPE<br>LSTGLMMIDNSDDSKKNITWFYNP<br>MFQHCIKESHYSEMVSDGQKGRSI<br>PVKSQLQEAVNPAEVYEIIHDAFAN<br>KLRLSLQIEENRPIVDLTADTLGIDSL<br>FAVDIRSWFIKELQLEIPVLKILGGAT<br>VGDILETAQQLLPKELTPNLDPNDK<br>GAARKRKPQVTSSTKEEAAEQTTKS<br>VSAADDKAGDHRADATKARAVPPL<br>SVQWNNIPKPGTPSGDSDNKSLSS<br>ESGVKVRVASLDTTYTEKSTSPSNT<br>GSDVDAYWGRSRSSVWSMDTTES<br>EVAVSKKTPITFGQSRIWFLEMYLK<br>DPASALNITLTIDLDGSLDVNRFER<br>AVKLVGQRHEALRTRFVTGDNSTQ<br>VMQEVLVDSTLVLEQQDIVGDVES<br>ERIYRELQRYRYKLAEGENMRILLLK | AGCAAACCGGCTTCTCGGGCATTGACACATCAATGCCGCATGATGAAAATTTGCCTGTTCCTCT<br>TTCCGTGATTGTGTCTCAGGCAACGGACGACAGGGTTGAGCTTTTGAAGCAGCCTCTACGGGA<br>AAAGAATACAACGGCTATCGTCGTACCACAATTGACGATCATCGGTGGAGCTGCACTGGCAAC<br>AGACGTCCGCCGGATTTTGGGCTCGTATTGCGGCAGTGTCACGTTCATTGAGTCGCTGAACGA<br>CCTGGGACCAGACGATCTTCCTGTTGGAGGCTCTGTACTCTGTCTTAGCGACATTGAGGAGCC<br>GGTCTTCAAGTCCATGGACCGGGACAAGTTGCGTGGATTCCAAACAATCTTCAAGCAATCCAC<br>CAGCGTCCTCTGGGTTACTCAGGGAGTTCGTTCTGGAGATCCATTCTCCAGGATGGTTGTCGGC<br>TTCGGGAGGACCATCGTACTAGAGATGCTACATCTGCGTTTACAGTTTCTTGACCTTGACACTGA<br>TGCACCTGCGGATCCTACCGAAATTGCCGAATCTCTGCTGCGCTTCCAGATGGCAGGCAATTG<br>GGAGAGCGACGGCGCGGAAAATTCACCTTTGCTTCATTCCATTGAACCTGAATTGTATCTTGAG<br>AAGAGCGGCCGGGTTTACATTCCCCGGTTCAAGCTAAACAGGAGACCGAACGATCGGTACAA<br>CTCGGCTCGCCGTAACATCACCGAGCAGATTTCTCTCCGAGAGAAGACTGTCGAGCTTGTCCA<br>GCGTGAAACGCAAGACGGGTCATGGTACCTCGTGGAGGGCAAACATATGCCTGAACAAGAGG<br>AAGCCGTAGAAATAGATGTCTTGCATTCAGTCAACCGGACAGTAGAGGTGTCGAGGGGCACCT<br>TTCTATTCCCTATTCTTGGACGGAATCGAGACACCGGAGAAACCGTGTTGGCCCTTTCACCGAA<br>GCAAGCCTCTAGGGTCAGAATTCCGCCAGCTTTTGTTATCCCTACACAGAAGTCGGCAGGTTAT<br>CTTCAGCTATTCTATACGGAGCTTCTTACCCGTGCTGCCATAAGAGATGTCTCGGCAGGAACAC<br>TTATCATCGTATTACATCCAAGCAAAATGCTTGTCCGTGCTATGGATCGCATTGCACCAGACAAA<br>GGGGCAAGAGTTTTCTACCTCGCCCCTGAACCTGGCTCGGAATGGGACTACATCCATCCCAAT<br>GCCACCAAGGCGGACATCCAGAACTTGATTACCTCAAAGATCGGAACGATTCCTCCATCTTCT<br>GTGTTGCTTCTAGACATGGGTGCAGACAGACCCCTCTCTGCAGGTCTATTGGAATGCCTGCCG<br>GCAGAAGTTGCGCAAGTCAAAGTCGGTACTCTGCTTACCTCCTCCATAGCACGCATTGCGCCT<br>GGTCATTTAGAGCAGGAAATTAGGTCTATTCTGGTAGAGATTAAGTATTCTCTGTGGCCAGCACAG<br>CAAGCAATGAATACTTGTGAGGGACCTGATGAGTTGGAAGTTGTTACCCTGGAGGACTTGACTA<br>CCCAGGATTATGTCGCCGGCACCGATGCGCGTGTGGTCTCTTGGCCACCCGAGTCTAACACC<br>GTTCCGGTTCATGTTGAGCCTGTCGATACCAGGGTCAAATTCAGGAAGGACAGGACATATTGGC<br>TTGTTGGCCTTACCGGCGGTCTTGGCCTCTCACTCTGCGAGTGGATGGCTAAACAGGGCGCTC<br>GGTATATAGTCATTTCCAGTCGGAACCCTAAAGTAGACAGACGGTGGGTGCACAAGATGAAAAC<br>CCTAGGTGTAAATGTTGAAGTTATATCCAAGTATGTACCGTCAACCTACTTCTGTATTGCTATCCAC<br>TAACCGCGCTACTCTAGCAATATCTGCGACCGAGAGTCTGTCCGATCTGTTTATAAAAAGATCTG<br>CCAGGAAATGCCTCCTATTGCCGGAGTTGCCCAAGGCGCCATGGTCCTTCACGATACCATATT<br>CTCCGAACTCGACATGGAACGAGTGAACAAAGTTACGCAGCCGAAAGTTAACGGCAGCAGCT<br>ATCTAGAGGAAATATTCCACGACATAAACCTCGACTTTTTCGTCTTCTTCTCGTCTATGGCCTGCG<br>TGACCGGCAATCCAGGCCAATCCGCTTACGCGGCTGCTAACATGTTCATGTCAGGACTGGCA |

| | | |
|---|---|---|
| | KSSKSFRLIIGYHHINMDGISLEVVL<br>RELQMAYDSKRLPSVSSILQYPTFA<br>EQQHREFESGKWQDEIAFWRNEF<br>GGRTPPVLPLLPLSKTRSRTALTSYS<br>THTAEFRLDQEALARVQSACEGSK<br>ATPFQFYLTVFYTLLFRLVDAEDICIG<br>ISSANRQDTAMMQSVGMYLNLLPL<br>LLKSHPHETFASTLKLIRSKAMAAFA<br>HSKVPFDVIVNELGVPRATTHSPLF<br>QVLANYRPGTSERRDFCDCRSEVV<br>SFEQGQAAYDLSIDIIENPGGECRII<br>LAGQSALFEPQDMEMLKNMYKHL<br>LVAFSRNPALRLSTPSLYDPEDVKG<br>ALQLGRGSFHTHQWPETLVHRIDE<br>MVERYGSKPAVIDGHGTSLTYSHLA<br>RRVNAIAASMRRIGSGNRVGVYVD<br>PGADWICSFLAILRRGAVYVPLDAV<br>AGSGRLLAILQDSKPDLLLVDNSTE<br>KDAKSQFVSLLAADQILNVDNVSTT<br>TTGTLNNAAKADSVAALMYTSGST<br>GVPKGIIMKHASFRNNIEIITSKLGY<br>REGQEVTLQQSSLNFDMSVFQVFL<br>ALSNGGAVYIVPKHLRADPVAISSII<br>ASRGITSTTATPSELISWIHYGNVGE<br>LRDSNWRSVQSGGEPVRDSLEAA<br>FRKMDKLDLRLIDCYGPTEITFCCH<br>TREIEYRGQGTSSNTGLEVLPNYST<br>YIVDASMKAVPVGIPGEILIGGAGVV<br>AGYLHTELNARGFAHDSFASPEFH<br>KQGWTRLHRTGDFGRISKVNGRLL<br>LEGRIADDTQVKLRGLRLDLQEVES<br>AIIQAAKGSIVDCAVSVRQFETTADE<br>YLVAFATTASATRVENLDQIVHQLPL<br>PQYMRPAALVLLDKIPTNASGKIDR | GCTCAGCGCAGGCGCAGAGGCCTCAATGCATCTGCAGTGCATATAGGCGCCATCTTCGGCAA<br>CGGCTACGTGACTAGAGAGCTGACCTTAGCACAGCAGGAATTTCTCCGAAAGGTCGGCAACAT<br>GTGGCTCTCGGAACAGGACTTCCGCCAACTCTTTGCTGAGGCAATCTATGCCGGTCATCCTGG<br>AAACGGAGCATCTCCGGAGCTCTCCACCGGTTTGATGATGATAGACAACAGCGACGATTCTAA<br>GAAGAACATCACATGGTTCTATAATCCGATGTTCCAACACTGTATTAAGGAAAGCCACTATAGCG<br>AGATGGTTAGCGACGGCCAAAAGGGCCGGAGCATCCCAGTCAAGTCACAGCTTCAAGAAGC<br>AGTCAATCCGGCTGAAGTATACGAGATAATCCATGATGCTTTTGCTAACAAACTACGCCTGAGCT<br>TGCAAATCGAGGAGAACCGTCCCATTGTTGACCTCACGGCGGACACCCTTGGCATTGATTCCC<br>TCTTTGCTGTCGACATTCGGTCGTGGTTCATCAAAGAGCTTCAGCTCGAGATCCCAGTGCTTAAA<br>ATTCTAGGTGGAGCAACGGTGGGCGACATTCTTGAGACTGCCCAACAGTTATTGCCCAAGGAA<br>TTGACACCAAATCTGGACCCTAACGACAAAGGCGCAGCGAGAAAACGGAAACCACAGGTTAC<br>TAGCTCAACGAAAGAAGAAGCAGCGGAACAGACCACGAAGAGTGTTTCGGCAGCGGACGATA<br>AGGCCGGCGACCACAGAGCTGACGCCACGAAAGCAAGAGCCGTCCCACCACTCTCGGTGC<br>AGTGGAATAACATACCTAAGCCCGGCACTCCCTCCGGAGACTCGGATAATAAGTCACTGTCCA<br>GTGAAAGCGGCGTGAAAGTCAGAGTTGCAAGCCTTGATACGACTTACACTGAGAAATCCACTTC<br>CCCCTCGAATACAGGGTCGGACGTTGATGCGTATTGGGGCCGATCCCGAAGCTCAGTGTGGT<br>CCATGGACACAACGGAGAGTGAAGTAGCCGTCAGTAAGAAAACCCCTATTACATTTGGCCAGT<br>CTCGAATCTGGTTCTTAGAAATGTACCTGAAGGATCCAGCTTCTGCACTTAACATTACTTTGACCA<br>TCGATCTCGATGGCTCTCTGGACGTCAACCGGTTTGAGCGAGCAGTGAAGCTTGTTGGGCAGC<br>GCCACGAGGCATTACGAACTCGATTCGTCACTGGCGATAACTCCACTCAAGTCATGCAAGAAG<br>TGCTAGTTGATTCGACGCTGGTTCTAGAGCAACAAGACATCGTCGGCGATGTGGAATCGGAAC<br>GGATCTATCGCGAGCTCCAACGGTACAGATATAAACTGGCGGAGGGGGAGAATATGAGGATCT<br>TGCTTCTCAAGAAATCCTCGAAATCATTCCGCCTTATCATTGGATACCATCACATCAACATGGATG<br>GCATCAGTTTAGAAGTAGTGCTTCGTGAGCTGCAAATGGCGTATGACTCCAAGAGACTCCCGAG<br>TGTGAGCAGTATTCTCCAGTATCCAACTTTCGCCGAGCAGCAGCACCGAGAATTCGAATCTGGA<br>AAATGGCAAGATGAGATAGCATTTTGGAGGAACGAATTTGGCGGCCGCACGCCGCCTGTTCTT<br>CCACTCCTACCTCTATCCAAGACCCGGTCGCGCACTGCGCTTACGTCTTACTCAACCCACACC<br>GCAGAATTTCGGCTGGATCAAGAGGCACTCGCTCGCGTTCAGTCGGCCTGTGAAGGGTCAAA<br>GGCTACACCGTTTCAGTTTTACCTGACGGTCTTTTACACTCTATTGTTCCGGCTGGTGGATGCGG<br>AGGATATTTGTATTGGTATCAGCTCTGCAAACAGGCAAGATACGGCAATGATGCAGAGTGTCGG<br>CATGTACTTGAATCTCTTACCCCTTCTTCTCAAGTCCCATCCGCACGAAACTTTTGCCAGCACTT<br>TAAAACTCATACGCAGCAAGGCAATGGCTGCATTCGCTCACTCCAAGGTTCCCTTTGATGTCATT<br>GTGAACGAGCTTGGTGTCCCACGCGCAACCACGCACAGCCCGCTCTTCCAGGTCTTGGCCA<br>ACTACCGACCAGGTACGTCAGAGCGGAGAGATTTCTGTGACTGCCGCAGTGAGGTGGTATCGT |

| | | |
|---|---|---|
| | SALRSIPLPQTNKSNGLEGQHSSG<br>DNLSSTESRLKRLWEGVLSKEIVSQ<br>HEISTTSDFFHVGGSSMLLISLRADI<br>QETFNVVVSLFQLFDASTLGGMATL<br>INGLSNNGSSDSVQEQEQEQRSD<br>VDINWENETAVSPSLLNVPVHKRF<br>FTNPEVVVLTGSTGFLGQAILTRLLD<br>DGIVKKIHCLAVRHDIPLFNSPKVV<br>VHRGDLGLPRFGLSEEELSKIFSEA<br>DAIIHNGTDVSFAKSYHTLKPANVE<br>ATKELVRLSLPHQTSFHYVSTAAVA<br>NLTGQDSWEQRSVGGFLPPAGAD<br>GYLATKWVSERYLEKVNDQCELPI<br>WIHRPSSITGPNAPATDLMENLIQF<br>SRKIAAIPDTSSWRGWLDFISVDRA<br>AIQIVDEVYEDYSWPGHVKYLYESG<br>EQVVALSDMKGVLERENGTVIETVS<br>MEEWVSRAEEEGLNPLLGEYLKRA<br>SGTPLVFPMLVRHDSFF | TCGAGCAGGGACAGGCGGCGTATGATCTCAGCATTGACATTATTGAGAATCCCGGAGGCGAGT<br>GCCGTATCATCTTGGCGGGTCAGTCAGCTCTGTTCGAGCCACAAGATATGGAGATGTTGAAGAA<br>TATGTACAAACATTTACTGGTCGCGTTTTCTCGCAACCCAGCTCTCCGGTTGAGTACTCCATCAC<br>TCTATGATCCAGAGGATGTGAAGGGCGCACTTCAACTTGGACGAGGTAGGTCATAGATAGTGATT<br>GCTTGCTTCTTTGAAAGCCACTAATATTATTTGAACAGGTTCCTTTCACACGCACCAGTGGCCTG<br>AGACGCTGGTTCACCGCATCGATGAGATGGTCGAGCGATATGGCAGTAAGCCAGCTGTCATAG<br>ACGGTCACGGAACCTCTCTAACTTATTCGCACCTGGCGAGGAGAGTAAATGCCATTGCTGCTTC<br>GATGCGTCGCATTGGTAGCGGGAACCGAGTCGGTGTCTACGTTGATCCTGGAGCGGACTGGAT<br>TTGTTCTTTCCTCGCTATTCTGCGCCGTGGTGCAGTATACGTGCCGCTCGATGCTGTGGCCGGA<br>TCGGGACGACTGCTCGCTATTCTCCAGGACAGTAAGCCGGATTTACTGCTTGTTGACAACTCAA<br>CCGAGAAAGATGCCAAAAGCCAGTTTGTGTCGCTGCTAGCAGCAGATCAAATCCTCAACGTGG<br>ACAATGTTTCGACGACCACGACTGGCACCCTTAACAATGCGGCCAAAGCGGATTCGGTCGCC<br>GCCCTTATGTATACCAGTGGTTCGACAGGTGTTCCTAAAGGGATCATAATGAAACATGCGTCTTTC<br>CGGAATAACATCGAAATCATAACGAGTAAGCTGGGCTATCGCGAGGGACAAGAGGTTACGCTA<br>CAGCAAAGCTCGTTGAACTTCGATATGTCGGTCTTTCAGGTGTTCCTAGCACTGTCAAATGGTGG<br>AGCAGTCTACATTGTGCCGAAACATCTCCGAGCGGACCCTGTGGCCATCTCGTCCATTATCGC<br>GAGCCGCGGGATTACTTCCACGACTGCGACCCCTTCGGAGCTCATTAGCTGGATTCACTACG<br>GTAACGTAGGCGAACTGCGCGATTCAAATTGGAGATCTGTTCAAAGCGGCGGTGAGCCGGTC<br>CGGGACAGTCTCGAGGCCGCTTTCCGGAAGATGGACAAGCTAGACCTGCGCCTGATTGATTG<br>CTACGGTCCGACCGAGATCACGTTCTGTTGCCACACCAGGGAAATTGAATACCGAGGGCAAG<br>GCACCAGTTCAAATACGGGACTCGAGGTCTTACCTAACTACTCTACCTACATCGTAGACGCTAG<br>CATGAAAGCCGTTCCCGTTGGCATTCCTGGAGAAATTCTCATCGGAGGAGCTGGTGTCGTCGC<br>TGGTTATCTACACACCGAGCTAAACGCGCGAGGTTTTGCACACGACAGCTTCGCCTCTCCCGA<br>ATTCCACAAGCAAGGCTGGACACGGCTTCACCGGACAGGTGATTTTGGGCGCATCAGCAAGG<br>TCAATGGGCGTTTATTGCTTGAGGGACGCATCGCGGACGATACGCAGGTGAAGCTCAGGGGG<br>CTTCGGCTCGATCTACAAGAGGTGGAGTCTGCGATCATCCAGGCTGCAAAGGGAAGCATCGTT<br>GATTGTGCCGTAAGTGTACGGCAATTCGAGACAACAGCAGACGAGTATCTAGTCGCCTTCGCC<br>ACCACCGCCTCGGCTACCAGGGTGGAAAATCTCGACCAGATCGTGCACCAACTTCCATTGCC<br>TCAGTATATGCGGCCAGCGGCGCTAGTTCTTTTGGATAAGATACCCACGAATGCTTCAGGCAAG<br>ATTGATCGGTCTGCTCTTAGGTCTATTCCACTTCCCCAAACGAATAAAAGCAACGGACTAGAAG<br>GTCAGCATTCTAGCGGCGATAATTTGAGTAGCACCGAGTCGCGTCTAAAACGATTATGGGAGGG<br>TGTCCTCTCCAAAGAGATCGTGTCCCAACATGAAATCAGCACTACGTCTGATTTCTTCCATGTTG<br>GCGGTAGCTCGATGCTTCTCATAAGCTTGCGAGCAGATATTCAAGAGACTTTCAACGTAGTGGTC<br>TCTCTGTTCCAGCTCTTCGACGCTAGCACGCTTGGGGGTATGGCAACTCTCATCAATGGCCTTT |

| | | |
|---|---|---|
| | | CAAACAATGGCAGCAGTGATTCGGTACAAGAACAAGAACAAGAACAAAGAAGCGATGTCGATAT<br>CAACTGGGAGAACGAGACTGCAGTCTCTCCCAGTTTACTGAACGTGCCGGTACATAAGCGGTT<br>CTTTACGAATCCGGAAGTCGTTGTCCTTACCGGCTCGACCGGCTTCCTCGGCCAGGCGATCCT<br>TACGCGTCTCCTGGACGACGGAATTGTTAAGAAGATACATTGCCTTGCTGTCCGGCATGATATTC<br>CGCTATTCAACTCACCCAAGGTTGTTGTTCATCGTGGAGATCTGGGGTTGCCAAGATTTGGTCTA<br>TCTGAGGAAGAACTCTCTAAAATATTCTCCGAGGCTGATGCAATAATTCACAATGGTACAGACGT<br>CTCCTTCGCAAAGTCATACCACACGCTCAAGCCAGCCAATGTGGAGGCTACCAAGGAACTCG<br>TTCGACTGAGCTTGCCCCATCAGACATCCTTCCACTATGTCTCGACTGCCGCGGTTGCGAATCT<br>GACCGGACAGGATAGTTGGGAGCAGCGGTCTGTTGGCGGGTTTCTGCCGCCTGCCGGAGCG<br>GATGGGTATCTAGCGACAAAATGGGTTTCTGAGCGGTACCTCGAGAAGGTCAACGACCAGTGC<br>GAGCTACCGATCTGGATCCACCGCCCATCCTCGATAACCGGTCCAAACGCACCCGCTACGG<br>ACCTCATGGAAAATCTGATCCAGTTTTCTCGCAAGATCGCAGCCATCCCTGACACGAGTTCGTG<br>GCGCGGCTGGCTTGACTTCATCTCGGTGGACCGGGCAGCGATACAGATTGTCGATGAGGTGTA<br>CGAAGACTACTCGTGGCCGGGCCATGTCAAATACCTGTACGAGTCAGGCGAGCAGGTCGTTG<br>CGCTGTCCGACATGAAGGGCGTGCTGGAGAGGGAGAACGGGACTGTCATTGAGACCGTCTC<br>GATGGAAGAATGGGTAAGCAGGGCAGAGGAGGAAGGATTGAACCCATTGTTGGGGGAGTATCT<br>GAAGCGTGCCTCTGGCACTCCGCTTGTCTTTCCCATGCTAGTGCGACACGATAGCTTCTTCTAG |
| XsOXR | MASWDHRHHMPNLQGRTALVTG<br>GNAGVGYQAVVFLAKAGAKVYFGA<br>RSTERAEAACEKMYQENPEVRQG<br>QIKWLLMDMASMKSILAACEKLRG<br>SESRLDLLINNAAHEGTEPSKLADS<br>GVQVTMQTNHVGVFALTQQLQPLL<br>RAAALEKDSDVRIVMVSSDAPSFS<br>HAEEYRPDFSNPHGGDLMYPSGQ<br>EDSFMGCIRRYSVSKMALNLQASE<br>LQARYDREGVPILVISVCPGTIWTP<br>GTKRAVPWYLYPVFWLKSYSEVMG<br>PRPVLFAAVSREVRAQERYYKGQFI<br>NRSHLVIKGHPATHNAALARELWS<br>STEEIVGKSGQTNG | ATGGCCTCTTGGGATCATAGACATCATATGCCAAACCTGCAAGGGAGAACTGCCCTCGTGACA<br>GGCGGCAAGTAAGCAACTACGTGTTTATTGCTGACAGATCCGTCTACCTATTGACCCTCACTGA<br>CCTCGGGACACAATGTAGTGCAGGTGTTGGTTACCAAGCCGTAGTCTTTCTTGCAAAGGCCGG<br>TGCAAAGGTGTATTTTGGGGCTCGCTCAACCGAAAGAGCCGAGGCGGCATGTGAGAAAATGTA<br>CCAGGAGAACCCCGAAGTGCGCCAAGGGCAGATCAAGTGGCTGCTCATGGACATGGCGAGC<br>ATGAAGAGCATTTTGGCTGCGTGTGAGAAGCTCCGCGGCTCGGAGTCGAGACTAGATCTCCTG<br>ATCAACAATGCCGCCCACGAGGGAACTGAGCCGTCCAAACTGGCTGATTCAGGCGTCCAAGT<br>CACTATGCAGACAAAGTAGGCCCGACATATCTTGCCTCTCTATTCATTCGAGCTGACCTTGGAC<br>GAAGCCACGTGGGCGTTTTTGCTCTGACACAACAGCTTCAGCCACTACTTAGAGCTGCCGCG<br>CTGGAGAAAGACTCTGACGTTCGTATTGTCATGGTATGTAGCCCAATGAACGGCTTCGGAGCAA<br>ATTGGTGCGGAATCTGTGATGGAGTGAAAGCTGATGCTCGAAAACAGGTGAGCTCAGATGCTCC<br>TTCATTTTCTCATGCCGAAGAGTACCGTCCGGACTTTTCCAACCCTCACGGCGGGGACCTGAT<br>GTACCCGAGCGGCCAAGAGGATAGCTTCATGGGTTGTATTAGGAGGTATAGCGTGTCCAAAATG<br>GCCTTGAACTTGCAGGCATCAGAGCTGCAAGCTCGTTACGACCGGGAGGGCGTGCCTATACT<br>TGTGATATCCGTCTGCCCCGGGACGATATGGACGCCCGGCACCAAGCGCGCTGTGCCGTGG<br>TATCTCTACCCGGTCTTTTGGCTCAAGAGTTACAGCGAGGTCATGGGCCCTAGGCCGGTGCTG<br>TTCGCTGCGGTGTCTCGTGAGGTCCGGGCACAAGAAAGGTACTATAAGGGGCAGTTCATCAAC |

| | | |
|---|---|---|
| | | AGGAGCCATTTGGTGATCAAGGGTCACCCGGCCACGCACAACGCTGCTCTGGCAAGGGAAC TATGGTCGTCGACTGAGGAAATTGTCGGCAAATCCGGACAAACCAATGGATGA |
| XsHYD | MHKLFPRSGFFDFETVRILGTTAYG GADVAEVLEAVGEIRSDDPASWEA AWRTQAQRAEKLADEARQHGDRD AALRGGYMYVSSLNEDGDLVQDP RALPIAEKVGQLFRTAMPLMEGETR VLSIPYDHYVLPGYLYLPPKSRRIPG RKKVPILVNTGGADSCQEELFYLNP AAGPGMGYAVLTFEGPGQGIMLRK YELEMRPDWEVVTSSVIDCLEVYSA QHPELELDLSCIAVSGASMGGYYA LRAASDPRVKACVSIADPFYDMWD FGTAHVSPLFISAWTSGIISSGFVDK LMAVISRLSFQLKWEISVAGTFFGL SSPAQILLNMKKYTLHGNGRDEKD VSFLSKVECPVLLSGAGKSLYLDVD NHTKRCYDALTSVALEDKEIWVPES EGQGSLQAKMGAFALCNQKTYRFL DKAFGIKREPLKGGLMHKLFPRSG FFDFETVRILGTTAYGGADVAEVLEA VGEIRSDDPASWEAAWRTQAQRA EKLADEARQHGDRDAALRGYLRAS NYTRASGYMYVSSLNEDGDLVQD PRALPIAEKVGQLFRTAMPLMEGET RVLSIPYDHYVLPGYLYLPPKSRRIP GRKKVPILVNTGGADSCQEELFYL NPAAGPGMGYAVLTFEGPGQGIML RKYELEMRPDWEVVTSSVIDCLEV YSAQHPELELDLSCIAVSGASMGG YYALRAASDPRVKACVSI | ATGCATAAGCTCTTTCCCCGGAGCGGCTTCTTCGACTTCGAGACCGTTCGCATTCTCGGCACA ACGGCGTACGGCGGTGCGGATGTTGCCGAGGTGCTCGAGGCCGTGGGCGAGATCAGGTCA GACGACCCGGCCTCCTGGGAGGCCGCTTGGCGCACGCAGGCGCAGAGGGCCGAGAAGCT CGCCGACGAGGCCCGTCAGCATGGAGACCGCGACGCCGCACTGCGCGGGTATTTGCGCG CGTCAAACTACACGCGAGCCAGCGGCTACATGTACGTTTCGTCGCTCAACGAGGACGGCGAC CTGGTGCAGGACCCGCGGGCGCTGCCCATCGCGGAGAAAGTCGGCCAGCTTTTCCGCACG GCCATGCCGCTTATGGAGGGCGAAACCCGTGTCTTGTCGATCCCTTACGACCATTACGTTCTG CCGGGATACCTGTATCTGCCGCCGAAGAGCAGACGGATCCCGGGGCGCAAGAAGGTACCG ATCCTAGTGAATACCGGCGGCGCCGATTCGTGCCAAGAGGAGCTGTTTTACCTTAACCCGGC GGCCGGACCTGGCATGGGCTATGCCGTTCTCACGTTCGAGGGGCCGGGCCAGGGGATCAT GCTGAGGAAATACGAGCTTGAGATGCGTCCTGATTGGGAGGTCGTCACCAGCTCCGTTATCGA CTGCCTTGAGGTGTACAGCGCACAACATCCCGAGCTAGAGCTGGATCTGAGCTGTATCGCCGT TTCGGGCGCGTCGATGGGAGGATACTACGCTCTTCGAGCGGCATCCGACCCCAGAGTGAAA GCCTGTGTTTCGATTGTAAGTTTGCCCTTGTTGTATTGGACTTTCGAACGCTTGTTCTCGTAACTAA TTGCACAGGACCCCTTCTACGACATGTGGGATTTCGGCACAGCACACGTATCCCCGCTCTTCA TCTCGGCTTGGACCTCGGGTATCATCAGCTCCGGCTTTGTGGATAAGCTGATGGCCGTGATATC TCGTTTGTCGTTCCAGCTCAAGTGGGAGATATCGGTAGCCGGAACCTTTTTCGGCCTGTCGTCT CCGGCTCAAATACTGCTGAACATGAAGAAATACACATTGCATGGCAACGGGCGGGACGAAAAA GACGTCAGCTTTTTGTCCAAGGTAGAGTGTCCGGTCTTGCTCTCCGGCGCGGGCAAGTCGCTC TATCTAGACGTGGACAACCATACCAAGAGATGTTACGACGCGCTCACATCCGTAGCGCTGGAA GACAAGGAGATCTGGGTGCCCGAGAGTGAGGGCCAGGGCAGTCTCCAGGCGAAGATGGGTG CTTTTGCGTTGTGTAATCAGAAGACGTATCGGTTTTTGGACAAGGCTTTTGGGATAAAGAGAGAG CCCCTGAAGGGCGGCCTGTAG |
| XsTF | MHRVSSRRSACDRCRSHKLRCVR LDQNPNSASATGGPDTMMPCQR CLKAGTNCVRTAQVTGKSTGEN | ATGCATAGGGTGAGCAGCCGACGGAGTGCTTGCGACCGCTGTAGGAGCCATAAACTTCGGTG CGTAAGGCTTGACCAGAACCCGAACTCAGCCTCGGCGACCGGCGGCCCAGACACCATGAT GCCGTGCCAGCGATGTCTAAAGGCTGGCACTAACTGTGTTCGTACCGCACAAGTGACCGGAA |

| | | |
|---|---|---|
| | DPARRVSTCSGRSATSYQQRPPAA DQQLKDFPIDISAQASPQVQRKSR GPGPGKSASPQWQMFAQPSSRPR LESPKPPHLASHDSRPDHRLPEPP TSLITPPASDKAHRSSDASYFDMDE MGLESRGFGPISSLLHDNTGTSQM LDYSTEPLVPSSFFDTAANVMAGQS LHEQQMLLPGSTTDDCLQRLSQLS SKLLMDFGKASPPNMEDVMPFLSS PIGTSTSRQDGTNGTMNSYLSSTVG KLFESLQVYLETVERLRPYPNSSSA SECSYSDQWDEPELVSTTDDNQLY QGTTLADHIHDFPNARQSAGERAG THPFDNPRPFDMPATLTILTCYTWL LKGYEMVLSGIQDTLASQDRLQGL KPLPSIFHGSRIGNFALEDHPDMQI EIVIHIGSQLLHRIEGVLGIHVVSER SSAQDGQGDRREILDTNSAAALLG IWFAKGGQGENHSGDPCGGRRIQ INHTVENIRRLLREYWRDFRGK | AGTCTACCGGAGAAAACGATCCCGCACGCCGTGTCTCGACCTGTTCCGGCCGGTCTGCTACA TCGTATCAACAACGGCCCCCGGCCGCCGACCAGCAACTCAAAGACTTCCCTATTGACATATC CGCACAGGCATCCCCGCAGGTGCAGAGAAAGAGTCGCGGCCCTGGACCAGGCAAAAGTGC ATCACCCCAATGGCAAATGTTCGCTCAACCCTCTTCAAGACCCCGATTAGAATCTCCGAAGCC GCCTCATCTGGCAAGTCACGATTCGCGGCCTGACCATAGATTACCCGAGCCCCCTACCTCGC TCATTACGCCGCCCGCCAGCGATAAAGCCCACAGATCGAGCGACGCTTCGTACTTCGACATG GATGAAATGGGTTTAGAGAGTCGGGGATTTGGGCCGATTTCCAGCCTGTTGCATGATAACACAG GGACATCCCAGATGCTGGATTACTCGACAGAACCACTTGTACCATCAAGTTTCTTTGACACGGC GGCAAACGTCATGGCAGGTCAGAGCCTGCACGAGCAGCAAATGCTTCTGCCTGGTTCCACTA CGGACGACTGCTTGCAACGTCTCTCTCAGCTCAGCTCGAAGCTTTTGATGGATTTTGGTAAAGC CAGTCCTCCAAACATGGAAGACGTGATGCCTTTCTTATCGTCTCCCATCGGCACGTCCACCTC GCGCCAGGACGGTACTAATGGTACGATGAATAGCTATCTCAGCAGTACGGTTGGTAAGCTGTTT GAGAGTTTGCAGGTCTATTTAGAGACCGTCGAGCGCTTACGGCCGTACCCGAATTCATCATCG GCCTCGGAATGTTCCTATTCGGATCAGTGGGATGAGCCCGAACTCGTTAGCACAACAGATGATA ACCAGTTATACCAGGGCACTACGTTAGCTGATCATATCCACGATTTTCCAAACGCGCGTCAAAG TGCGGGAGAACGCGCAGGTACACATCCATTCGACAACCCTAGGCCGTTCGACATGCCGGCA ACCTTGACGATACTTACTTGCTATACCTGGCTATTGAAGGGCTATGAGATGGTTCTCTCGGGAATA CAAGACACACTTGCATCGCAAGATCGCTTGCAGGGCTTGAAACCGCTGCCGTCGATTTTTCAT GGCTCGAGAATCGGCAATTTTGCTCTGGAGGACCATCCAGACATGCAGATCGAGATCGTGATC CATATCGGCTCGCAGTTGCTGCATCGCATCGAGGGAGTTCTAGGAATCCACGTGGTATCGGAA CGCAGCAGTGCCCAAGATGGCCAGGGCGACAGGCGTGAGATCCTCGATACAAACTCGGCAG CTGCTTTGCTGGGCATCTGGTTCGCAAAGGGTGGCCAGGGAGAAAATCACTCAGGAGACCCC TGTGGCGGCAGGCGGATACAGATTAACCACACCGTAGAGAACATCCGAAGACTTCTGAGAGA GTACTGGAGAGACTTTCGGGGAAAATAG |
| XsMFS | MPKCSVGLRACRVRVHPIDRLLQL PGVLNVSDAWSAGLGWFGVVRAA RTTAADHLTKSQLPVADARLAGACL HGLAGLCPLPSRSAFPPFPPSPPAP RALVPVTLPRNADAQANPLLFFLLF LSAPSASRFLSSSYPQPPGLIIWTST YKMESQPTAASVHAPADSVRSAAD SVHDANPPASDPKTSKGPLFWLTFI GLVITAFLSALEGSIVSTALPSIARAL DASQNYIWVVNVYYLTSAAVQPFY | ATGCCCAAGTGTACGTACCGGTAGTTGCAAATCTCTGGTGAATACCAATAATTACGATGTCTGTAG GTTCTGTTGGTCTTCGTGCATGTCGTGTACGCGTCCATCCTATCGATAGGCTCCTACAGCTTCCT GGCGTGCTTAATGTCTCTGACGCCTGGTCAGCAGGTCTTGGCTGGTTCGGCGTGGTCCGTGTG GCAAGAGATATTAGAGTCCCGCTCTGAATGGCGGCATCGAAGGCGTGGCCCTGTGGACGCTG TACTGTAATCCGCGGTGACAGGCGCAGCTCGCACCACGTAAGTGGCTACAGTGGACTTGTGC CGTCGCTATAAGGGACAGTGCAGCTGACACGCAGCAGCGCGGCAGATCATCTCACCAAGTCA CAGTTACCCGTGGCCGATGCGCGCCTCGCGGGTGCGTGCCTCCACGGTACGTACCGCCATT TACGGGTGCCTGTTGTCCACGTAAGGCAGGGTCGTATTATTAGGTGGCAGGCGCAATCAACCT CGCCTCATAGAGTACCGGTACGGTAGATAGTACCTAGACTGTGCAACTGCCCTCTCTCTCCTCT ACGAGCCTATCTTAGCTAACCAGGCCTCGCCGGTCTCTGCCCACTGCCCAGTCGCTCGGCAT |

| | | |
|---|---|---|
| | GQLADLWGRRWLTIGTVAIFTVGSA<br>ICGSATSTNILIGGRTIQGLGAAGIN<br>VLVELILCDLLPLRERGQFFGILFLFI<br>ILGSVIGPFLGGILVDRVSWRWAFYI<br>NVPFGGASTILLFFVLRLKHTAPGST<br>IEKIKKIDFAGNLLISASVASVLYALTY<br>GGTRYNWTNAGVIVSLVLGLLGHG<br>LFLAFESSRWCANPVMPMALFKN<br>RTSTAAYVATFLQTIVSFWALYFLPLY<br>FQSVQLVSATRSGVMLLPFSVFYAL<br>AAFVGGGLTTKLGRYRIIHFIGFAIM<br>TIGMGTFTIFNRNTTLAVLVVLEVIFA<br>FGIGVVTPNLLTAIQAALPDDLNAA<br>STGTFAFVRSIGTIWGVSIPAAIFNN<br>RFDQLLGELSDQKAIAALAHGGAY<br>ESASSVFVDSFPPRVRDVIISIYERS<br>LMQVWQIGIVFAGLGFLVIAFEKDL<br>KLRKEKKTQDVALEDRPPSTGRDIT<br>SQDRMVENGGSN | TCCCCCCTTTCCCCCCTTCCCCCCCAGCCCCAAGGGCCCTCGTACCCGTAACCCTACCCC<br>GTAACGCGGATGCCCAAGCGAACCCCCTCCTGTTTTTTCTTTTGTTTTTAAGTGCTCCCTCTGCT<br>TCCCGCTTCCTCTCATCGTCATACCCGCAACCTCCAGGACTCATCATCTGGACATCGACGTACA<br>AGATGGAGTCCCAGCCGACGGCAGCCTCTGTCCACGCACCAGCAGACTCGGTCCGGAGTG<br>CCGCAGACTCTGTTCACGATGCAAACCCCCCGGCATCCGACCCAAAGACATCCAAGGGCCC<br>GCTGTTCTGGCTTACCTTTATCGGCCTTGTTATTACCGCTTTTTTAAGCGCCTTGGAGGGCTCCAT<br>CGTGTCCACAGCGCTCCCGTCCATTGCACGTGCCCTCGACGCATCCCAGAACTATATCTGGG<br>TAGTCAACGTTTACTATCTGACCAGGTATGTGTCATGGCCGCGGCCCCGGTTCTTCGCTTCGCC<br>ACCCGTCTAATTCTCTATCGTCCCCAGTGCTGCCGTCCAACCATTTTACGGCCAGTTGGCTGAT<br>CTATGGGGCCGACGATGGCTCACCATCGGCACCGTCGCCATCTTCACCGTCGGCAGCGCCA<br>TCTGCGGTAGCGCCACTTCAACTAACATCCTCATCGGCGGTCGGACAATCCAGGGTTTGGGTG<br>CAGCCGGCATCAATGGTGAGTGTTACCTCGACATTGGCCCTATAAATCTACGTCCTAACGCCAG<br>CAAATTGCAGTACTCGTCGAATTGATTCTCTGTGACCTGCTGCCCTTGCGGGAGCGCGGACAG<br>TTCTTTGGCATTCTATTCCTTTTCATCATCCTTGGATCCGTCATCGGCCCGTTCTTGGGCGGTATT<br>CTCGTTGATCGAGTTTCATGGAGATGGGCATTCTACATTAACGTGCCATTTGGCGGGGCTTCCA<br>CGATTCTTCTGTTCTTCGTTCTTCGTCTAAAACACACCGCACCGGGCAGTACGATCGAGAAGATA<br>AAGAAAATTGATTTCGCCGGTAACTTGCTAATTTCCGCATCGGTCGCCTCGGTCTTGTATGCCCT<br>CACGTACGGGGGTACTCGATACAATTGGACGAATGCAGGCGTCATCGTGAGTCTTGTCTTGGG<br>GCTCCTAGGCCACGGTCTATTCCTGGCCTTTGAATCTTCCCGCTGGTGCGCCAACCCTGTCAT<br>GCCAATGGCGCTGTTCAAAAACAGAACATCTACAGCTGCGTATGTCGCGACATTTCTACAGACC<br>ATTGTCTCGTTCTGGGCGCTCTATTTCCTCCCGCTCTATTTCCAATCGGTACAGCTAGTCTCGGC<br>CACACGGTCCGGCGTCATGCTCCTGCCATTCTCCGTCTTCTATGCCCTTGCCGCATTTGTAGGA<br>GGCGGGTTGACAACAAAATTGGGACGCTATCGAATTATCCATTTTATTGGCTTCGCCATCATGAC<br>CATTGGTATGGGCACGTTCACAATTTTCAACAGGAACACGACTCTAGCGGTCTTGGTTGTGCTG<br>GAAGTCATTTTCGCCTTTGGCATCGGCGTTGTTACCCCTAACCTCCTGACTGCCATTCAAGCTG<br>CCTTGCCGGATGATTTGAATGCCGCCAGCACTGGCACTTTTGCATTTGTGAGAAGTATAGGGAC<br>TATCTGGGGAGTCAGTATCCCCGCGGCTATATTCAATAACCGGTTTGACCAGCTTCTCGGAGAG<br>CTCTCCGACCAGAAGGCGATAGCAGCCCTAGCGCATGGGGGTGCCTACGAATCTGCGTCGA<br>GCGTGTTTGTTGATTCCTTCCCACCCCGTGTTCGAGATGTGATTATCAGCATTTACGAGCGGAGT<br>CTGATGCAAGTCTGGCAAATTGGAATTGTCTTTGCGGGTCTTGGGTTCCTCGTCATCGCTTTTGA<br>GAAGGATTTGAAACTGCGAAAAGAGAAGAAGACCCAGGATGTTGCACTGGAAGATCGCCCCC<br>CAAGTACGGGTCGTGACATCACTTCTCAAGATCGTATGGTAGAGAATGGAGGGTCAAACTGA |
| XsOAcT | MQFVTFRDWFGLTRYPIIGTDEVYP<br>LSFLDNLGAERGGVLSETLLFNHVL | ATGCAATTCGTAACGTTTCGTGATTGGTTCGGGTTGACCCGTTACCCAATAATAGGAACCGATGA<br>AGTCTACCCTCTATCTTTCCTCGACAACCTGGGCGCCGAGCGCGGTGGCGTCCTAAGCGAGA |

| | | |
|---|---|---|
| | DANKLYNGLTRLIQHGDWRKLGG<br>RLRYRSDGYLEVHVPREFTEDRPAV<br>QFTSRAFDIAIEDHKLGSTLPKSAD<br>GPRLQPGPRSFDHFNPIPDIPDKL<br>RDYTSSDRPILKLHVTSFNNATLVTL<br>IWPHAIAGGRGIKEIILAWSKALQS<br>DKDIPELLCARSNILEDVGTDKDNP<br>PPYHLSPSQVKGWGFVRLTLNLLW<br>SVFRHPQVESRTICLPGEYVSQLKE<br>TCLEDLRAVHQGEKVPFLSENDVL<br>EAWGTRFVAQARGGERPALVVTSL<br>DIKGRLDVPWGTEGVHVQNTGNN<br>VYTSAEPEVLLRRPLGELAQLIRKSI<br>QEAATDEQIRAQIRLFRANTRSLPV<br>FGHPNAHLMSFSNWTKFGIFEVAD<br>FSPAIVTQTSPTVSDGIPIGKPAYMH<br>CQALGENRFLRNCFNITGKDWDG<br>NYWMTTFLYPEDWAKLEEYMEQTR<br>QRIKEAKMR | CGCTCCTATTTAACCATGTGCTAGATGCCAATAAGCTTTATAATGGACTGACAAGACTGATTCAAC<br>ACGGCGACTGGAGAAAACTGGGCGGCCGACTTCGATATCGGGTGCGTAACTGTTCTCCGTCG<br>ACGTGGTCTCGTCGAGATGCTGACTCGTCTCATACCACAAGTCCGACGGCTACCTTGAGGTCC<br>ACGTACCAAGGGAGTTCACCGAGGATCGTCCAGCTGTCCAGTTCACCTCCAGAGCCTTCGAC<br>ATCGCCATTGAAGACCACAAGCTTGGCAGCACATTACCCAAGTCGGCAGACGGTCCTCGCTTA<br>CAACCAGGGCCAAGGTCATTCGACCATTTCAACCCCATTCCAGATATACCTGACAAACTGCGG<br>GACTACACATCCAGCGACCGTCCTATCCTTAAGCTGCACGTCACATCCTTCAATAATGCGACCC<br>TCGTCACTCTTATCTGGCCACATGCCATCGCCGGCGGCCGTGGCATAAAGGAGATCATCCTAG<br>CCTGGTCCAAGGCGCTCCAAAGCGACAAGGATATACCCGAGTTACTATGCGCTCGCAGCAAC<br>ATCCTAGAAGACGTCGGAACTGACAAGGACAACCCGCCACCGTACCATCTCAGCCCGAGCC<br>AAGTCAAAGGCTGGGGCTTTGTCAGACTCACCCTCAATCTGTTGTGGAGCGTATTCAGACATCC<br>ACAAGTCGAGTCCCGCACGATATGCCTGCCTGGTGAGTACGTATCGCAACTGAAAGAAACCTG<br>TCTAGAGGATCTCCGCGCCGTTCACCAAGGAGAAAAGGTGCCCTTCCTAAGCGAGAACGACG<br>TCCTAGAGGCCTGGGGTACAAGATTCGTAGCCCAAGCGCGCGGCGGAGAACGACCAGCGCT<br>AGTCGTCACCTCCCTCGATATCAAGGGCCGACTAGACGTTCCCTGGGGCACCGAAGGAGTAC<br>ATGTGCAGAACACAGGCAACAACGTGTATACCTCAGCCGAGCCCGAAGTCCTGCTGAGGAGG<br>CCGCTAGGGGAGCTAGCACAACTAATACGCAAGAGCATCCAGGAGGCAGCCACGGACGAG<br>CAGATCCGAGCCCAGATACGCCTCTTCAGAGCGAACACAAGGAGCCTACCAGTCTTCGGACA<br>CCCAAACGCCCACCTGATGAGTTTCTCCAACTGGACCAAGTTCGGCATATTCGAAGTTGCTGA<br>CTTCAGCCCAGCCATCGTAACTCAGACGTCGCCAACCGTCTCCGATGGCATACCTATCGGGA<br>AACCTGCCTATATGCACTGCCAAGCGTTGGGAGAAAACCGATTCCTCAGGAATTGTTTCAACAT<br>CACCGGAAAAGACTGGGATGGCAACTACTGGATGACGACATTCCTTTATCCCGAGGACTGGGC<br>GAAGCTTGAAGAATACATGGAGCAAACCCGTCAACGGATCAAAGAGGCAAAGATGCGATAA |

**Table S4**: Putative cytochalasan BGC identified in *Pseudoxylaria* sp. X802 (NCBI genome accession number = JAJFDH000000000).

| Protein Name | Synthetic DNA Sequence | Protein Sequence |
|---|---|---|
| CHGG_ 01243 | ATGTCCATGGTTGGCGCGTGTGTGCTGATTGGGACGGTGGTGTCGGTTCTGGCTTGGACCACGTCAAAGTT<br>GTTCTTTCGACCCTCCACTCCATCACTCGAAGATCTAGGAATTCCTTTGCTGGGCAAATCTCGGGGGGGTA<br>AATTTGACTTCCGCCAGATGATGCGAGACTGTGCACAACAGGCAAGTAATAAGCAGGTTGCGGGCTCCCTT<br>GCGGCCCCGGGCTCCCCTGACTGACAGCAAGCTCACCAGCTTCCGGACCGGCCCTACAGAATACGGG<br>CCTTCGGGACCGAGTACGTGGTCTTCCCATCCCGGTACTTCAACGATATCAAGCGTATTCCGGCAAGAGA<br>CGCGTCGGCTTATGAATTCTTCCGCCATGCCTTTCACACCAACTGGTCCGGGATTCCCCGGCACAGCGA<br>GGCCATGATGAAGTGAGTATATGTACATACTTTTGGGCGACCCCAGTTGCGATCACTGTGCTAACCCGCCC<br>CTTCAGGGCCGTAGCGGTCGATGCCACCCGTGCAATACCGTCACTCATCCACGACCGGCAAAACGACT<br>GCGCGGCAGCATGTGATGCTTCCATTGGCGAGTGTCCGGACTGGACCGAAATCACCATGTTTCCGGCGC<br>TGCAGAAGATTGTCGTATCGACGAATGCGAGCCCCCTTGTTGGAAGGGAGCTGGCGTCCAATTCCACTTG<br>GATATGGCACGTCGAGCGGCTGCCAATGCTGCTGGGCATTCCCACTGTGATTCTCAGCCTGACCCCGGC<br>GGTCTTGCGGCTATTGATCAAACCGCTCCTGTTCGTCCCCATCCGGTACTCGAGCTTTGTGCTAACGCGG<br>CTCATCACGCCGGTGCTCAAGGAAGACATGTTGGAGTTCGAGTCGACCGCAGACAAGAAGTCGCCAGCG<br>GGCCCCAAAGCAAAAGGAAAGCTGGCATTGACGAGCTGGCTTTTGAGCCGCTACCCAGCATCGCTGAAG<br>GACAGAATGTCCCAGCTCATTCGCGACTACCTCGCCATTACGTTCGAGTCCACCCCATCGACGTCGGGG<br>GTGCTTTTTTACATCTTGATAGAGCTGGCAGCAGCCCCGGAGCTAGCGGAAGCGGTGCGCCGGGAGCTA<br>AGAGAGGTCGCTCCCAACGGCGAGTTGCCGTCCACCCACCTCAACGAGCTCAAGGTTATGGATAGCGTC<br>ATGAGGGAGTCAGCCCGAGTCAACCCTTTCAGCCATTGTAAGCCATTCAGCTGTAAAACTCGCATGAACAC<br>AGGCATTAACTCCCAATTTGCAGTGGTCCTCTATCGAAAGCTTCTGCGCCCCTTGAAGCTTGAAGGTTGCC<br>CCGAGTTGCCGGCTGGTTGCTTCATCTGCGTAGACGCCCACCACATTGACTTCTCGCCGCAGCTTTGGGA<br>AAACCCGGAGCGGTTCGACGGGCTTCGGCACTACCGGGCTCGCCAGAAGCCCGAAAACGGCAACCG<br>GTTCAAGTTTGCCAACCTGGGATCCGACGCGCCCGGCTGGGGAGACGGCCCGCAGGCCTGCCCGGG<br>GAGGATGTTCGCCGACAACACCATCAAGATCATCCTCGCACACATTCTGACGCACTACGACCTTGAGCTG<br>CCCCCGGGGCAGGGGAAGCCCGAGAAGGGATCGATGCCCAATGGGTCCATGAGTCCGGATACCAAGG<br>CTAAGGTCTTGTTTCGGTCGAGGAAGCTGTAG | MSMVGACVLIGTVVSVLAWTT<br>SKLFFRPSTPSLEDLGIPLLGKS<br>RGGKFDFRQMMRDCAQQLP<br>DRPYRIRAFGTEYVVFPSRYFN<br>DIKRIPARDASAYEFFRHAFHTN<br>WSGIPRHSEAMMKAVAVDATR<br>AIPSLIHDRQNDCAAACDASIG<br>ECPDWTEITMFPALQKIVVSTN<br>ASPLVGRELASNSTWIWHVER<br>LPMLLGIPTVILSLTPAVLRLLIKP<br>LLFVPIRYSSFVLTRLITPVLKED<br>MLEFESTADKKSPAGPKAKGKL<br>ALTSWLLSRYPASLKDRMSQLI<br>RDYLAITFESTPSTSGVLFYILIEL<br>AAAPELAEAVRRELREVAPNGE<br>LPSTHLNELKVMDSVMRESAR<br>VNPFSHLVLYRKLLRPLKLEGC<br>PELPAGCFICVDAHHIDFSPQL<br>WENPERFDGLRHYRARQKPE<br>NGNRFKFANLGSDAPGWGDG<br>PQACPGRMFADNTIKIILAHILT<br>HYDLELPPGQGKPEKGSMPN<br>GSMSPDTKAKVLFRSRKL |
| CcsD | ATGGTGAACGAGATAAGCCCAAGAACCCTCGTGCTGTTGGCGGTAACCTGTTCGCTGCTGGTGCTCTACTT<br>CTCAACGCGAGAGAGGCAACCGGTCATGCCGCGAGAGCTCCCGGTAGTCAAAGCAAAGAGTTATCACTT<br>TGAAGATATAATCGTCGAAGGCAGAAAAAAGGTTCCTTTTATCTACAATCGCTTGCGACAAATCTGTGGCTGA<br>CCAACGGCAGTATCCGGACAGACCTTATCTCGCCGTGAACAATCGACACAGCTTCGTTGTCTATCCGCCA<br>AGCTGCTTTGACGAGATTAAACGCCTTCCAGAGCACAGCGCATCAGCGAAAGATTTCTTCCACACGATGAA | MVNEISPRTLVLLAVTCSLLVLY<br>FSTRERQPVMPRELPVVKAKSY<br>HFEDIIVEGRKKYPDRPYLAVN<br>NRHSFVVYPPSCFDEIKRLPEH<br>SASAKDFFHTMNAGDWTYVG |

| | | |
|---|---|---|
| | TGCAGGAGATTGGACATACGTTGGCCACGAGACCACGCCCCTTCTGAAGACAATCATCGCAGACCTCAC<br>CCGTGTCATCCCTGCGCGTGTGAACAAGCGGCAACAGGACACCCGCATGGCGTTTGAATCGATTGTGGG<br>ATACGCTCCAGAGTGGAAAGAAATTGGGTTGCTGATGACCACGTTCGAAATTGTCGCCAAGATCAACGCCT<br>GTGCTTTTGTTGGGAGAGAGCTTGGGACGAACAACAAATGGGTAAAAGCGGTGATGCAGTCGCCTCTGGT<br>CATTCATGTTGCAGTGCTGATCATGAACGCGTGTCCTGCTCTGCTCCGGCCACTCCTGGCGCCCCTGGCC<br>TTCCTGCCCACGAAAATGAACCAATGGGATATGCGCCGGCTGCTGACTCCCATGTTGCAGGAGGATATGG<br>CGATATTCAAAGAGACCAAGGATCGATCTGAATTGTTGCGTCCAAAGCAGAATGGAAAGATCCCTTTGACAG<br>CGATGTTGCTATCGCGGTACAAGCAAGGCGAGGCCACTATTAGGCAGCTGATCGTGGACTATATCCTCATC<br>AGCTTCGACTCGACGCCATCGACCGCGTCGGCGCTCTATCATGTCATTTGCGAGCTCGCGGCACACCCG<br>GAGGCGGCAGATGTCTTGCGCCAGGAGCTCGATGAAGTTATGGTAGATGGAAAGCTTCCTCAGACGCATTT<br>GCAGGAGCTCAAGAGAATGGACAGCTTTCTGCGCGAGTCCTTTCGTTTACACCCCGTCAGCTTGTGTGAGT<br>ATCTGCACCTGCACGCAATGTTGGTATTGAGATCTCACGCTAATCTCTGAAATAGTCAGTCTTCAGCGTGTCC<br>TGGCTAAGCCCGTGAAGCTGAGTGTCGGCCCCACGATTCCTGCCGGAGCCATCATCGCCGTGGATGCA<br>GCGGCCATCAATCGCTCTCCCAGTCTCTGGAAGGACCCGGACGAATTCGATATGAATCGATTCCATGACTT<br>GCGCCAGGTACCGGGAAATGAGAACAAGTTCCACCTGCTGAACACTGGCTCGGATTCTCCAGGATGGGG<br>GGATGGCACGCAAGCCTGCCCTGGCCGCTTTTTCGCAAATAGCACTCTGAAGATTGCATTCGCATATATTC<br>TTCAAAACTATGATGTTAAGAGGAAGGAGGGATCTTCTCCACCCAAGATGACTCCGTTGGCTAATGGGACAT<br>GGGCACCGGACGATAAAGCAGCGGCTCTGTTCAAGTCTCGAAACTAA | HETTPLLKTIIADLTRVIPARVNK<br>RQQDTRMAFESIVGYAPEWKEI<br>GLLMTTFEIVAKINACAFVGREL<br>GTNNKWVKAVMQSPLVIHVAV<br>LIMNACPALLRPLLAPLAFLPTK<br>MNQWDMRRLLTPMLQEDMAI<br>FKETKDRSELLRPKQNGKIPLTA<br>MLLSRYKQGEATIRQLIVDYILIS<br>FDSTPSTASALYHVICELAAHPE<br>AADVLRQELDEVMVDGKLPQT<br>HLQELKRMDSFLRESFRLHPV<br>SLFSLQRVLAKPVKLSVGPTIPA<br>GAIIAVDAAAINRSPSLWKDPD<br>EFDMNRFHDLRQVPGNENKF<br>HLLNTGSDSPGWGDGTQACP<br>GRFFANSTLKIAFAYILQNYDVK<br>RKEGSSPPKMTPLANGTWAPD<br>DKAAALFKSRN |
| CcsD (revised sequence with missing N-terminal sequence highlighted in blue) | ATGTGGAGCCTACAGTCCAACATCATCGAGCCATACCAAGTGCTACGGCAGTCGCTTGAGCCGTTGAAAC<br>TCTCAAGATGGCAAATGACGAAATTGATAGCGAGGAGCATGGTGAACGAGATAAGCCCAAGAACCCTCGT<br>GCTGTTGGCGGTAACCTGTTCGCTGCTGGTGCTCTACTTCTCAACGCGAGAGAGGCAACCGGTCATGCCG<br>CGAGAGCTCCCGGTAGTCAAAGCAAAGAGTTATCACTTTGAAGATATAATCGTCGAAGGCAGAAAAAAGGTT<br>CCTTTTATCTACAATCGCTTGCGACAAATCTGTGGCTGACCAACGGCAGTATCCGGACAGACCTTATCTCG<br>CCGTGAACAATCGACACAGCTTCGTTGTCTATCCGCCAAGCTGCTTTGACGAGATTAAACGCCTTCCAGAG<br>CACAGCGCATCAGCGAAAGATTTCTTCCACACGATGAATGCAGGAGATTGGACATACGTTGGCCACGAGA<br>CCACGCCCCTTCTGAAGACAATCATCGCAGACCTCACCCGTGTCATCCCTGCGCGTGTGAACAAGCGGC<br>AACAGGACACCCGCATGGCGTTTGAATCGATTGTGGGATACGCTCCAGAGTGGAAAGAAATTGGGTTGCT<br>GATGACCACGTTCGAAATTGTCGCCAAGATCAACGCCTGTGCTTTTGTTGGGAGAGAGCTTGGGACGAACA<br>ACAAATGGGTAAAAGCGGTGATGCAGTCGCCTCTGGTCATTCATGTTGCAGTGCTGATCATGAACGCGTGT<br>CCTGCTCTGCTCCGGCCACTCCTGGCGCCCCTGGCCTTCCTGCCCACGAAAATGAACCAATGGGATATG<br>CGCCGGCTGCTGACTCCCATGTTGCAGGAGGATATGGCGATATTCAAAGAGACCAAGGATCGATCTGAATT<br>GTTGCGTCCAAAGCAGAATGGAAAGATCCCTTTGACAGCGATGTTGCTATCGCGGTACAAGCAAGGCGAG<br>GCCACTATTAGGCAGCTGATCGTGGACTATATCCTCATCAGCTTCGACTCGACGCCATCGACCGCGTCGG<br>CGCTCTATCATGTCATTTGCGAGCTCGCGGCACACCCGGAGGCGGCAGATGTCTTGCGCCAGGAGCTCG | MWSLQSNIIEPYQVLRQSLEPL<br>KLSRWQMTKLIARSMVNEISPR<br>TLVLLAVTCSLLVLYFSTRERQP<br>VMPRELPVVKAKSYHFEDIIVE<br>GRKKYPDRPYLAVNNRHSFVV<br>YPPSCFDEIKRLPEHSASAKDF<br>FHTMNAGDWTYVGHETTPLLK<br>TIIADLTRVIPARVNKRQQDTRM<br>AFESIVGYAPEWKEIGLLMTTFE<br>IVAKINACAFVGRELGTNNKWV<br>KAVMQSPLVIHVAVLIMNACPA<br>LLRPLLAPLAFLPTKMNQWDM<br>RRLLTPMLQEDMAIFKETKDRS<br>ELLRPKQNGKIPLTAMLLSRYK<br>QGEATIRQLIVDYILISFDSTPST<br>ASALYHVICELAAHPEAADVLR |

| | | |
|---|---|---|
| | ATGAAGTTATGGTAGATGGAAAGCTTCCTCAGACGCATTTGCAGGAGCTCAAGAGAATGGACAGCTTTCTGCGCGAGTCCTTTCGTTTACACCCCGTCAGCTTGTGTGAGTATCTGCACCTGCACGCAATGTTGGTATTGAGATCTCACGCTAATCTCTGAAATAGTCAGTCTTCAGCGTGTCCTGGCTAAGCCCGTGAAGCTGAGTGTCGGCCCCACGATTCCTGCCGGAGCCATCATCGCCGTGGATGCAGCGGCCATCAATCGCTCTCCCAGTCTCTGGAAGGACCCGGACGAATTCGATATGAATCGATTCCATGACTTGCGCCAGGTACCGGGAAATGAGAACAAGTTCCACCTGCTGAACACTGGCTCGGATTCTCCAGGATGGGGGATGGCACGCAAGCCTGCCCTGGCCGCTTTTTCGCAAATAGCACTCTGAAGATTGCATTCGCATATATTCTTCAAAACTATGATGTTAAGAGGAAGGAGGGATCTTCTCCACCCAAGATGACTCCGTTGGCTAATGGGACATGGGCACCGGACGATAAAGCAGCGGCTCTGTTCAAGTCTCGAAACTAA | QELDEVMVDGKLPQTHLQELKRMDSFLRESFRLHPVSLFSLQRVLAKPVKLSVGPTIPAGAIIAVDAAAINRSPSLWKDPDEFDMNRFHDLRQVPGNENKFHLLNTGSDSPGWGDGTQACPGRFFANSTLKIAFAYILQNYDVKRKEGSSPPKMTPLANGTWAPDDKAAALFKSRN |
| XsCYP2 (cDNA sequence) | ATGTTGTCAAGCATCCAAACGAATATTGTCGAGCCTTTTCTGGTGCTGCGCCAGAGTGTGGCACCTTTGAAACTGTCCAGATGGCAGCTCACGAAGCTTATGATCCGCACTGCACTCAATGGGCTGCCCGACGGAAGCCTCTTTTTCCTCCTAGCGGCCTTGGCAGCGACTATTGCTGTGTACTACCTTATATTCAACAATACAAACACACGCGTACAAGTGCCTCCCGGGCTCGCCGTGGTGAAGAGGGATGACATGCATTTCCTTGATATCATCGATGAAGGACGGAAGCTGTACCCAGGTCAACCCTTCTTGGCGGTAAACAGGCGGCATAGTTTTGTCATATACCCGCCTCAATGCTTTGATGAGATCAAGCGTCTTCCGGAGCACACTGCGTCCGCGAGGGCATTCTTCCACGCTACCAACTATGGCCACTGGAGCCACGTTGGCACTGAGACACCCGAACTCATCAAATCGGTCATTGCGGATTTGACCCGTTCGCTCCCTGCTCGAGTGCTCGCTCGTCAAGAAGACTGCCAGAACGCATTCGACGGGGTCCTTGGGCGCCGGCGCGATTGGAAGGAATTCCCTCTGATGATGACTACATTCGAAATCGTCACCCAGATCAATGCGTGTTCTTTTGTCGGAAGAAAACTAGGCACTAGCCGCGGCTGGGTCAAGTCTGTCATGATGTCGCCTATCTTCATTCACGTTGCTGTCACGCTCCTGGATGCCTGCCCATTCATCCTCCGGCCGCTCATGGCCCCGATTTACTTTTTCCCTACTATGAAGAACCGATGGGATATGAAGCGACTGCTCACGCCTATTCTGGAGGAGGACATAAAAGATTTCTATGCAGCGACGGACAAGAAAGAAATCTTGCGACCCCGGCCAGACGGCAAGATTCCCTTCACTGGGTTCCTCCTCTCACGCTACCGGACGGCCGAAGCAAGCATCAGACAGCTGATCTCAGACTACATCCTCATCAGCTTCGACTCAACCCCATCCACCGCATCGGCATTTTTCCATGCTCTCTGTGAGTTGGCATTGCATCCGGAAGCCGCCGACATTCTGCGAGAAGAACTAGACGAATATGTCGTTGACGGAAACCTCCCTGGGACTCATCTTCAGGAGCTAAGGAAAATGGACAGCTTCCTCCGCGAGTCCTTCCGGTTGCATCCTATTGGCATATTCAGCCTCCAACGCGTCGTTGAAAAGCCGATAAAACTCTCGGTGGGCCCGACTATCCCGCCTGGCACAATCATCGCTGTCGATGGACAAGCTATCAACCGGTCCCCGGATTTGTGGCCAAATCCGGATAGTTTCGACATGGACCGATTCTATAAGTTGCGGCAAAAACCCGGCAACGAAAATCGGTTTCATTTCCTGACCACTGGCTCCGACTCGCCGGGTTGGGGCGACGGGACACAGGCCTGCCCGGGTCGGTTCTTCGCGACAAGCACGTTAAAGATCGCGATGGCGCATTTTCTCAGGAATTATGACATCGAGATAAAGCCAGAGTGTCTGCCGCTGAAGCACACGCCGCTTTCAAATGGATCTTGGAAGCCCGACGATACGGCAATCGCACGTATTAGGGCCAGATGT | MLSSIQTNIVEPFLVLRQSVAPLKLSRWQLTKLMIRTALNGLPDGSLFFLLAALAATIAVYYLIFNNTNTRVQVPPGLAVVKRDDMHFLDIIDEGRKLYPGQPFLAVNRRHSFVIYPPQCFDEIKRLPEHTASARAFFHATNYGHWSHVGTETPELIKSVIADLTRSLPARVLARQEDCQNAFDGVLGRRRDWKEFPLMMTTFEIVTQINACSFVGRKLGTSRGWVKSVMMSPIFIHVAVTLLDACPFILRPLMAPIYFFPTMKNRWDMKRLLTPILEEDIKDFYAATDKKEILRPRPDGKIPFTGFLLSRYRTAEASIRQLISDYILISFDSTPSTASAFFHALCELALHPEADILREELDEYVVDGNLPGTHLQELRKMDSFLRESFRLHPIGIFSLQRVVEKPIKLSVGPTIPPGTIIAVDGQAINRSPDLWPNPDSFDMDRFYKLRQKPGNENRFHFLTTGSDSPGWGDGTQACPGRFFATSTLKIAMAHFLRNYDIEIKPECLPLKHTPLSNGSWKPDDTAIARIRARC |

| | | |
|---|---|---|
| AhCYP2 | ATGTCTGCCACCGTCGTTCTTACGGGGCTCGCAACACTGATGTTTGCATTCCTGCTTCTCCTGAAGTTCTGG<br>ACCAGGAAAAGCGCACTGGAAAGACTTGGGATCCCCACAGTTGGGGGCAGCAGACAACATACAAAGGA<br>CTTCGAGCGCTTGGTGGAAGAGGGGTACTACAAAGTACGTTTGTCCCGTGTTGGTCAATTCTTTCCAATCTC<br>GTCGCATCATTCTGACCGAGTATTGTCGACTGAACAACAGTATCCACACTCGCCTTATGCGATCAAAACTCA<br>AGGGTTGGAATACGTTGTTTTCCCACCGGAGGCATTCGATGAAATCAAGAAGCTCCCACCGCATATTGCAT<br>CGGCACAAGACTTTTTCGTCAAAACCTACTTTGGGCACTACACCACGGCGGGGACCGAGACTCCCGCGT<br>TGCTGAAGGCCATCAGCGTGGACCTAGCACGAAGCATGCCCCTGACGGTAGCGAGTCGGCAAGAGGAC<br>GCGAAAGCCGCGGCAGACGACGTCCTAGGATTCTGCCCGGAATGGAAAGAAGTCTCGCTCTTTCCCGC<br>CGTGACAAGAATGATCGCGATGACCAACGCCTGTAGCCTCGTGGGTAGGTCCCTCGCCCGGAGCGAGG<br>GGTGGATTGGACTAGTTAGTCGGTTCCCCTTCGAGGTAATGGCTGGTACCTTTGCCATCAGTGTGTTTCCGC<br>GCTTTCTTCAACCCGTCCTGGCGCCCCTCATCTTCCTCCCCGCTCTGGTCACCAAATGGCGCATGAAATG<br>GTCACTGCGCGGAGTGGTCCAACGAGACATGCAGGAGTACCAGTCCACGTCCGACAAGAAGATGCTGCT<br>GACGCTGAAGGAGGACGGCAAAGTCCCCTTCACCGCCGCGCTCATGACGCGGTACAAACCGAGCGAG<br>GCCACATTGTCTCAATTGCTGCATGACTACGTGACCGTCTCGTTCGAGTCCACCCCTTCGAGCACAGCGG<br>CCTTGTACCTGATTCTGATGGAACTGGCCACCCGGCCACAGCTGGTCGAGGTCCTGCGACAGGAACTAA<br>ACGAGGTCATGGTGGATGGGATGCTGCCAAGAACGCATCTGGCCGAACTCCGGAAGATGGACAGCGTGA<br>TGAGAGAATCGGCCCGAGCAAATCCATTCAGTCTCTGTAAGGAATCAGTCCCAGTCTTTGATCGAAGGTTG<br>GTCTAATTGAGACAGTGGCCCTGTACCGTTTGCTCCGAGTTCCCACCAAGCTATCAACGGGGCCAACCCT<br>TCCTGCCGGAACTCTGATCTGTGTGGATGTGCACCACATCCACACTTCGGAGGACCGTTGGACCAAGCCC<br>CAGGACTTCGACGGTCTGCGGTTCCATGAGATACGAAAGACCCCAGGGAAGGAGAACACGTATCAGTTTG<br>TCAGCACAGGTGCCGACTCGCCCGGTTGGGGGGACGGGGCCATGGCCTGTCCAGGGCGCATGTTCGC<br>CAACAGCACCCTCAAGATTGCATTGGCTCATTTGATCATGCACTATGACTTCCGATTTGCCGATGGCGAGG<br>GGAAGCCAATCAAAACCTCGCTTCCGAATGGCTCTTGGAACCCCGGCTTGAAGGTTCGGGTTTTGTTCAA<br>AAGTAGAAAGTGGGACGCGTGA | MSATVVLTGLATLMFAFLLLLKF<br>WTRKSALERLGIPTVGGSRQHT<br>KDFERLVEEGYYKVRLSRVGQF<br>FPISSHHSDRVLSTEQQYPHSP<br>YAIKTQGLEYVVFPPEAFDEIKK<br>LPPHIASAQDFFVKTYFGHYTT<br>AGTETPALLKAISVDLARSMPLT<br>VASRQEDAKAAADDVLGFCPE<br>WKEVSLFPAVTRMIAMTNACSL<br>VGRSLARSEGWIGLVSRFPFEV<br>MAGTFAISVFPRFLQPVLAPLIF<br>LPALVTKWRMKWSLRGVVQR<br>DMQEYQSTSDKKMLLTLKEDG<br>KVPFTAALMTRYKPSEATLSQLL<br>HDYVTVSFESTPSSTAALYLILM<br>ELATRPQLVEVLRQELNEVMV<br>DGMLPRTHLAELRKMDSVMR<br>ESARANPFSLLALYRLLRVPTKL<br>STGPTLPAGTLICVDVHHIHTSE<br>DRWTKPQDFDGLRFHEIRKTP<br>GKENTYQFVSTGADSPGWGD<br>GAMACPGRMFANSTLKIALAH<br>LIMHYDFRFADGEGKPIKTSLP<br>NGSWNPGLKVRVLFKSRKWD<br>A |

| | | |
|---|---|---|
| CsCYP1 | ATGTCATTCTCCATCGAATCCAAGATCCTCGAGCCGTACCTGGTTCTGAGGCAGAGCCTCGCGCCCCTCC<br>GGCTGTCCAGATGGCAAATGTTCAAGATAATCACCAGAACCCTCCTCTACGAGACGCTGCCCCGCGTCAT<br>CTTCTTTGGCACGATTCTGTGTCTTTTCCTCCTTGTCCTATCTCGCAAGGCTTGGGGTCGGCAGAAGAAGGA<br>GATCGACCACGGCCTCCCCATCGTCCAGAGAAACGACTACCACTTTGATACCATCATTGCCGAAGGCAAG<br>CAAAGGGTGAGAGACTCACAGACGTCCAACCGGCGGGCACTGGAGTCCATTGACTGACAGTCAACAGTA<br>CCCCGACCGCCCCTTCATGGCCATCAACAAACGGTACAGGTTCGTCGTCTACCCCACGAGCAGCTGGG<br>ACGAGTTCAAGCGCATCCCCGAGCAGACGGCGTCCATCATGGACTTCCAGCACGTCTGCAACTCGGGC<br>GACTGGAGCCTCATCGGCGGCGAGACCCACGAGCTCGTCAAGACCATCACGGCCGAGCTGACCCGCT<br>CGCTGCCGGCGCGCGTCCTCAACCGCCAGCAGGACGCCAAGATGACGTTTGACACCATCGTCGGCCA<br>CTGCCCCGAGGAGAAGGGCTTCAACCTGCTCATGACGTCGCTCGAAATCATCGCCAAGATCAACGCCTG<br>CACCTTTGTCGGCAGGGACCTCGGCCGCAACCAGCGTTGGGTAAAGTCCGTCATCTACTCGCCCCTGTT<br>CGTCTACATGGCCGTGACGCTCCTCAACGCCACGCCGGCCGTCCTGAGGCCGATTCTGCGCCCGCTGT<br>ACTTCCTGCCGACGCTGAGGAACTACTGGGGCATGCACAAGCTGCTGAAGCCGAAGCTCGACCGGGAG<br>ATTGCGGCCTTCAGGGCCGCGGGCCAGGACAAGCGCAAGCTGCTCGTGCCCAAGACGGACGAGGAC<br>CTCCCCTTTACGCACTTCTTGCTGTCGCGGTACACCGAGGCCGCGGCGACCATCAAGCAGCTCGTGACG<br>GACTACATCCAGGTCAGCTACACGTCGACGCCGACGACGGCCTCGGCGCTGTACCACGCGCTCTGGGA<br>GCTGGCGCAGCACCCCGAGGCCGCCGAGGTCATGCGGCGGGAGCTCGACGAGGTCATGGTCGACGG<br>CCAGCTCCCGCGGACGCACCTCCAGGAGCTGAAGCGCATGGACAGCTTCCTCCGCGAGTCGTTCCGG<br>CTGCACCCCATTACCCGCTGTAAGTCCTGGCCTGGCCCAAGTCATCGGCCTCTTTCCTTTGGTGTTGAAG<br>GGTTGGACTAATGCTTCCATGCTCTCGCTCTCGCAGTCACCCTCCAGCGCTACGTCAAGGAACCCTTCCA<br>GCTCTCGGACGGGGCCACGATCCCGCCCGGGATCATGGCCGTCTGCGACGCGCAGGAGATCAACCG<br>CTCGCCCGAGCTGTGGGAGGACCCGGACCGCTTCGACATGGACCGCTTCTACCGCCTGCGCGAGCTC<br>GAGGGCAACGACAACCGCTACCACTTTGTCACGATGAGCTCCAACTCGCCCGGCTGGGGCGACGGCA<br>CCCAGGCCTGCCCCGGCCGCTTCTTCGCCACGAGCACGCTCAAGATTGTCATGGCGCACGTCGTGACC<br>AACTACGACATCCGGCTGGGCAAGGTGGCGCCGCTGAAGAGCAGGCCCCTGGTGAACGGGTCGTACG<br>CGCCGGACGATACGGTGCAGATTCTGTTCAAGTCGAGGGCTGGCAAGTGA | MSFSIESKILEPYLVLRQSLAPL<br>RLSRWQMFKIITRTLLYETLPRVI<br>FFGTILCLFLLVLSRKAWGRQK<br>KEIDHGLPIVQRNDYHFDTIIAE<br>GKQRYPDRPFMAINKRYRFVV<br>YPTSSWDEFKRIPEQTASIMDF<br>QHVCNSGDWSLIGGETHELVK<br>TITAELTRSLPARVLNRQQDAK<br>MTFDTIVGHCPEEKGFNLLMTS<br>LEIIAKINACTFVGRDLGRNQR<br>WVKSVIYSPLFVYMAVTLLNAT<br>PAVLRPILRPLYFLPTLRNYWG<br>MHKLLKPKLDREIAAFRAAGQ<br>DKRKLLVPKTDEDLPFTHFLLS<br>RYTEAAATIKQLVTDYIQVSYTST<br>PTTASALYHALWELAQHPEAAE<br>VMRRELDEVMVDGQLPRTHL<br>QELKRMDSFLRESFRLHPITRF<br>TLQRYVKEPFQLSDGATIPPGI<br>MAVCDAQEINRSPELWEDPDR<br>FDMDRFYRLRELEGNDNRYHF<br>VTMSSNSPGWGDGTQACPGR<br>FFATSTLKIVMAHVVTNYDIRLG<br>KVAPLKSRPLVNGSYAPDDTVQ<br>ILFKSRAGK |
| CsCYP2<br>(cDNA) | ATGTTGCAGCTAGTTGGGTTCGGGCTCTCGCTCCTGACCGCGAAAGCCGCCGTCGTGCTGCTCGGGGG<br>CGGCGCCCTCGTAACCCTCGTCTACCTTTTGCACTTTTACAGCCTCTGGCCCTTTGGCAGCGACGTCGAG<br>CCCGCGTCCGCCGACCAGCTGCGGCACATACCGCTGCTGCGCTTCGACGGATCCAACAGCGTCGAGC<br>GCTACATGAACGAGACGCGGAGCCTCCTGAGGCTGGGCTACGAGAGGTACCTGCGCCGGGGCATCCC<br>CTTTCAGATCCGCCACCCCGTCGGCGAGCTGGGATCGCAGGTCCTGCTGCCCGTCAAGTACCTGGACG<br>AGGTCAAGAAGGCCCCGACGGACCTCTTCAGCTTCGAGGCCTTTTCCGAAAAGTCGTTTCTGCTGAACTA<br>CAGCCGCGCTCCGAGGCAGACCGAGGCCGCGGCGCACGTCGTCAGGGTCGACCTCAACAGAAACCT<br>GGGCCCCCTCGTGACGGACCTCTGGAACGAAGCCGCCCGCCACCTCAACGAGACCGTCGGCTCCGA | MLQLVGFGLSLLTAKAAVVLLG<br>GGALVTLVYLLHFYSLWPFGSD<br>VEPASADQLRHIPLLRFDGSNS<br>VERYMNETRSLLRLGYERYLRR<br>GIPFQIRHPVGELGSQVLLPVK<br>YLDEVKKAPTDLFSFEAFSEKSF<br>LLNYSRAPRQTEAAAHVVRVDL<br>NRNLGPLVTDLWNEAARHLNE |

| | | |
|---|---|---|
| | GTACAAGTCCGTCCCGGCGTACGATCTCGTCTGCGGCTTCGTCGCCCGCGTGGCCTCGGTCGCCCTCG<br>TCGGGGCCCCGCTGTGCCGCAACCCGGCGTGGCAGCGCATCGTCGTCGAGACGACCTTTGTCGCCTTT<br>GGCGCCGCCCAGGCCATCAAGGACAAGTACACGCCGCGCTGGAGGTGGCTGGCCCCCTGGTCCGAG<br>TCCATCCAGAAGGACCTCCGCCGCATCCGCAGGCAGTCCATCGACCTGCTGCGGCCGCTGTACAAGGA<br>GCGCCGCGACGCCATGGCCGACCGGGACGGCTTCCGGGACCCGGCCGACACCTTCAGGGACGTCGT<br>CTACTGGCTCATGAAGAGCGGCCAGCGGGACAGGTCGCTGTACGGCGTCACCGAGTCGCAGCTGTTCC<br>TGTCGCTGACGGCCATCCACACGACGTCGGGCACGCTCAACTCGTTCGTCTACGACTGGATCGCCCACC<br>CCGAGTACCACGACGACATCCTGCACGAGGTCACCGAGACCCTGGCCAAGGTGCGCGCGAACAACGG<br>CGAGTGGACGCTGCAGCACGTCGCCATGATGCGCAAGCTGGACAGCTTCATCAAGGAGAGTGCGCGCC<br>TCAACCCCATCGGCTTTGTGTCGACGCAGCGCTACACGCTCAAGCCCTACACCTTCAAGGACGGCTTCC<br>ACCTCCCCGCCGGCACGACCTTCATGTTCCACTCGGACGGCGCCCACTACGACGCCGACAACTACCC<br>GGACCCGGACAAGTTCGACGGCTACCGCTTCCTCCGCCTGCGCGAGACCGTCGACCCCAACCGCTTC<br>CACTACGCCTCCGTCTCGGACAGCTCGCTGGGCTTCGGCGCGGGTATCCACGCGTGCCCGGGCCGCT<br>TCCTCAGCGCCGTCGTCATGAAGTTCTTCCTCGTCCACTTCATGACGGCGTACGAGCTCAAGTACGAGCA<br>CGGCGGCACCGAGCGGCTGCCGAACCTGTACAACGACAACACCAGCAGACCCAACCCGACGGTGAA<br>CCTGCTGGTCAGGAAGAGGCAGTAG | TVGSEYKSVPAYDLVCGFVARV<br>ASVALVGAPLCRNPAWQRIVV<br>ETTFVAFGAAQAIKDKYTPRWR<br>WLAPWSESIQKDLRRIRRQSID<br>LLRPLYKERRDAMADRDGFRD<br>PADTFRDVVYWLMKSGQRDR<br>SLYGVTESQLFLSLTAIHTTSGTL<br>NSFVYDWIAHPEYHDDILHEVT<br>ETLAKVRANNGEWTLQHVAM<br>MRKLDSFIKESARLNPIGFVST<br>QRYTLKPYTFKDGFHLPAGTTF<br>MFHSDGAHYDADNYPDPDKF<br>DGYRFLRLRETVDPNRFHYASV<br>SDSSLGFGAGIHACPGRFLSAV<br>VMKFFLVHFMTAYELKYEHGGT<br>ERLPNLYNDNTSRPNPTVNLLV<br>RKRQ |
| MrCYP1 | ATGTTGTCGAGCATTCAGACAAAAATTGTCGAGCCGTTTCTGGTGCTGCGCCAGAGTGTTGCACCTCTGAAA<br>CTGTCCCGATGGCAGTTTACAAAGCTCATGGTCAGAAGTGCGCTTGACGGGTTGCCAGACGGAAGCGTAT<br>TTCTCCTACTTTACCTCTTCGCAGCACTGCTGATAATCATTACCGTGCGCCTTCGAAACAGCAAGAAGGGTG<br>TGCACGCGCCTCCTGGGCTTGCCGTGGTGAGGAGGAACCATGCGCATTACCTTGACATTATCAAGGAGG<br>GACGGGAGCTGGTATGTGCAGACTTGAATCTAGACTTGACCCATTGATATGTCATGATTACCGTCTACCCTTC<br>TGCCTGCCGAAAAGAACTTGGAGGCTCCGATTTTGACAGATGGCAGTACCCAGGTCAACCCTTCCTCGCT<br>GTCAACAACCGGCACAGTTTTGTCATATTCCCTCCCCAATGCTTCGACGAGATCAAGCGTCTTCCGGAGCA<br>CACCGCGTCTGCCAAGGGTTTCTTTCACGCTACAAACTATGGCCACTGGAGTCACATTGGTACAGAGACA<br>CCGCAACTCATCAAATCCGTCATTGCCGATCTGACTCGTTCGCTTCCTGCTCGAGTCCTCACGCGCCAAC<br>AAGACTGCCAGATGGCCTTTGACGACGTAATCGGGCGCTCACGCCAGTGGAAAGAATTCCCTCTGATGAT<br>GACTACATTCGAAATTGTCACCCGGATAAATGCGTGTTCCTTCGTTGGAAGAGAACTGGGCACGAACCGAA<br>GCTGGGTGCGGGCCGTCATGATGTCGCCCATCTTCATTCACGTTGCTGTCACGCTCCTCAACGCCTGCC<br>CCCTTATTCTGCGGCCGCTCATGGCGCCGATTTGCTTCTTCCCTACCATCAAGAACCGGTGGGATATGGC<br>GCGCCTTCTGACCCCAGTTCTAAAGACAGACATGAAGGATTACTACGAAGCCGAGGACAAGAAGGAGATC<br>CTGCGGCCCCGGGCAGAGGGAAAGATACCCTTCACCGGATTCCTCCTCTCACGCTACCAGGCTGCTGA<br>AGCAACCATTAAACAGCTGGTGGCTGACTACATCATCATCAGCTTCGACTCAACCCCGTCCACTGCATCGA<br>CGCTGTTCCACGTTCTCTGTGAGTTGGCATTGCACCCTAAAGCTGCCGACATTCTGCGCCAGGAACTAGAT | MLSSIQTKIVEPFLVLRQSVAPL<br>KLSRWQFTKLMVRSALDGLPD<br>GSVFLLLYLFAALLIIITVRLRNSK<br>KGVHAPPGLAVVRRNHAHYLD<br>IIKEGRELYPGQPFLAVNNRHS<br>FVIFPPQCFDEIKRLPEHTASAK<br>GFFHATNYGHWSHIGTETPQLI<br>KSVIADLTRSLPARVLTRQQDC<br>QMAFDDVIGRSRQWKEFPLM<br>MTTFEIVTRINACSFVGRELGTN<br>RSWVRAVMMSPIFIHVAVTLLN<br>ACPLILRPLMAPICFFPTIKNRW<br>DMARLLTPVLKTDMKDYYEAE<br>DKKEILRPRAEGKIPFTGFLLSR<br>YQAAEATIKQLVADYIIISFDSTP<br>STASTLFHVLCELALHPKAADIL<br>RQELDEVLVDGNLPGTHLQEL |

| | | |
|---|---|---|
| | GAGGTTCTTGTCGACGGAAACCTCCCCGGGACCCATCTCCAGGAGCTGAGGAAAATGGACAGTTTCCTTC<br>GTGAGTCCTTCCGACTACATCCAATTGTCATGTGTAAGTCGTCTTCACCCGTTGGGCCAGGCCTGCCAAAG<br>GCGCCCATTCCCTTCCCCGACGGCCCTATGCTGACTTTGCCTTCCAGTCACCCTCCGACGCCACCTAGA<br>AAAGCCGGTGAAACTCTCTCTTGGGCCAACGCTTCCCGCTGGCTTGATCATCGCTGTTGACGGACAAGCT<br>ATCAACCGCAACCCGGATTTGTGGCCAAACCCAGATAGTTTCGACATGGATAGATCTTATAAGCTTCGACAG<br>AAGCCAGGGAATGAAAACCGCTTCCATTTCCTGACTACCGGCTCCGACTCGCCGGGATGGGGTGACGG<br>GACACAGGCTTGCCCGGGACGGTTTTTTGCGACGAATACACTCAAGATTGCGCTGGCGCATTTTCTCAAG<br>AATTATGACATTGAGATTAAGCCAGAATGTTTGCCACTCAAGCATACACCCCCTTTCGAATGGATCTTGGAAAC<br>CTGATGATGCGGCCATCGCACGCATTAGGTCCAGATCTTAG | RKMDSFLRESFRLHPIVMFTLR<br>RHLEKPVKLSLGPTLPAGLIIAV<br>DGQAINRNPDLWPNPDSFDM<br>DRSYKLRQKPGNENRFHFLTT<br>GSDSPGWGDGTQACPGRFFA<br>TNTLKIALAHFLKNYDIEIKPECL<br>PLKHTPLSNGSWKPDDAAIARI<br>RSRS |

**Table S5:** Cytochrome P450 monooxygenase sequences investigated in this study. Where BGCs have been experimentally validated, the sequence deposited in NCBI was used; for the cryptic BGCs under investigation the putative gene sequences (or mRNA sequences) listed in this table were used.

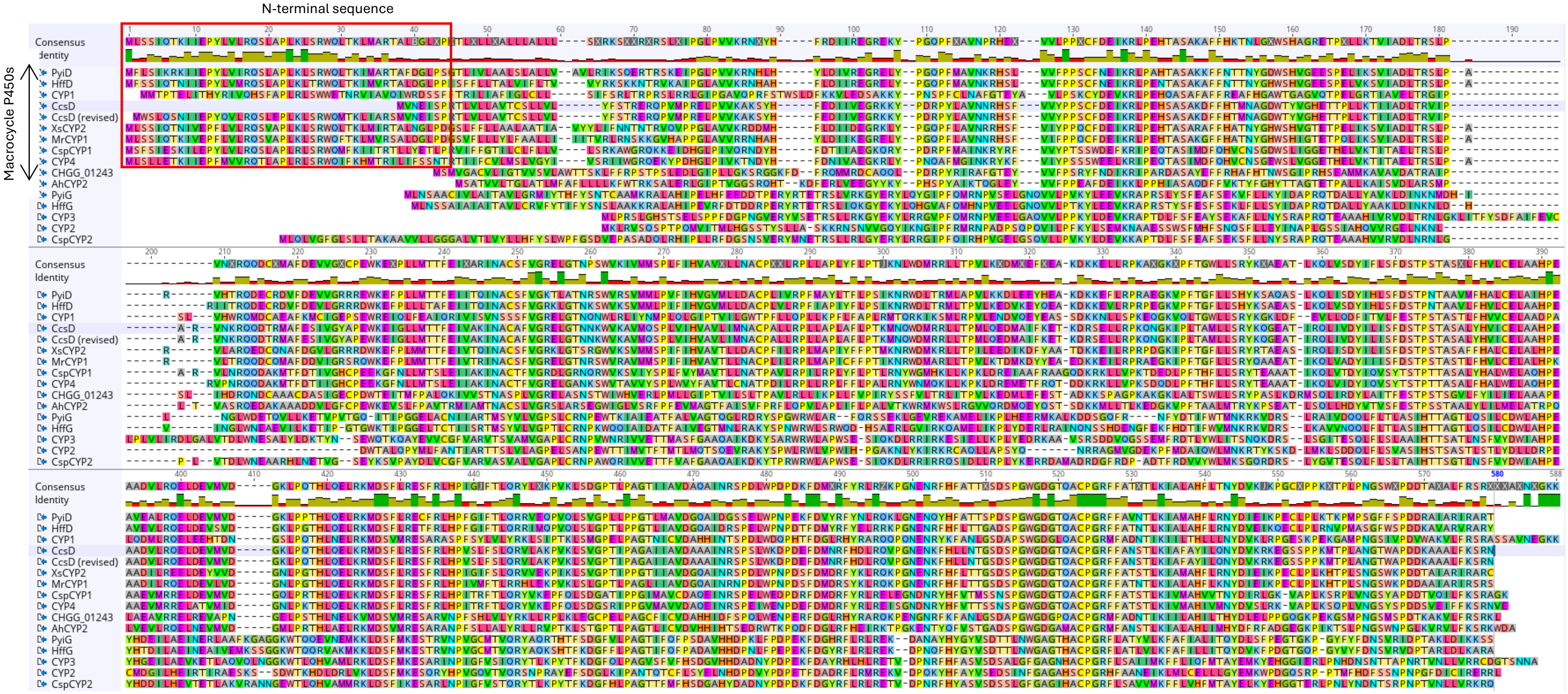


**Figure S5**: Multiple sequence alignment (MSA) of experimentally validated and / or bioinformatically linked P450 sequences used in this study. Macrocycle P450s are indicated with the black arrow and the N-terminal $\alpha$-helix sequence is highlighted with a red box.

#### 1.3 Strains and culture conditions

*Aspergillus heteromorphus* CBS 117.55 was previously purchased from the Agriculture Research Service (ARS / NNRL) culture collection and cultivated on GMM agar at 25 °C. The media panel used consisted of complete medium (CM; 2 g/L yeast extract, 2 g/L malt extract broth, 2 g/L mycological peptone, 2 g/L casamino acids), marine broth (MB; 37.4 g/L Millipore Marine Broth 2216), MEP medium (MEPA recipe without agar), GNB medium (20 g/L glucose, 30 g/L Oxoid Nutrient Broth No. 2), and potato dextrose broth (PDB; 24 g/L potato dextrose broth) grown as surface cultures at 25 °C for 5 – 10 days. Cells were filtered from the surface cultures and stored at –80 °C until ready for total RNA extraction and cDNA synthesis following the manufacturer's instructions. The supernatant was extracted as described below to prepare extracts for HRLCMS/MS analysis.

*Aspergillus clavatus* NRRL1 was purchased from the ARS / NRRL culture collection and cultivated on either PDA or MEPA (30 g/L malt extract, 3 g/L papaic digest of soybean meal, and 15 g/L of agar) for 5 days. To prepare protoplasts, we followed the previously published protocol established to knock-out *ccsB*. [16] Briefly, *A. clavatus* was grown on glucose minimal medium (GMM: 10 g/L glucose, 50 mL 20x salt solution, 1 mL trace elements, 15 g/L agar; 20x salt solution: 120 g/L $NaNO_3$, 10.4 g/L KCl, 10.4 g/L $MgSO_4.7H_2O$, 30.4 g/L $KH_2PO_4$; trace element solution: 2.2 g $ZnSO_4.7H_2O$, 1.1 g $H_3BO_3$, 0.5g $MnCl_2.4H_2O$, 0.5g $FeSO_4.7H_2O$, 0.16 g/L $CoCl_2.5H_2O$, 0.16 g/L $CuSO_4.5H_2O$, 0.11g $(NH_4)_6Mo_7O_{24}.4H_2O$, 5g $Na_4EDTA$ made up to 100 mL with $ddH_2O$) for 2 -3 days and resultant spores / mycelia were used to inoculate 50 mL of glucose minimal medium (GMM without agar) in a 250 mL Erlenmeyer flask, incubated at 28 °C with shaking (200rpm) for 16 hours. After transformation of protoplasts, 100 μL of the protoplast mixture was plated on individual SMMT agar plates (1.84 g/L ammonium tartrate, 0.52 g/L KCl, 1.52 g/L $KH_2PO_4$, 0.52 g/L $MgSO_4.7H_2O$, 10 g/L glucose, 218.6 g/L sorbitol, and 18 g/L agar, supplemented with 22 mg/L of $ZnSO_4$, 11 mg/L $H_3BO_3$, 5 mg/L $MnCl_2.4H_2O$, 5 mg/L $FeSO_4.7H_2O$, 1.6 mg/L $CoCl_2$.5H2O, 1.6 mg/mL $CuSO_4.7H_2O$, 1.1 mg/L $(NH_4)_6Mo_7O_{24}.4H_2O$, and 50 mg/L $Na_2EDTA$, pH 6.5) containing 100 μg/mL hygromycin B. 5 mL of soft agar SMMT was overlayed on top of the SMMT plates once set and incubated for 5 – 7 days at 25 °C. To confirm cytochalasin E production (Figure S15) *A. clavatus* NRRL1 was grown in MEP media as a surface culture for 5 days at 25 °C and the culture broth was extracted as described below.

For heterologous expression studies *M. grisea* NI980 Δ*pyiD* and *M. grisea* NI980 Δ*pyiG* were used as the wild-type strains in this study. [8,9] Strains were cultivated on TNK-(SU)-CP Agar (10 g/l glucose, 2 g/L yeast extract, 2 g/L $NaNO_3$, 5 g/L $MgSO_4.7H_2O$, 0.1 g/L $CaCl_2.2H_2O$, 4 mg/L $FeSO_4.7H_2O$, 2 g/L $KH_2PO_4$ and 15 g/L agar. For TNK-SU-CP agar, 200 g/L sucrose was added) or Complete Medium (CM) agar (10 g/L glucose, 2 g/L peptone, 1 g/L yeast extract, 1 g/L casamino acids, 0.1 % (v/v) trace elements (22 mg/L zinc sulphate heptahydrate, 11 mg/L boric acid, 5 mg/L manganese (II) chloride tetrahydrate, 5 mg /L iron (II) sulphate heptahydrate, 1.7 mg /L cobalt (II) chloride hexahydrate, 1.6 mg/L copper (II) sulphate pentahydrate, 1.5 mg/L sodium molybdate dehydrate, 50 mg/L ethylenediaminetraacetic acid), 0.1 % (v/v) vitamin supplement (0.001 g/L biotin, 0.001 g/L pyridoxine, 0.001 g/L thiamine, 0.001 g/L riboflavin, 0.001 g/L, 0.001 g/L nicotinic acid), 6 g/L $NaNO_3$, 0.5 g/L KCl, 0.5 g/L $MgSO_4$, 1.5 g/L $KH_2PO_4$, [pH adjusted to 6.5 with NaOH], 15 g/L agar) containing 200 mg/mL phosphinothricin (Basta / glufosinate) at 25 °C for 7 - 14 days. For cytochalasan production *M. grisea* host strains and transformants were cultivated in DPY liquid medium (2% w/v dextrin from potato starch, 1% w/v polypeptone, 0.5% w/v yeast extract, 0.5% w/v monopotassium phosphate, 0.05% w/v magnesium sulfate, 2.5% w/v agar) for 7 days at 25 °C, 120 rpm. Strains generated in this study are listed in Table S6.

| **Fungal Strain** | **Description** |
|---|---|
| *A. clavatus NRRL 1* | Cytochalasin E and K producer. [6] |
| *A. heteromorphus* CBS 117.55 | Potentially produces cytochalasans but none have yet been reported. |
| *M. grisea* NI980 | Pyrichalasin H producer. [8,9] |
| *M. grisea ΔPyiD* | *M. grisea ΔpyiD* mutant, no longer produces pyrichalasin H. Instead produces precursors lacking C-18 hydroxyl group. [8,9] |
| *M. grisea ΔPyiG* | *M. grisea ΔpyiD* mutant, no longer produces pyrichalasin H. Instead produces precursors lacking C-7 hydroxyl group. [8,9] |
| *M. grisea ΔPyiD* + CcsD | *M. grisea ΔpyiD* mutant, expressing CcsD. |
| *M. grisea ΔPyiD* + CHGG_01243 | *M. grisea ΔpyiD* mutant, expressing CHGG_01243. |
| *M. grisea ΔPyiD* + XsCYP2 | *M. grisea ΔpyiD* mutant, expressing XsCYP2. |
| *M. grisea ΔPyiD* + CsCYP1 | *M. grisea ΔpyiD* mutant, expressing CsCYP1. |
| *M. grisea ΔPyiD* + AhCYP2 | *M. grisea ΔpyiD* mutant, expressing AhCYP2. |
| *M. grisea ΔPyiD* + MrCYP2 | *M. grisea ΔpyiD* mutant, expressing MrCYP2. |
| *M. grisea ΔPyiG* + CspCYP2 | *M. grisea ΔpyiG* mutant, expressing CspCYP2. |

**Table S6**: Strains generated in this study

#### 1.4 Transformation methods

Transformation of *S. cerevisiae* YPH 499 for yeast recombination and *E. coli* Top10 or CopyCutter EPI400 was performed using standard methods described previously. Preparation and transformation of *M. grisea* protoplasts followed previous methods. [8,9]

To transform *A. clavatus* NRRL1, we followed the previously published protocol established to knock-out *ccsB* with minor modifications. [16] Briefly, *A. clavatus* was grown on GMM for 2 -3 days and resultant spores / mycelia were used to inoculate 50 mL of glucose minimal medium (GMM without agar) in a 250 mL Erlenmeyer flask, incubated at 28 °C with shaking (200rpm) for 16 hours. The culture was transferred to a 50 mL centrifuge tube and centrifuged at 4 °C for 10 minutes (3750 rpm). The resultant cell pellet was transferred to a new 250 mL Erlenmeyer flask containing 30 mg Driselase and 20 mg Yatalase dissolved in 10 mL of Osmotic medium (1.2 M $MgSO_4$, 10mM sodium phosphate buffer) and incubated at 25 °C for 6 - 8 hours (80 rpm). The solution was transferred to a fresh 50 mL centrifuge tube and overlayed with 10 mL of trapping buffer (0.6 M sorbitol, 0.1 M Tris-HCl, pH 7.0) and centrifuged at 4 °C for 15 mins (5000 rpm). The buffers were gently decanted from the opposite side of the centrifuge tube from the resultant protoplast pellet. The pellet was gently dissolved in 1 – 5 mL of STC buffer (1.2 M sorbitol, 10 mM Tris-HCl, 10 mM $CaCl_2$, pH 7.5) and centrifuged at 4 °C for 5 mins (6000 rpm). The supernatant was carefully discarded and the protoplast pellet diluted with ~ 1mL of STC buffer. 100 μL of PEG solution (400 mg/mL polyethylene glycol 3350, 50 mM $CaCl_2$, and 10 mM Tris-HCl, pH 8) was added to the protoplast solution, followed by 2.5 - 5 μg of the PCR fragments for targeted gene inactivation, and the mixture was incubated for 20 mins at 4 °C. 1 mL of ice-cold PEG solution was added to the reaction mixture, and then incubated at room temperature for 5 minutes. 100 μL of the protoplast mixture was plated on individual SMMT agar plates containing 100 μg/mL hygromycin B. 5 mL of soft agar SMMT was overlayed on top of the SMMT plates once set and incubated for 5 – 7 days at 25 °C.

#### 1. 5 Attempted knock-out of *ccsX*

A bipartite knock-out strategy was attempted using synthetic DNA fragments (Table S7) that targeted ~ 1000 bp of *ccsX* including upstream / downstream regions due to the small size of the thioredoxin-like gene (Figure S6A). The 5' targeting fragment included the *trpC* promoter and part of the hygromycin resistance gene (*hph*). The 3' targeting fragment included 500 bp overlap of the hygromycin resistance gene (*hph*). Transformation of the fragments proceeded as described above. As colonies appeared on original SMMT plates supplemented with hygromycin (Figure S6B), they were transferred to small SMMT agar plates and once visible growth appeared were screened using Platinum Direct PCR Universal master Mix kit (Invitrogen) to determine if the TRX gene was successfully disrupted (Figure S6C).

| **Fragment name** | **Synthetic DNA sequence** |
|---|---|
| 5'-TRX-TrpC-Hyg | GAATCTCCTAACATTGCGAACCTGCTTGTCAACCAAGATCTCTAGTGATTTATCCCCAACAATT<br>ATTCTCTGGTTGGTTATTCCAAAAATACACTATCTAGTAGTCAAATGGCGAAAGAGCGTCCATAT<br>GCTCCGATGATATTGGTTCGGCTCACCGAGTCTGAGACTCAACATCGGAGCCGCGGGGTG<br>GAAGATCCGACAAACCCCTGCAGCTTGTTGCGGTTGGGCATCATCAGACATATTCTGTTCCC<br>CCCATCATCATCTTTCATCCGTCGTCTTGGTCTGTTTATCTATCTCCAAGTAGGCCCTTCTTTTT<br>TTCGAATCCGGTAGTCGCGACCGCATCCATTTCACCATGTCCGTTATCGAGATCGAAGTGGT<br>CTATGATTTTGTTTGCGCGGTACACAGCGATGACCTTCCACCGTTCGAGGAGAGAAAGGCTG<br>ACAAGGGTCTAGTGGTGCTACATTGGCAAACGCAAACTCGACCGAGCCATCGCGCTGTATC<br>AGAAAACCTACCCTGGCGGTCGATCTGATGTTTTCTCGATCAAATGGAGCCCGTATTTCTTGA<br>ACTACAACCCTCACCCTCACAGCGTGCCCAAGAGCGACCTGGTGGATGAACGGCTGAAG<br>GACATGACTCCGGAGCAGCGAACCGCCTTGTTCAACCGGATGAACCAAATTGGACGTGCG<br>GTGGGCATCTACTTCAAGGCTGGGGGGATGATCGGGAGCACCCGTGATGCCCATCGCCTG<br>GTCCACCTGAGCCGGACCAAGCCCGCAGACGTTCAAAATTGCCTGGTCGAGAACATTATAT<br>TGAAGGAGCACTTTTTGGGCTTGGCTGGAGCTAGTGGAGGTCAACAATGAATGCCTATTTTG<br>GTTTAGTCGTCCAGGCGGATCACAAAATTTGTGTCGTTTGACAAGATGGTTCATTTAGGCAACT<br>GGTCAGATCAGCCCCACTTGTAAGCAGTAGCGGCGGCGCTCGAAGTGTGACTCTTATTAGC<br>AGACAGGAACGAGGACATTATTATCATCTGCTGCTTGGTGCACGATAACTTGGTGCGTTTGTC<br>AAGCAAGGTAAGTGAACGACCCGGTCATACCTTCTTAAGTTCGCCCTTCCTCCCTTTATTTCA<br>GATTCAATCTGACTTACCTATTCTACCCAAGCGCTTCGATTAGGAAGTAACCATGCCTGAACT<br>CACCGCGACGTCTGTCGAGAAGTTTCTGATCGAAAAGTTCGACAGCGTCTCCGACCTGATG<br>CAGCTCTCGGAGGGCGAAGAATCTCGTGCTTTCAGCTTCGATGTAGGAGGGCGTGGATATG<br>TCCTGCGGGTAAATAGCTGCGCCGATGGTTTCTACAAAGATCGTTATGTTTATCGGCACTTTG<br>CATCGGCCGCGCTCCCGATTCCGGAAGTGCTTGACATTGGGGAATTCAGCGAGAGCCTGA<br>CCTATTGCATCTCCCGCCGTGCACAGGGTGTCACGTTGCAAGACCTGCCTGAAACCGAACT |

| | |
|---|---|
| | GCCCGCTGTTCTGCAGCCGGTCGCGGAGGCCATGGATGCGATCGCTGCGGCCGATCTTAGCCAGACGAGCGGGTTCGGCCCATTCGGACCGCAAGGAATCGGTCAATACACTACATGGCGTGATTTCATATGCGCGATTGCTGATCCCCATGTGTATCACTGGCAAACTGTGATGGACGACACCGTCAGTGCGTCCGTCGCGCAGGCTCTCGATGAGCTGATGCTTTGGGCCGAGGACTGCCCCGAAGTCCGGCACCTCGTGCACGCGGATTTCGGCTCCAACAATGTCCTGACGGACAATGGCCGCATAACAGCGGTCATTGACTGGAGCGAGGCGATGTTCGGGGATTCCCAATACGAGGTCGCCAACATCTTCTTCTGGAGGCCGT |
| Hyg-3'-TRX | GACATTGGGGAATTCAGCGAGAGCCTGACCTATTGCATCTCCCGCCGTGCACAGGGTGTCACGTTGCAAGACCTGCCTGAAACCGAACTGCCCGCTGTTCTGCAGCCGGTCGCGGAGGCCATGGATGCGATCGCTGCGGCCGATCTTAGCCAGACGAGCGGGTTCGGCCCATTCGGACCGCAAGGAATCGGTCAATACACTACATGGCGTGATTTCATATGCGCGATTGCTGATCCCCATGTGTATCACTGGCAAACTGTGATGGACGACACCGTCAGTGCGTCCGTCGCGCAGGCTCTCGATGAGCTGATGCTTTGGGCCGAGGACTGCCCCGAAGTCCGGCACCTCGTGCACGCGGATTTCGGCTCCAACAATGTCCTGACGGACAATGGCCGCATAACAGCGGTCATTGACTGGAGCGAGGCGATGTTCGGGGATTCCCAATACGAGGTCGCCAACATCTTCTTCTGGAGGCCGTGGTTGGCTTGTATGGAGCAGCAGACGCGCTACTTCGAGCGGAGGCATCCGGAGCTTGCAGGATCGCCGCGGCTCCGGGCGTATATGCTCCGCATTGGTCTTGACCAACTCTATCAGAGCTTGGTTGACGGCAATTTCGATGATGCAGCTTGGGCGCAGGGTCGATGCGACGCAATCGTCCGATCCGGAGCCGGGACTGTCGGGCGTACACAAATCGCCCGCAGAAGCGCGGCCGTCTGGACCGATGGCTGTGTAGAAGTACTCGCCGATAGTGGAAACCGACGCCCCAGCACTCGTCCGAGGGCAAAGGAATAGCTCCGGGCCTACCACGAAATGGAGATGGATATCTCATCAAAGGACGTACTTCTGGAGCTCGCCTTGAGCGCGGGACTCGACAAGGCCGAAGTGACTGAATGGCTCGAATCCGACTTGGCCGGGGATATCGTCGACGAAGCGTCTCAAAGAAACAGGCAGCCGGGCAACACCGGAGTTCCACGGTATATCATTCAGGGAGTGCATTGCGTGGATGGTGCAGAGGATCCGTCGGAGTTCATCGAGGTGTTTGCAAAGGTCAAGGAAGGCGAAAATCAGGCTTAGGGGGTCACCCCTGACGAGTGCTTGGAACTATTACCGTAACCAGCTATTGGATCCCGGGAGGTTCTTTCGAATCATAGCAAATCTTGGAATCCACGATCAGGTCAGATGATCCCAGTTTCGTATCCGCTTCCGATGGTACTTTGGACTTGTGGTGTGGGTTGTTGAGATGGTGTTGGGATCGACGGATTCGAGCTCTGTACTCCGTACGGTCGCACCGCCCCCAACAACAAAGAACAGGGTGTTCTGGGGGGGGTCAAGATAAGTAATGATGGTTAGGATTAGTTAGGATGCGGATTGACTAATAATTCCTGTCTGCGATTTCTCATGCGAAGTCGGAGTTGATCCCTTGGGGGGGGGGGAAGATTGTGTAGAATGCAGATGCTTTTTCCGGCGCCCAAACTTCCTAAAAAGAAAAAACAAAGAAACCCGAAAAAAACCACCGAGTTGGAGAGTTGGAGCTGTCATCTTAACCCCTTAAAATAATATGAAAGTATTTTTGGATGCATAGCTCGGTTGTG |

**Table S7:** Synthetic DNA fragment sequences used to disrupt the TRX-like gene *ccsX* in *A. clavatus* NRRL1.

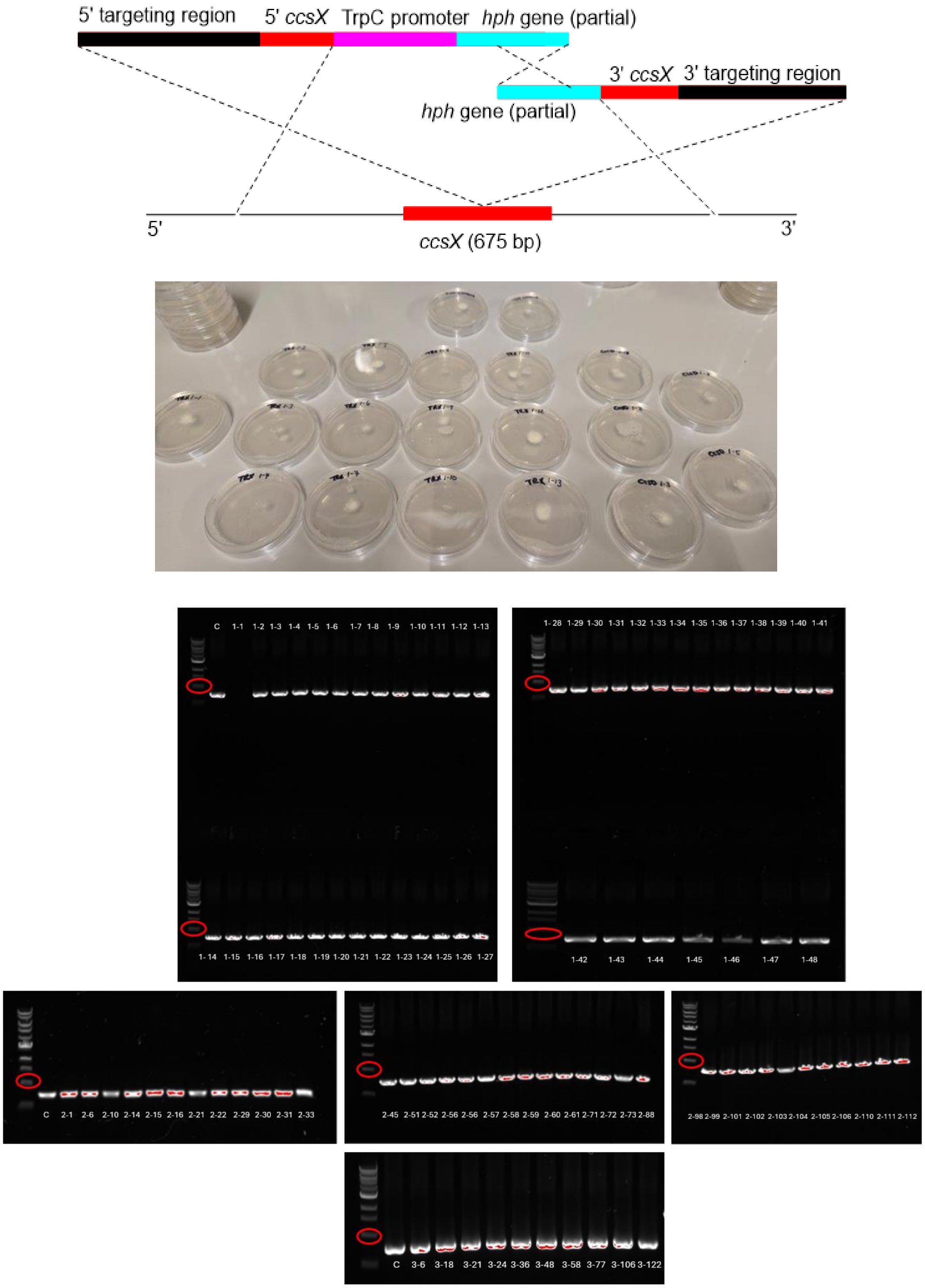


**Figure S6**: A) Bipartite KO strategy for *ccsX* using synthetic DNA fragments; B) Secondary transformant plates containing hygromycin to check stability of transformants; C) PCR screening of transformants from secondary plates showing *ccsX* (675 bp) has not been disrupted. The 1 kb ladder band is circled in red.

#### 1.5 Cloning and expression plasmids used in this study

Construction of fungal expression vectors was performed using cloning vectors compatible with Invitrogen's Gateway system and LR Clonase II to transfer genes from pTwist ENTR / pDONR plasmids to pTYGSbar (Figure S7). Alternatively, plasmids were constructed *via* homologous recombination using *Saccharomyces cerevisiae* to insert gene fragments into pTYGSbar.

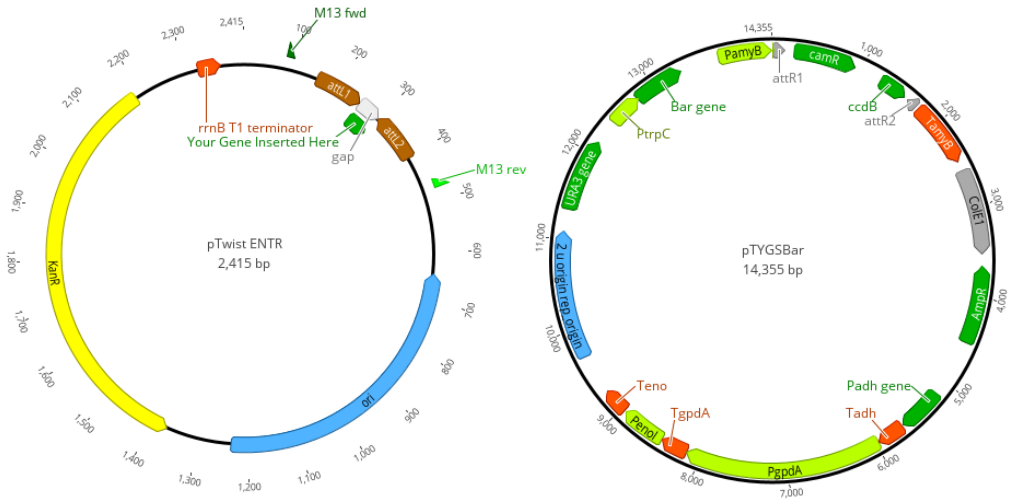


**Figure S7**: Plasmid backbone maps of pTwist ENTR and pTYGSbar. P450s are located between the attL1 and attL2 sites for LR recombination with pTYGSbar, resulting in the P450 of interest being expressed under the control of the amyB promoter.

| Plasmids used in this study | Description |
|---|---|
| pTYGSbar | Fungal/yeast/*E.coli* shuttle vector for expression of P450 genes in *M. grisea* mutant strains. [17] |
| pTwist ENTR + CHGG_01243 | Commercial cloning plasmid containing synthetic gene *CHGG_01243* |
| pTwist ENTR+ CcsD | Commercial cloning plasmid containing synthetic gene *ccsD* |
| pTwist ENTR+ CcsDrev | Commercial cloning plasmid containing synthetic gene *ccsD* with revised sequence |
| pDONR221 + XsCYP2 | Commercial cloning plasmid containing synthetic gene *XsCYP2* based on cDNA sequence |
| pTwist ENTR + AhCYP2 | Commercial cloning plasmid containing synthetic gene *AhCYP2* |
| pTwist ENTR + CsCYP1 | Commercial cloning plasmid containing synthetic gene *CsCYP1* |
| pTwist ENTR + CsCYP2 | Commercial cloning plasmid containing synthetic gene *CsCYP2* |
| pE-YA + MrCYP1 | Initial cloning vector prepared via yeast recombination containing a synthetic gene *MrCYP1* |
| pTYGSbar + CHGG_01243 | *M. grisea* expression vector; *CHGG_01243* cloned under the *amyB* promoter |
| pTYGSbar + CcsD/CcsDrev | *M. grisea* expression vector; *ccsD/ccsDrev* under the *amyB* promoter |

| | |
|---|---|
| pTYGS bar XsCYP2 | *M. grisea* expression vector; XsCYP2 cloned under the *amyB* promoter |
| pTYGSbar + AhCYP2 | *M. grisea* expression vector; AhCYP2 cloned under the *amyB* promoter |
| pTYGSbar + CsCYP1 | *M. grisea* expression vector; CsCYP1 cloned under the *amyB* promoter |
| pTYGSbar + CsCYP2 | *M. grisea* expression vector; CsCYP2 cloned under the *amyB* promoter |
| pTYGSbar + MrCYP1 | *M. grisea* expression vector; MrCYP1 cloned under the *amyB* promoter |

**Table S8**: Plasmids used in this study.

**1.6 DNA isolation, polymerase chain reaction (PCR) and Reverse Transcription PCR (RT-PCR)**

Genomic DNA and / or RNA was prepared from stored frozen cells using commercial kits, following the manufacturer's protocol unless otherwise stated. Plasmid DNA was prepared from fresh *E. coli* / yeast cells using commercial kits, following the manufacturer's protocol. DNA and RNA integrity was confirmed using agarose gel electrophoresis stained with SYBRSafe and visualized using a GelDoc Go imaging system (BioRad) (Figure S8). cDNA was prepared from wild-type strains or randomly selected transformants where LCMS analysis did not appear to show chemical changes.

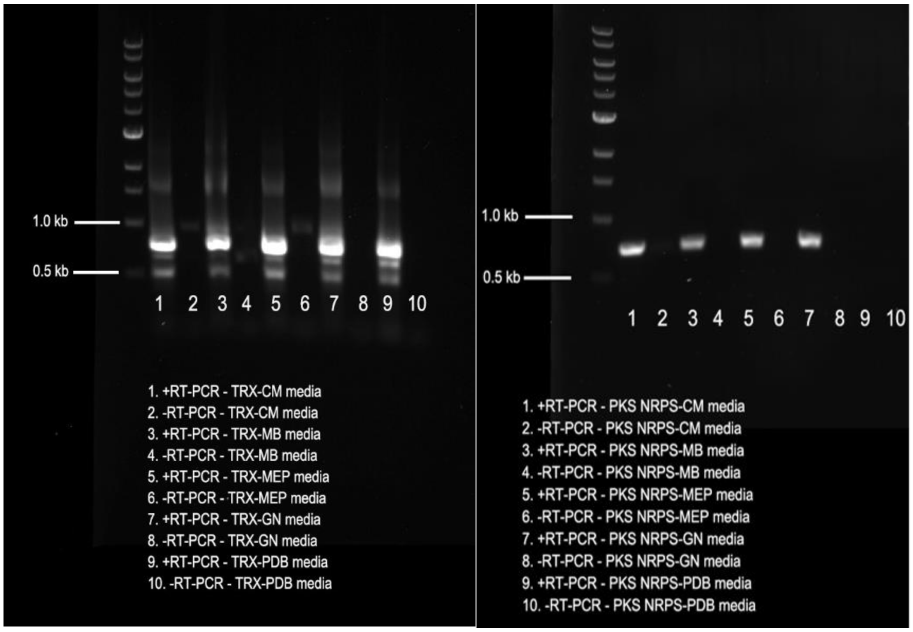


**Figure S8**: Gel electrophoresis image of the RT-PCR products for the *A. heteromorphus* TRX-like gene and a fragment of the PKS-NRPS gene present in the *ahe* putative cytochalasan BGC. Cells were obtained from the media panel investigation and the gels are labelled accordingly.

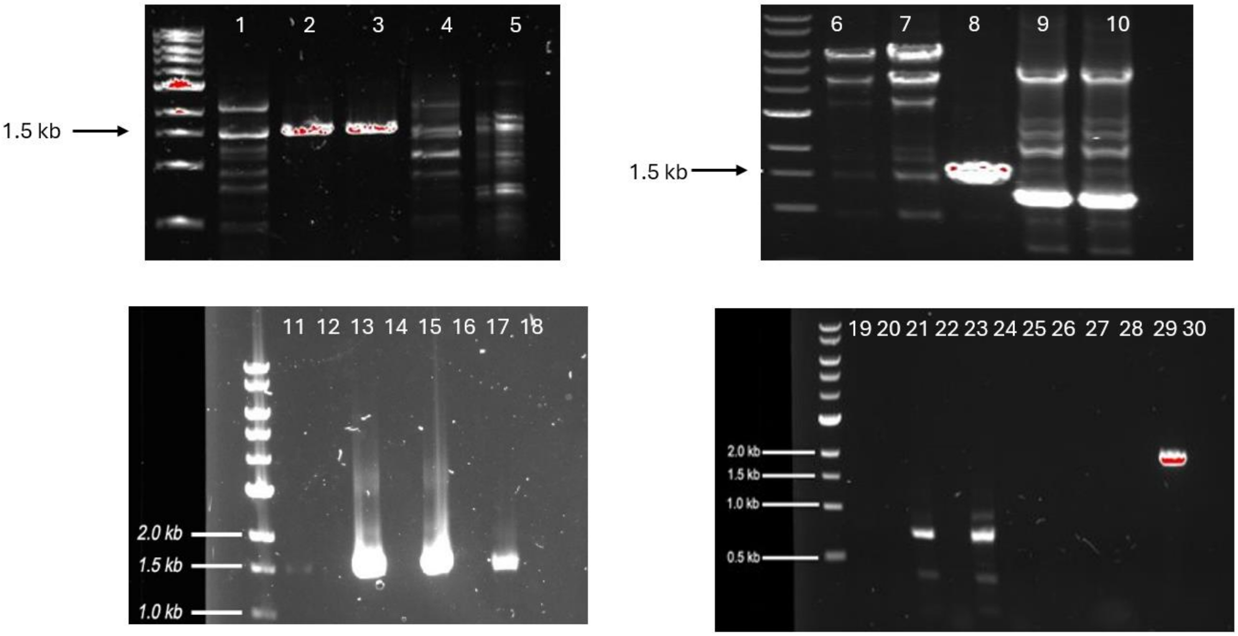


**Figure S9**: Gel electrophoresis images of PCR / RT-PCR products relevant to this study. Lane 1 = CcsD-Tf11 PCR product (50 °C annealing temperature); Lane 2 = CcsD-Tf12 PCR product (55 °C annealing temperature); CcsD-Tf13 PCR product (55 °C annealing temperature); Lane 4 = MrCYP1-Tf1 PCR product (50 °C annealing temperature); Lane 5 = XsCYP2-Tf3 PCR product (50° C annealing temperature); Lane 6 = XsCYP2-Tf1 PCR product (50 °C annealing temperature); Lane 7 = XsCYP2-Tf2 PCR product (50 °C annealing temperature); Lane 8 = XsCYP2-Tf3 PCR product (50 °C annealing temperature); Lane 9 = CsCYP1-Tf1 PCR product (50 °C annealing temperature); Lane 10 = CsCYP1-Tf1 PCR product (50 °C annealing temperature); Lane 11 = CcsD-Tf11 RT-PCR product; Lane 12 = CcsD-Tf11 RT-PCR product (no RT enzyme added); Lane 13 = CcsD-Tf12 RT-PCR product; Lane 14 = CcsD-Tf12 RT-PCR product (no RT enzyme added); Lane 15 = CcsD-Tf13 RT-PCR product; Lane 16 = CcsD-Tf13 RT-PCR product (no RT enzyme added); Lane 17 = *A. clavatus* NRRL1 total RNA; Lane 18 = *A. clavatus* NRRL1 total RNA (no RT enzyme added); Lane 19 = MrCYP1-Tf1 RT-PCR product; Lane 20 = MrCYP1-Tf1 RT-PCR product (no RT enzyme added); Lane 21 = CsCYP1-Tf1 RT-PCR product; Lane 22 = CsCYP1-Tf1 RT-PCR product (no RT enzyme added); Lane 23 = CsCYP1-Tf2 RT-PCR product; Lane 24 = CsCYP1-Tf2 RT-PCR product (no RT enzyme added); Lane 25 = XsCYP2-Tf1 RT-PCR product; Lane 26 = XsCYP2-Tf1 RT-PCR product (no RT enzyme added); Lane 27 = XsCYP2-Tf2 RT-PCR product; Lane 28 = XsCYP2-Tf2 RT-PCR product (no RT enzyme added); Lane 29 = XsCYP2-Tf3 RT-PCR product; Lane 30 = XsCYP2-Tf3 RT-PCR product (no RT enzyme added).

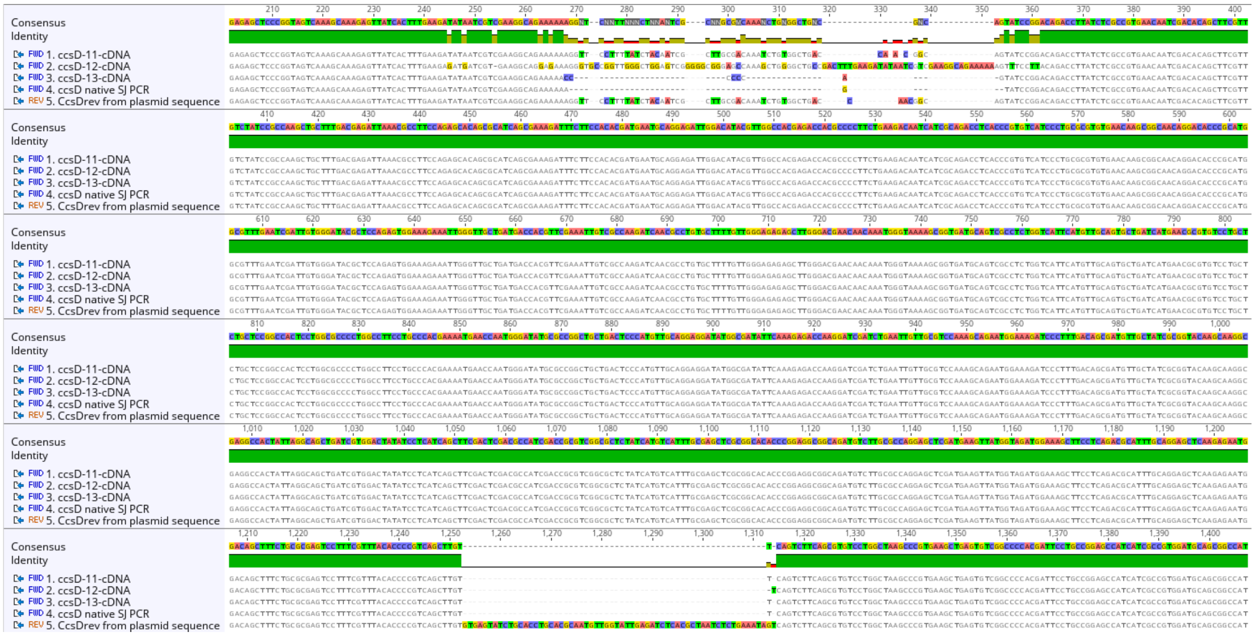

**Figure S10**: cDNA sequence results of CcsD transformants 11, 12, and 13 showing differences with splicing intron 1 (top), but precise splicing of intron 2 (bottom).

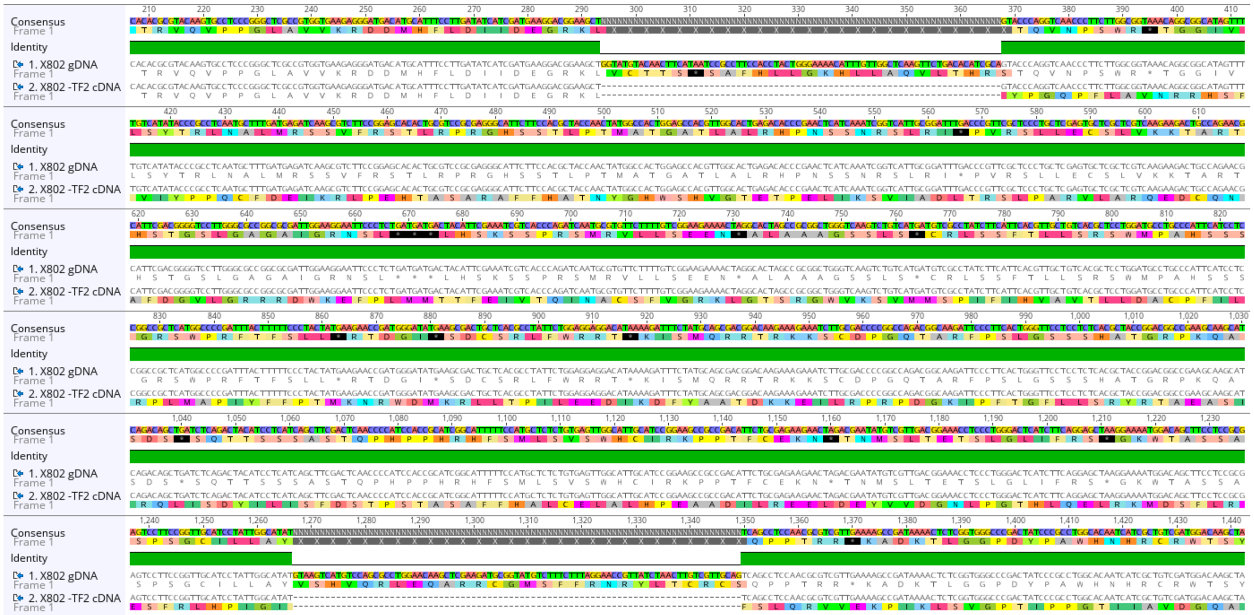

**Figure S11**: cDNA sequence results of XsCYP2-Tf2 showing expected splicing of introns 1 and 2.

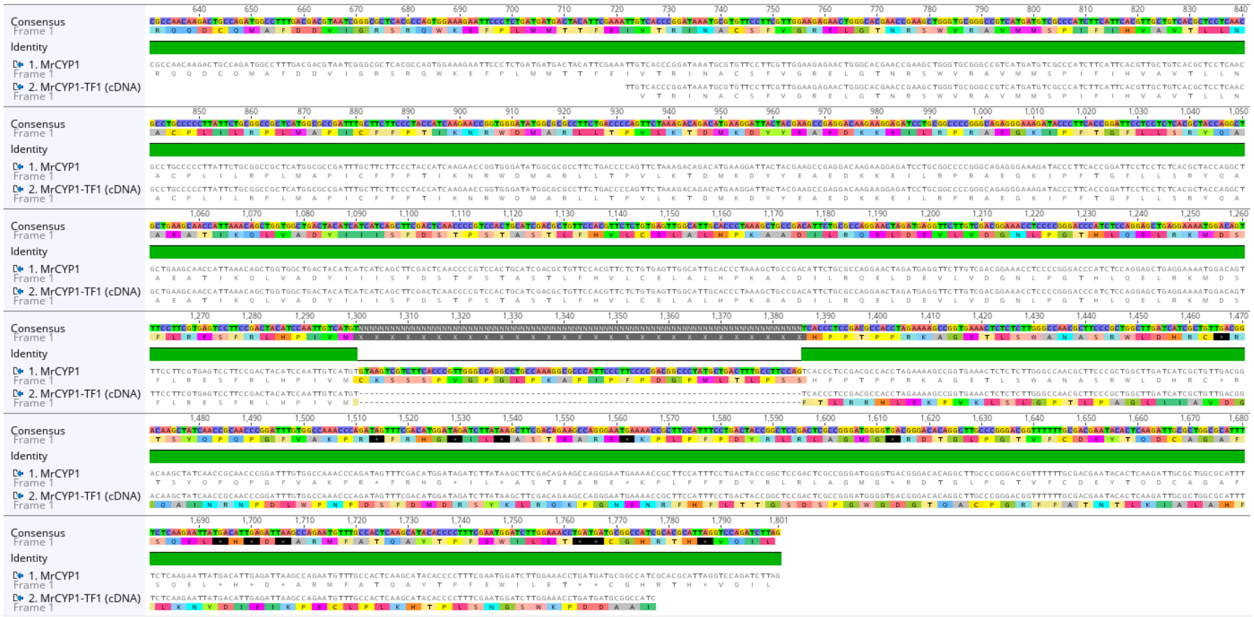


**Figure S12**: partial cDNA sequencing results of MrCYP1-Tf1 confirming intron 2 is correctly spliced.

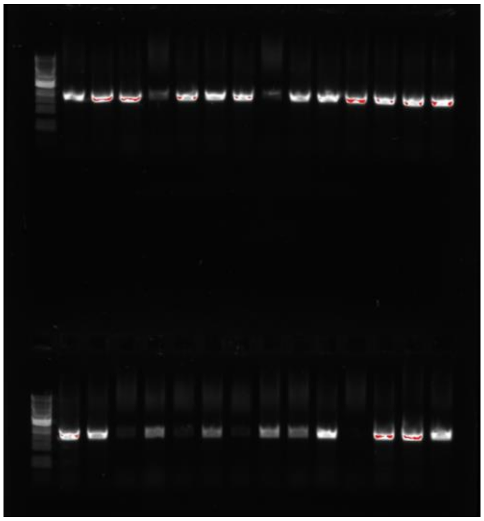

**Figure S13:** PCR screening of *ccsDrev* transformants confirming integration into the genome of *M. grisea* Δ*pyiD* in 23 out of 28 transformants generated.

### 1.7 Extraction method

Fungal strains were initially cultivated on solid agar and up to 1mL of mycelial spore suspensions were used to inoculate either 100 mL of appropriate media in 500 mL Erlenmeyer flasks or 20 mL of media in a Petri dish. After 7 days, the cultures were filtered using a sterilized Buchner gravity filter funnel and extracted twice with an equal volume of EtOAc. The combined EtOAc layers were dried over anhydrous $Mg_2SO_4$ and evaporated using a reduced pressure rotary evaporator. The dry crude extract was resuspended in methanol prior to LCMS analysis. Cells were flash frozen in liquid $N_2$ and stored at -80 °C for DNA and RNA analysis.

#### 1.8 Biotransformation of exogenous cytochalasans

5 mg of 7-deshydroxypyrichalasin H was purified from a 7-day old culture of *M. grisea ΔpyiG* using preparative LCMS. The structure was confirmed using NMR (Figures S69-70). 7-deshydroxypyrichalasin H was dissolved in 300 μL of DMSO and pulse fed to 50 mL cultures of *M. grisea ΔpyiD* on days 4, 5, and 6. The culture was extracted on the 7th day and the extract was analyzed using LCMS to confirm pyrichalasin H production. DMSO was used as a control in a second culture of *M. grisea ΔpyiD* grown under the same conditions.

#### 1.9 Chromatography methods

Analytical HPLC-PDA-MS screening of fungal extracts was performed on a Shimadzu instrument (LC2030C 3D Plus Prominence) coupled to a Shimadzu LCMS-2020 mass spectrometer. Analyses were performed using a Phenomenex Kinetex $RP_{18}$ column (100 mm × 4.6 mm i.d., 2.6 μm) combined with the Security Guard $RP_{18}$ protective guard column (4.6 mm i.d.) and eluting with $H_2O$ + 0.1% formic acid and MeCN + 0.1% formic acid using a gradient from 90:10 to 10:90 of $H_2O$/MeCN over 15 min, maintaining at 10:90 $H_2O$/MeCN for 3 min, from 10:90 to 90:10 over 1 min, and maintaining at 90:10 for 1 min, using a flow rate of 0.8 mL/min. The PDA detector scanned between λ = 190 and 700 nm. The MS was optimized using the following conditions: interface voltage 4.5 kV; interface temperature 350 °C; DL temperature 250 °C; heat block 200 °C; ESI mode, acquisition range 100 to 1000 Da; nebulizing gas 1.5 L $min^{-1}$; drying gas flow 15 L $min^{-1}$.

Fractionation of the samples for purification was performed on a Shimadzu LC-20AP preparative liquid chromatograph (SCL-40 System Controller and LH-40 Liquid Handler) coupled to a Shimadzu SPD-M40 Photo Diode Array Detector (PDA) system using a RP-18 column (Phenomenex, Kinetex 250 × 30 mm i.d., 5 μm, flow rate of 18.0 mL $min^{-1}$). Further purification of compounds was performed on a Shimadzu LC-20AD liquid chromatography (CBM-20A Communication Bus Module, CTO-20A column oven, DGU-20A Degassing Unit and SIL-20A AutoSampler) coupled to a Shimadzu SPD-20A UV–vis Detector system using a RP-18 column (Shimadzu, Premier 250 × 10 mm i.d., 5 μm, flow rate of 3.0 mL $min^{-1}$).

High-resolution mass spectra were recorded on an ABSciex TripleTOF 6600+ mass spectrometer using the same column, solvents and method as the analytical LCMS work. The parameters such as declustering and entrance potentials remained constant for MS and MS/MS were set up at 150 V and 10 V, respectively. Collision energy for MS and $MS^2$ scan surveys was 10 V and 45 V, respectively, with a collision energy spread of 12 V for $MS^2$ scan survey. Precursor ion was impacted with three different collision energies (33, 45, 57 V), and the resulting $MS^2$ spectra were combined into one final $MS^2$ spectrum. The mass spectra were acquired using Turbo Spray Ionization set to 5.5 kV in positive ion mode with an accumulation time of 100 ms. The mass ranges for MS and $MS^2$ scan surveys were 400–800 amu and 30–800 amu, respectively. The curtain gas (nitrogen), nebulizing and heating gas were fixed at 25 psi, 20 psi and 15 psi, respectively. The temperature of the source was 25 °C. MS spectra were acquired and processed using Analyst TF 1.8.1 software.

### 1.10 Structural Elucidation methods

Raw positive and negative HPLC–QTOF MS/MS data (.wiff files, SCIEX 6600+ system) were processed using MZmine (v4.8.30) and MS-DIAL (v5.2.2).[18,19] Raw files were first converted to mzML format using MSConvert (v.3.0.2) and then imported into MZmine.[20] The mzwizard module was used to establish a standardized LC–MS/MS workflow optimized for the HPLC–QTOF system. The ion mode was selected as positive or negative according to the type of mass spectrometry data files. Chromatographic settings included smoothing, an RT range of 0 – 20 mins, up to 15 peaks per chromatogram, at least four consecutive scans, feature FWHM of ~ 0.10 min, intra-sample RT tolerance of 0.05 min, and inter-sample RT tolerance of 0.20 min. Noise thresholds were set to $5.0 \times 10^2$ (MS1) and $1.0 \times 10^2$ (MS2), with a minimum feature height of $1.0 \times 10^3$. Mass tolerances were 0.0050 Da (20 ppm) for scan-to-scan, 0.0015 Da (3 ppm) intra-sample, and 0.0040 Da (8 ppm) inter-sample alignment. All aligned features were retained (minimum aligned sample threshold = 1). Spectral library files from the MZmine database were used for annotation. For DIA analysis, a minimum correlation coefficient of 0.80 and at least five correlated points were applied. Finally, the automated-mzwizard workflow generated an aligned feature list, which was used to explore ms/ms spectra and analyze the presence and structural hits related to cytochalasin molecules in the extract.

1D and 2D NMR experiments were recorded on a Varian INOVA 400 instrument ($^1$H: 400 MHz; $^{13}$C: 1oo MHz). The chemical shifts (δ) were expressed in ppm and recorded with reference to TMS. Spectral data was analyzed using Mestrenova.

### 1.11 3D Molecular Models

ChemDraw Professional (version 23.1.2.7) was used to prepare 3D models of various cytochalasans

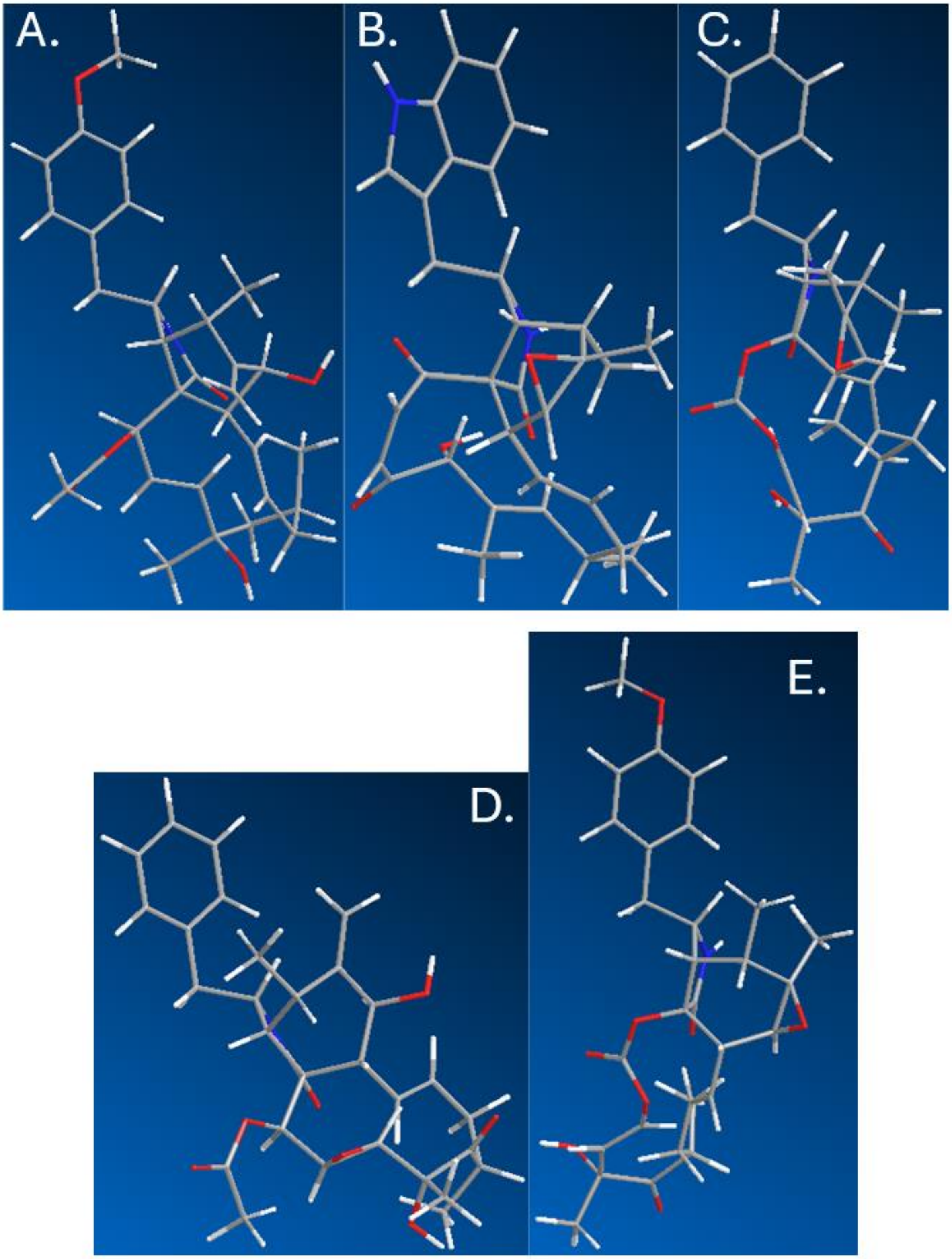


**Figure S14**: Structural models of cytochalasans relevant to this study A) pyrichalasin H; B) chaetoglobosin A; C) cytochalasin E; D) 19,20-epoxycytochalasin C; E) phenochalasin B

### 2. LCMS Chromatograms

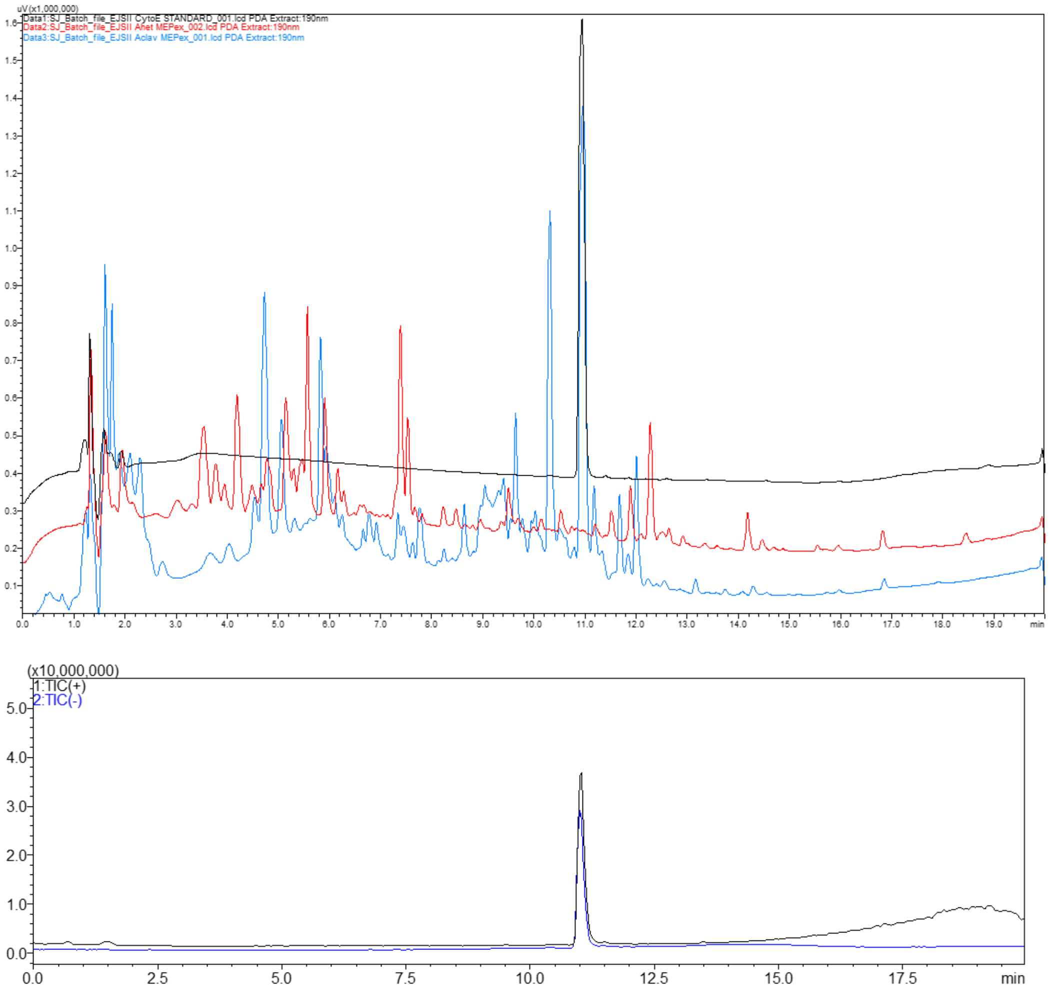


**Figure S15**: EtOAc extracts of *A. clavatus* NRRL1 (blue chromatogram) and *A. heteromorphus* CBS 117.55 (red chromatogram) after being grown in MEP medium for 5 days. A chemical standard of cytochalasin E (black chromatogram) confirms production of cytochalasin E by *A. clavatus* but not *A. heteromorphus*.

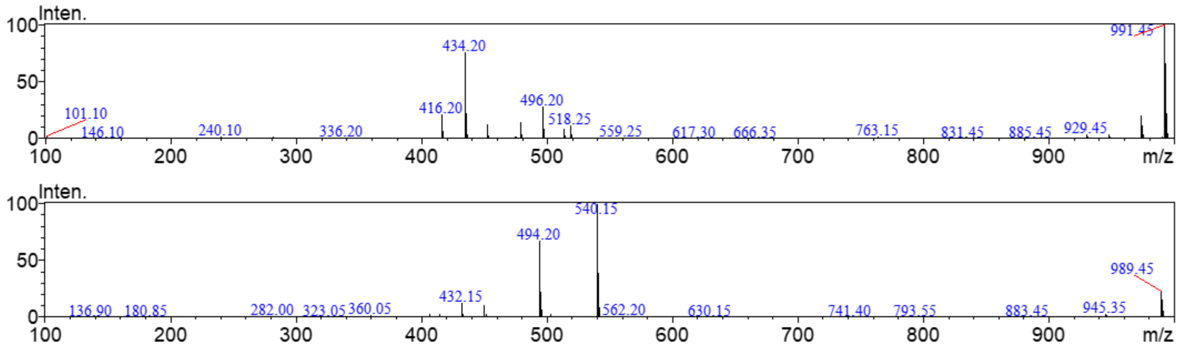


**Figure S16**: LCMS trace of a commercial cytochalasin E standard (*m/z* = 495). The top chromatogram overlays mass data obtained in positive and negative ion mode. The middle spectrum shows the mass spectrum of cytochalasin E (Rt = 11 mins) in positive ion mode and the bottom spectrum shows the mass spectrum obtained in negative ion mode.

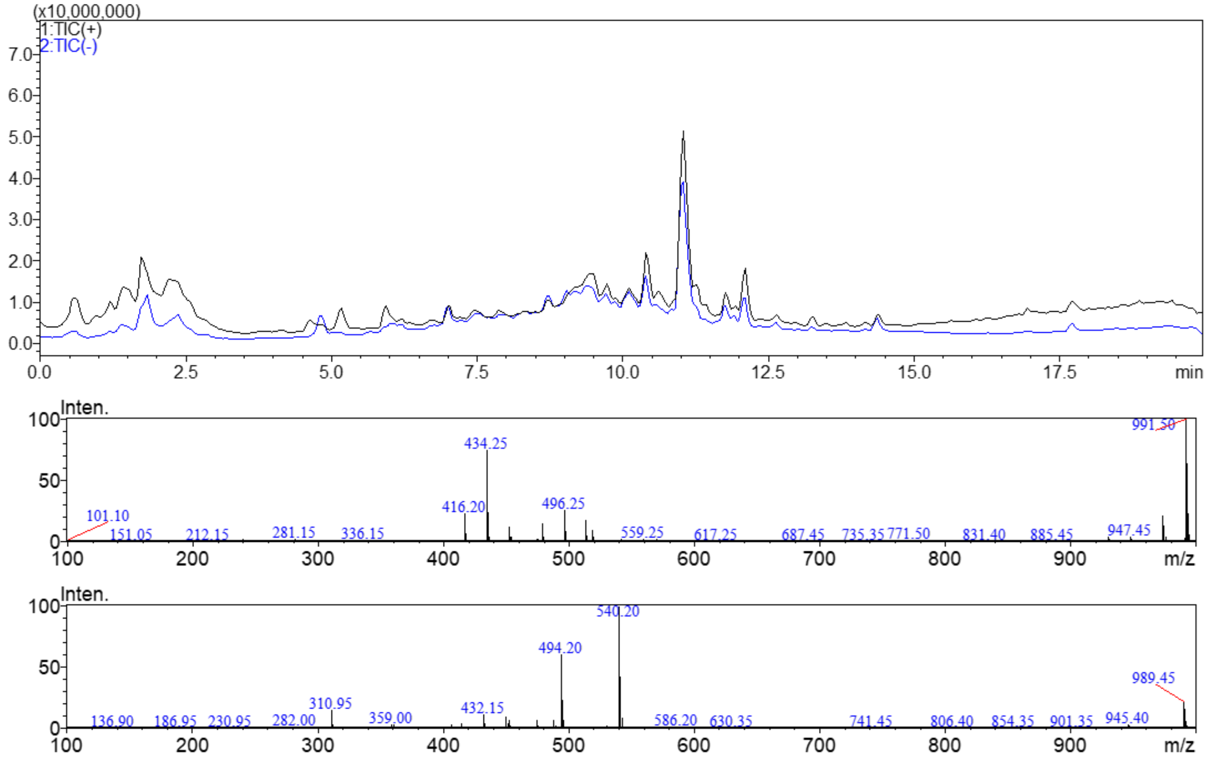


**Figure S17**: LCMS trace of *A. clavatus* EtOAc extract. The top chromatogram overlays mass data obtained in positive and negative ion mode. The middle spectrum shows the mass spectrum of cytochalasin E (Rt = 11 mins) in positive ion mode and the bottom spectrum shows the mass spectrum obtained in negative ion mode.

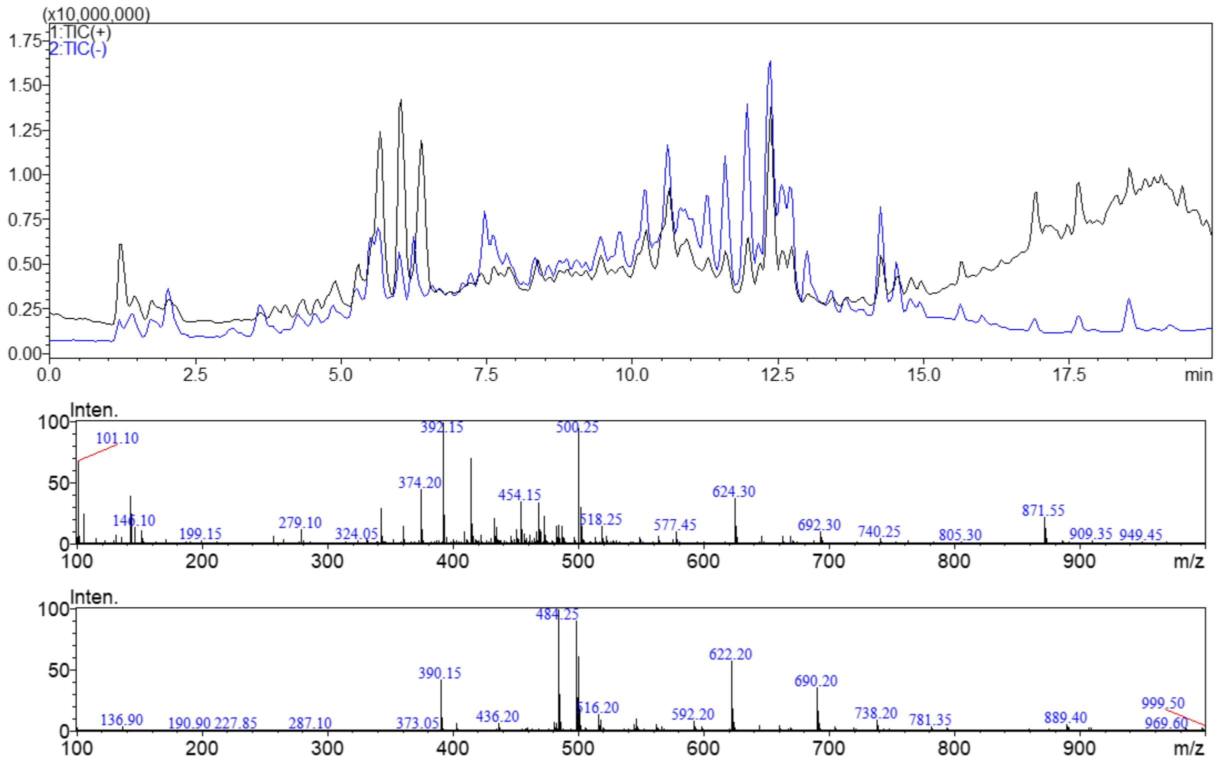


**Figure S18**: LCMS trace of *A. heteromorphus* EtOAc extract. The top chromatogram overlays mass data obtained in positive and negative ion mode. The middle spectrum shows the mass spectrum at Rt = 11 mins in positive ion mode and the bottom spectrum shows the mass spectrum obtained in negative ion mode. No cytochalasin E is detected.

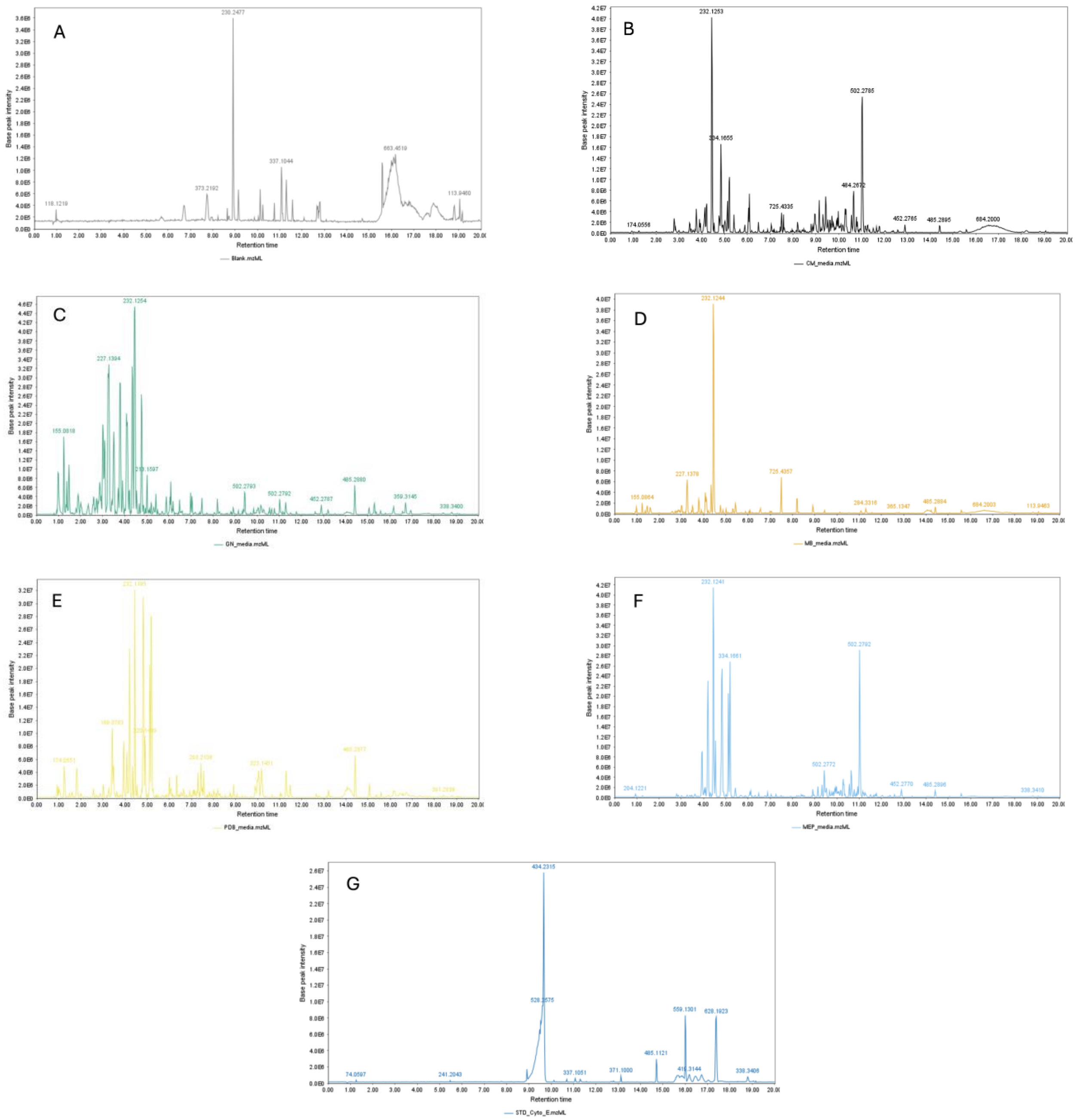


**Figure S19.** Total ion chromatograms (TICs) recorded in positive electrospray ionization (ESI+) mode. (A) Blank control; (B–F) extracts from cultures grown in CM, GN, MB, PDB, and MEP media, respectively; (G) Cytochalasin E standard.

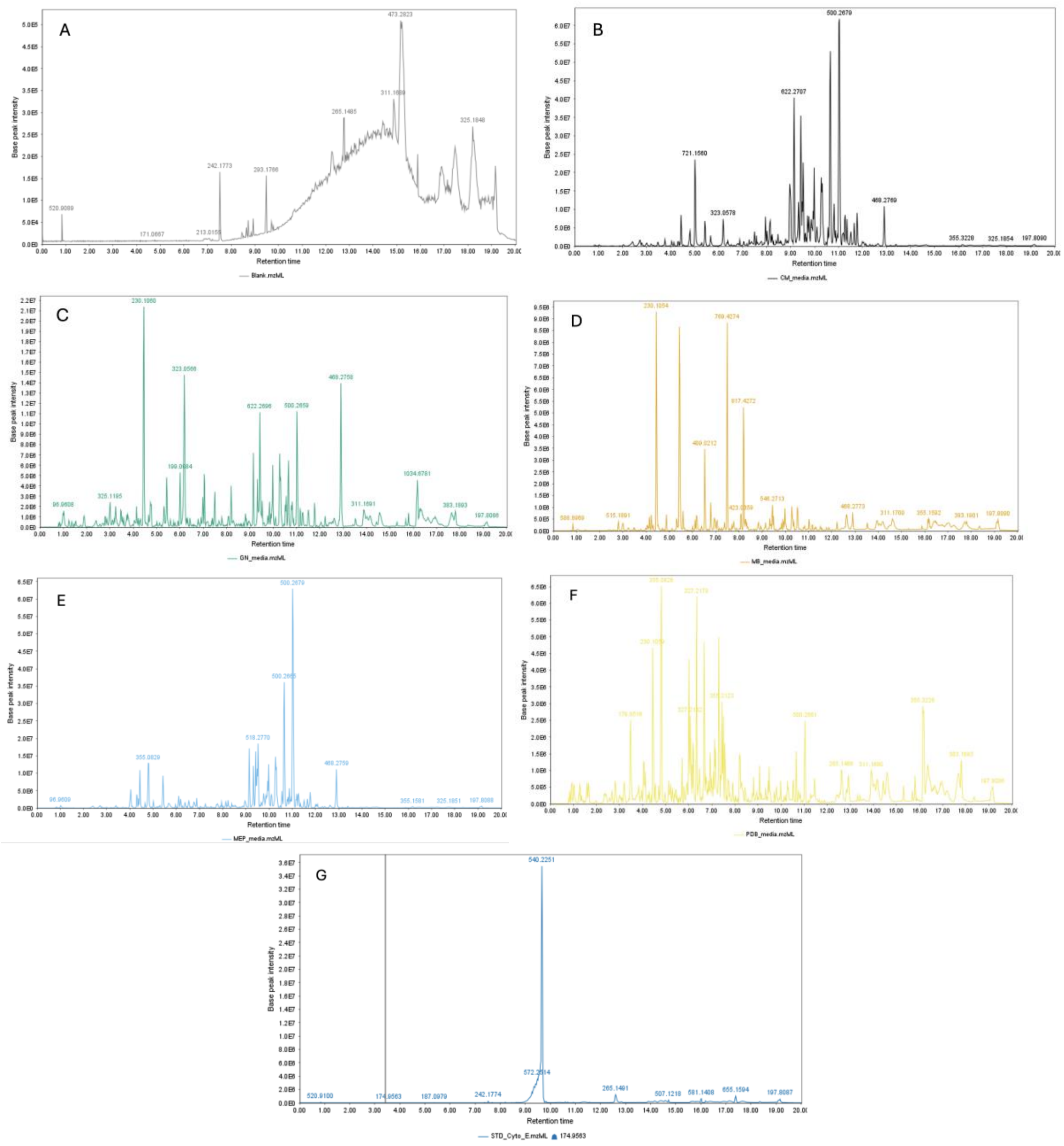

**Figure S20.** Total ion chromatograms (TICs) acquired in negative ion mode. (A) Blank; (B–F) extracts obtained from cultures grown in CM, GN, MB, MEP and PDB respectively; (G) Cytochalasin E standard.

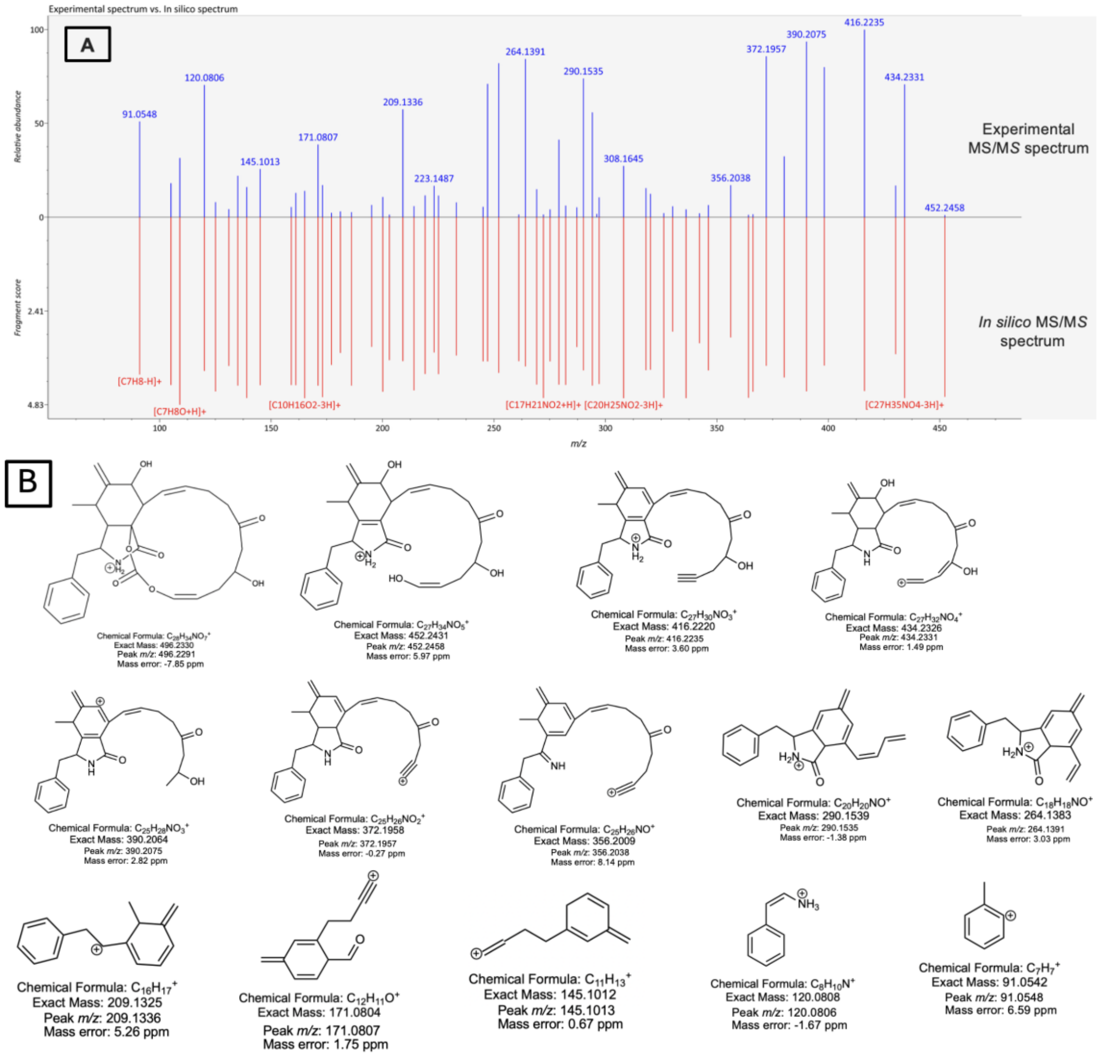


**Figure S21.** (A) Experimental MS/MS fragmentation spectrum of the ion at *m/z* 496.2291 acquired in positive ion mode, compared with the *in-silico* MS/MS spectrum generated by MS-FINDER. (B) MS/MS analysis of the ion at *m/z* 496.2291 reveals characteristic fragment ions consistent with a putative cytochalasin analogue.

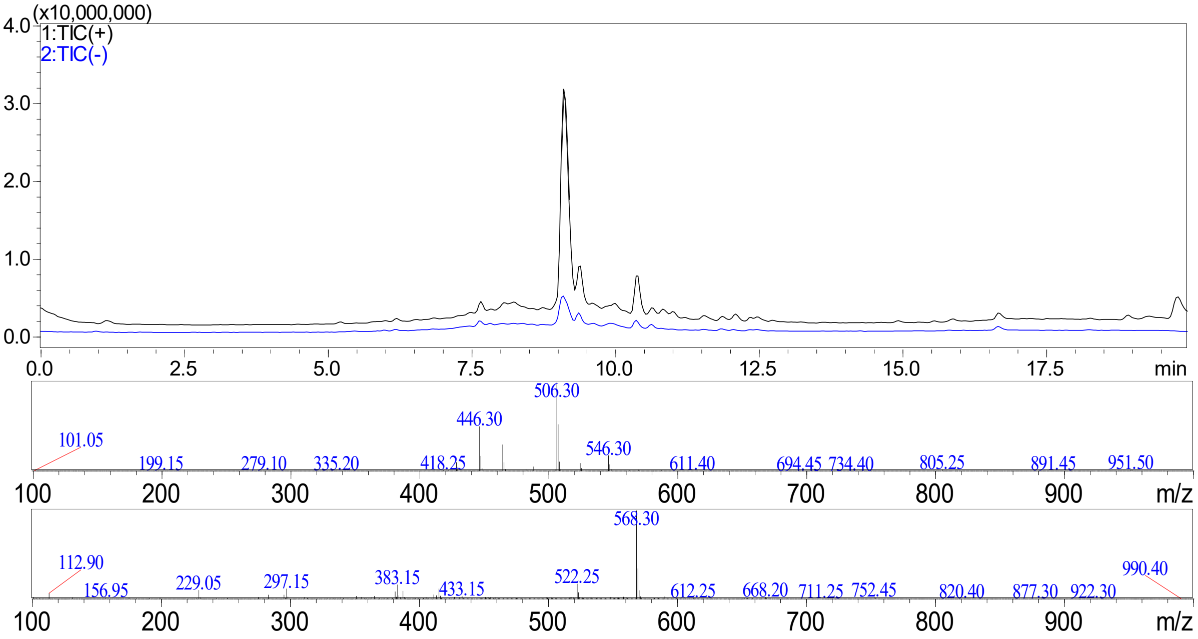


**Figure S22**: LCMS trace of pyrichalasin H purified from *M. grisea* NI980 for a standard. The top chromatogram overlays data obtained in positive and negative ion mode. The middle spectrum shows the mass spectrum of pyrichalasin H (Rt = 9.1) in positive ion mode and the bottom spectrum shows the mass spectrum of pyrichalasin H obtained in negative ion mode.

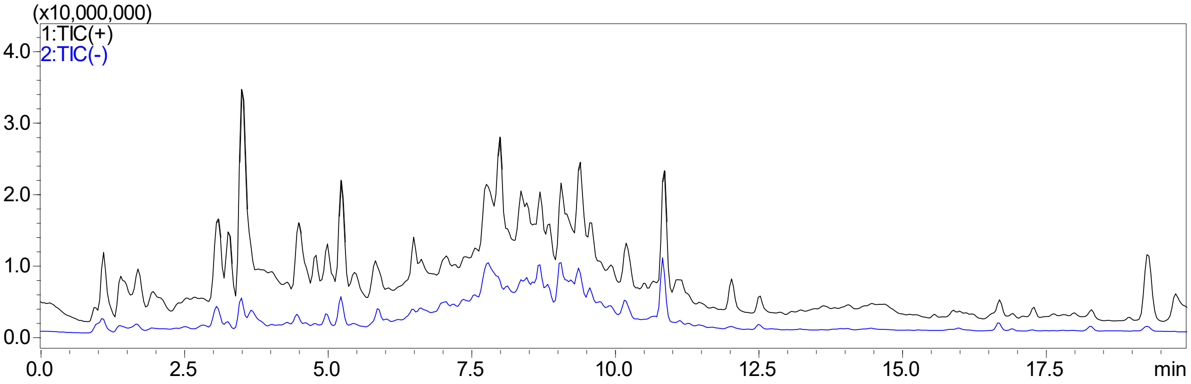


**Figure S23**: LCMS trace of *M. grisea* Δ*pyiD* host strain, overlaying data obtained in positive and negative ion mode. No pyrichalasin H is produced by this strain instead, unnamed cytochalasans *m/z* 509 (Rt = 7.8 mins), 451 (Rt = 8.7 mins), 523 (Rt = 9.4) and 493 (Rt = 10.9 mins) are produced.

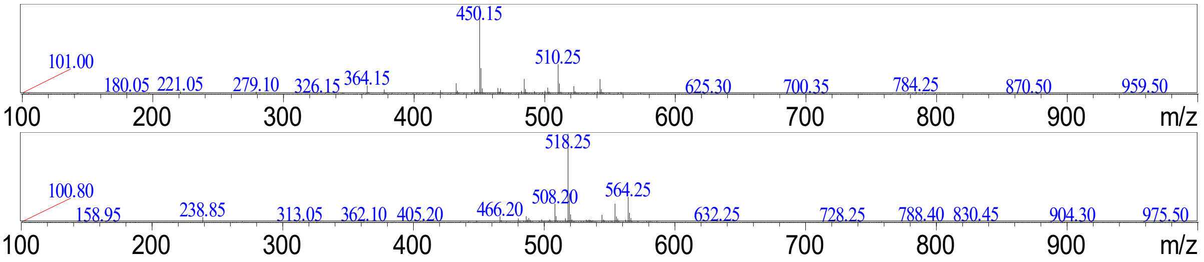


**Figure S24**: Mass spectrum of cytochalasan *m/z* 509 (Rt = 7.8 mins) in positive ion mode (top) and negative ion mode (bottom).

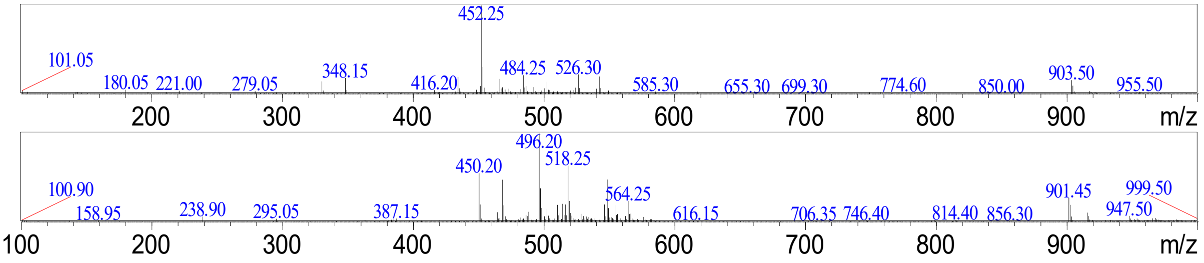


**Figure S25**: Mass spectrum of cytochalasan *m/z* 451 (Rt = 8.7 mins) in positive ion mode (top) and negative ion mode (bottom).

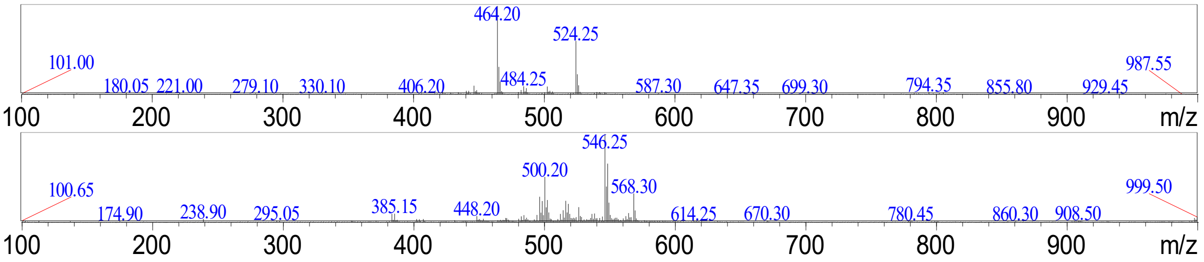


**Figure S26**: Mass spectrum of cytochalasan *m/z* 523 (Rt = 9.4 mins) in positive ion mode (top) and negative ion mode (bottom).

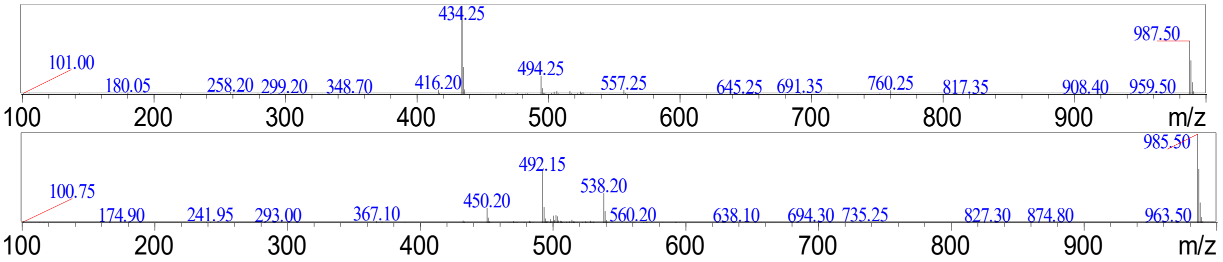


**Figure S27**: Mass spectrum of cytochalasan *m/z* 493 (Rt = 10.9 mins) in positive ion mode (top) and negative ion mode (bottom).

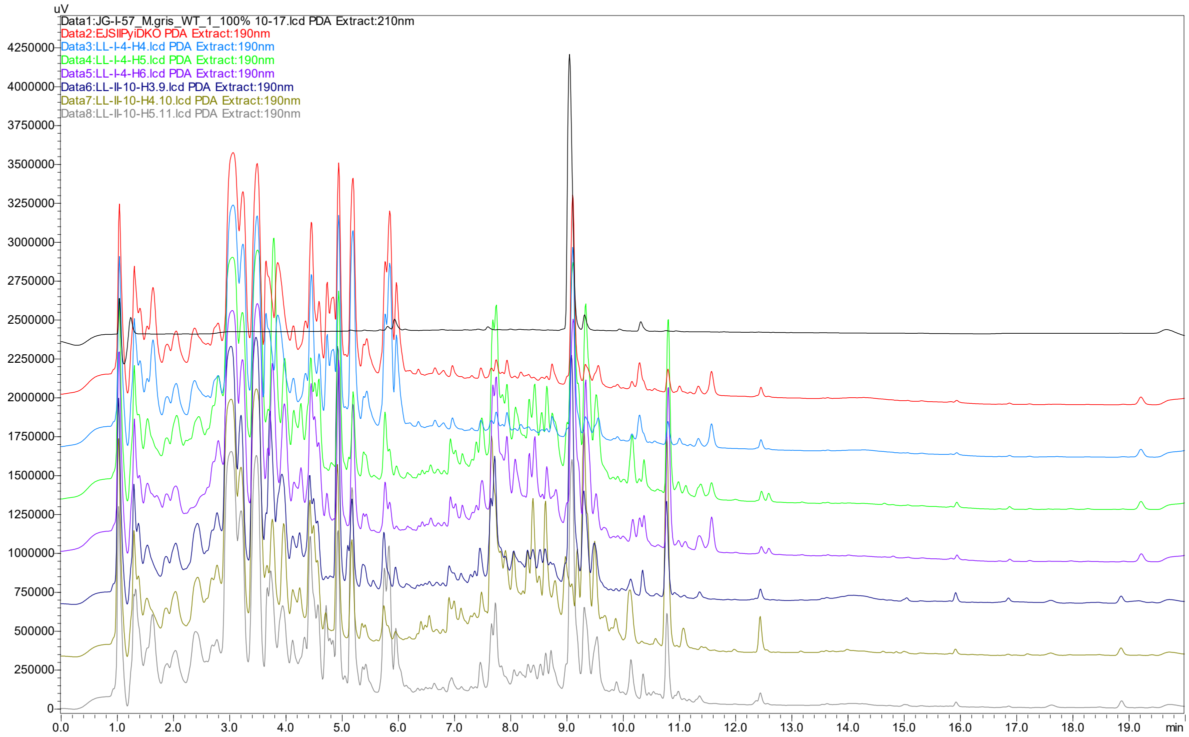


**Figure S28**: LCMS traces of *M. grisea* Δ*pyiD* + CHGG_01243 transformants, showing production of pyrichalasin H based on comparison to a standard (Rt = 9.1 mins). Compared to the control strain (EJSIIPyiDKO) no new compounds are detected.

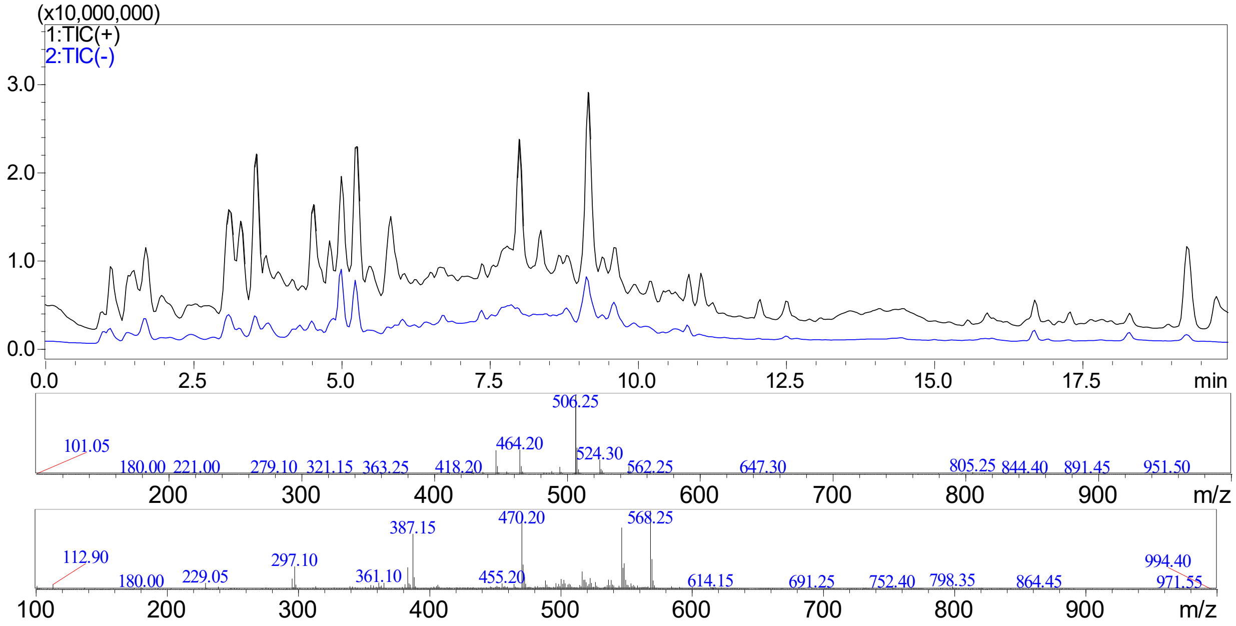


**Figure S29**: LCMS traces of *M. grisea* Δ*pyiD* + CHGG_01243 transformant LL-I-4-H4, showing production of pyrichalasin H at 9.1 mins (top chromatogram), and the corresponding mass spectra in positive ion mode (middle) and negative ion mode (bottom).

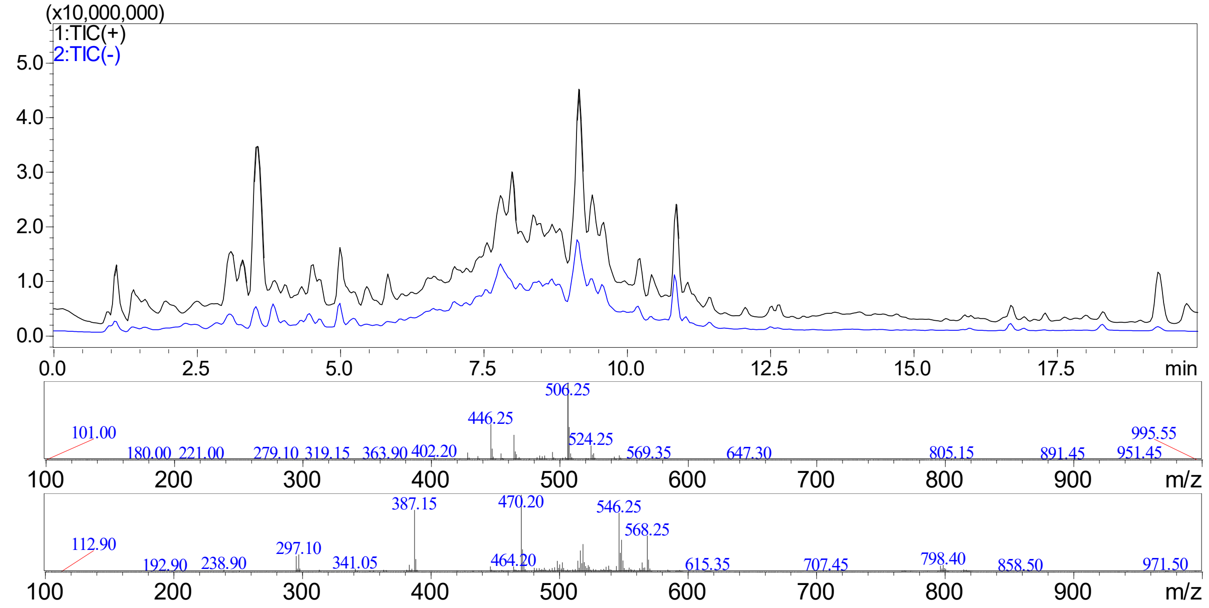


**Figure S30**: LCMS traces of *M. grisea* Δ*pyiD* + CHGG_01243 transformant LL-I-4-H5, showing production of pyrichalasin H at 9.1 mins (top trace), and the corresponding mass spectra in positive ion mode (middle) and negative ion mode (bottom).

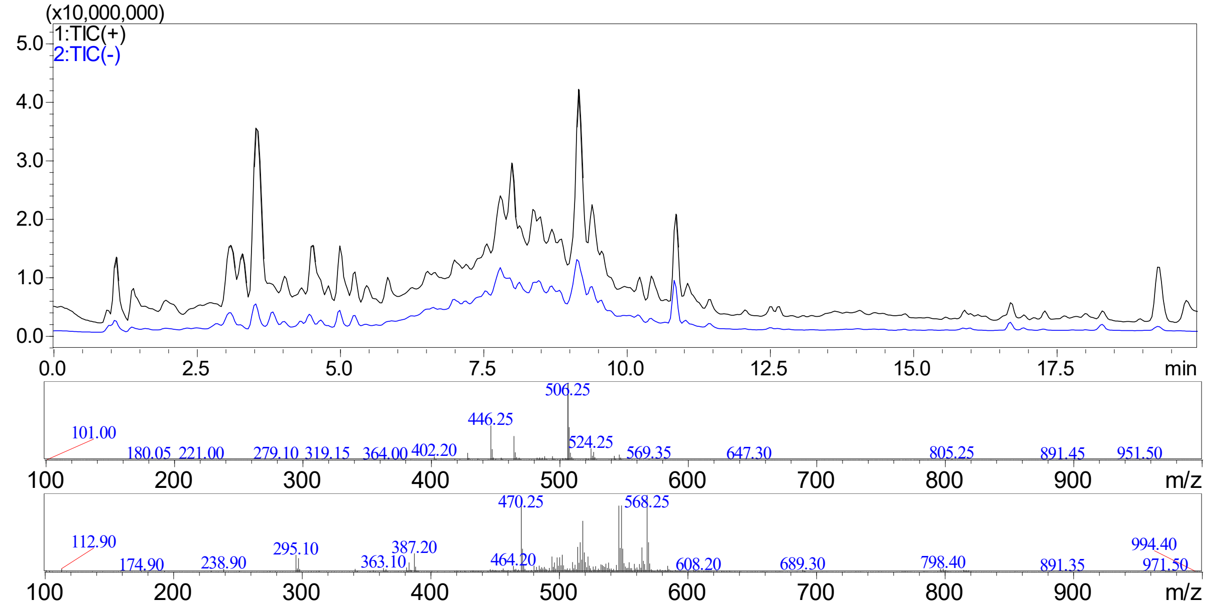


**Figure S31**: LCMS traces of *M. grisea* Δ*pyiD* + CHGG_01243 transformant LL-I-4-H6, showing production of pyrichalasin H at 9.1 mins (top trace), and the corresponding mass spectra in positive ion mode (middle) and negative ion mode (bottom).

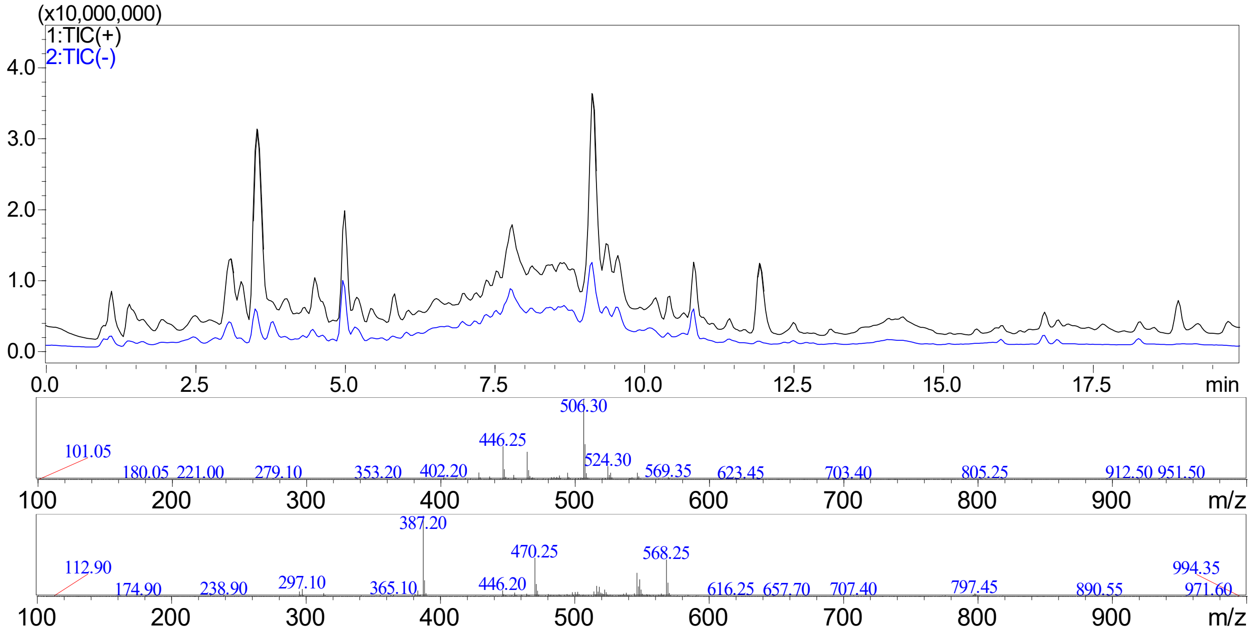


**Figure S32**: LCMS traces of *M. grisea* Δ*pyiD* + CHGG_01243 transformant LL-II-10-H3, showing production of pyrichalasin H at 9.1 mins (top trace), and the corresponding mass spectra in positive ion mode (middle) and negative ion mode (bottom).

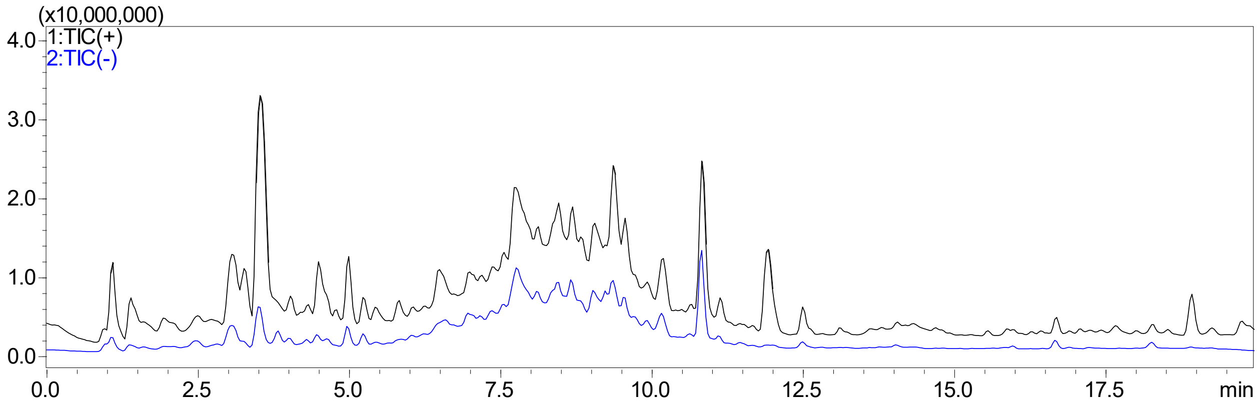


**Figure S33**: LCMS trace of *M. grisea* Δ*pyiD* + CHGG_01243 transformant LL-II-10-H4, showing no production of pyrichalasin H at 9.1 mins.

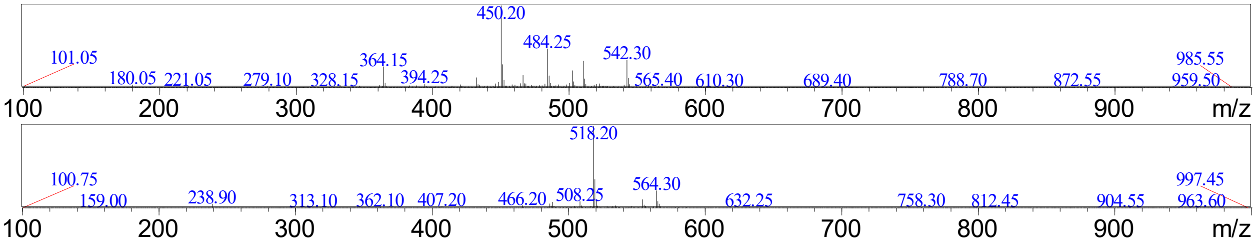


**Figure S34**: Mass spectra of *M. grisea* Δ*pyiD* + CHGG_01243 transformant LL-II-10-H4, showing production of cytochalasan "509" at Rt = 7.8 mins in positive ion mode (top) and negative ion mode (bottom).

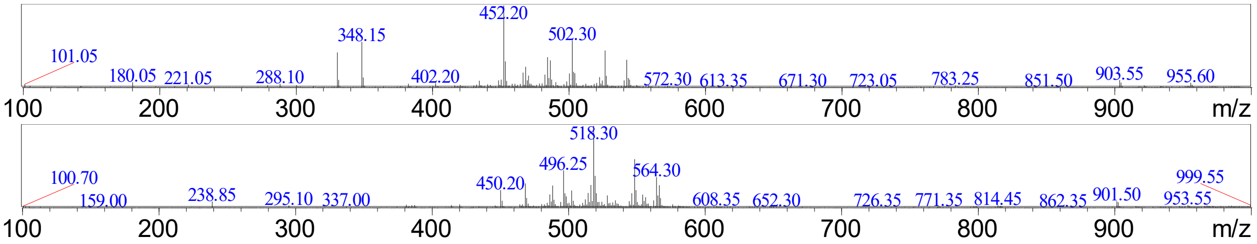


**Figure S35**: Mass spectra of *M. grisea* Δ*pyiD* + CHGG_01243 transformant LL-II-10-H4, showing production of cytochalasan "451" at Rt = 8.7 mins in positive ion mode (top) and negative ion mode (bottom).

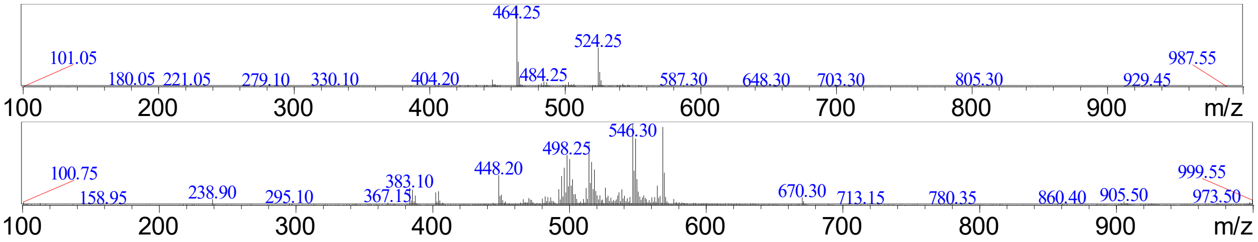


**Figure S36**: Mass spectra of *M. grisea* Δ*pyiD* + CHGG_01243 transformant LL-II-10-H4, showing production of cytochalasan "523" at Rt = 9.4 mins in positive ion mode (top) and negative ion mode (bottom).

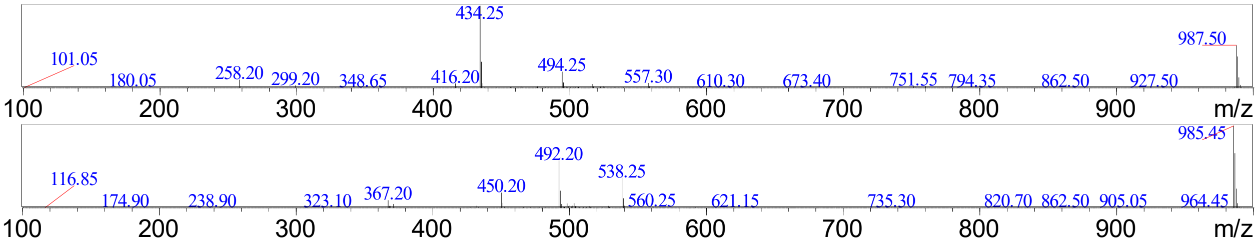


**Figure S37**: Mass spectra of *M. grisea* Δ*pyiD* + CHGG_01243 transformant LL-II-10-H4, showing production of cytochalasan "493" at Rt = 10.9 mins in positive ion mode (top) and negative ion mode (bottom).

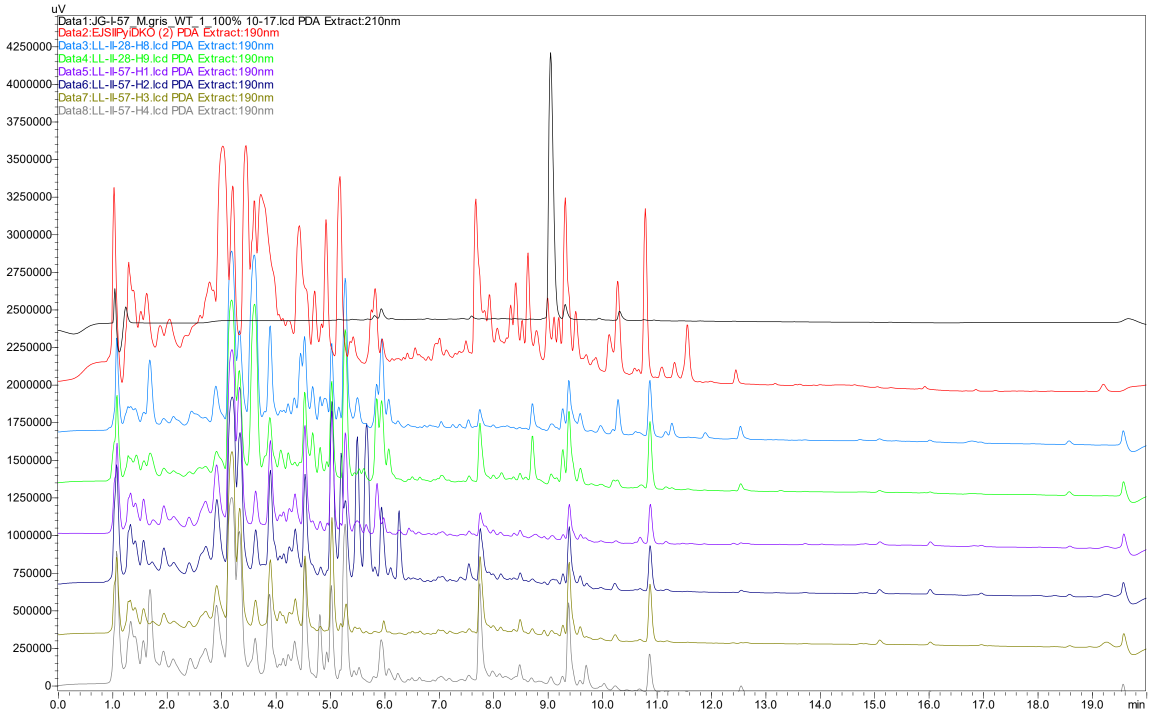


**Figure S38**: LCMS traces of *M. grisea* Δ*pyiD* + CcsD transformants, showing no evidence of new compounds compared to the control (EJSIIPyiDKO) or production of pyrichalasin H based on comparison to a standard (Rt = 9.1 mins).

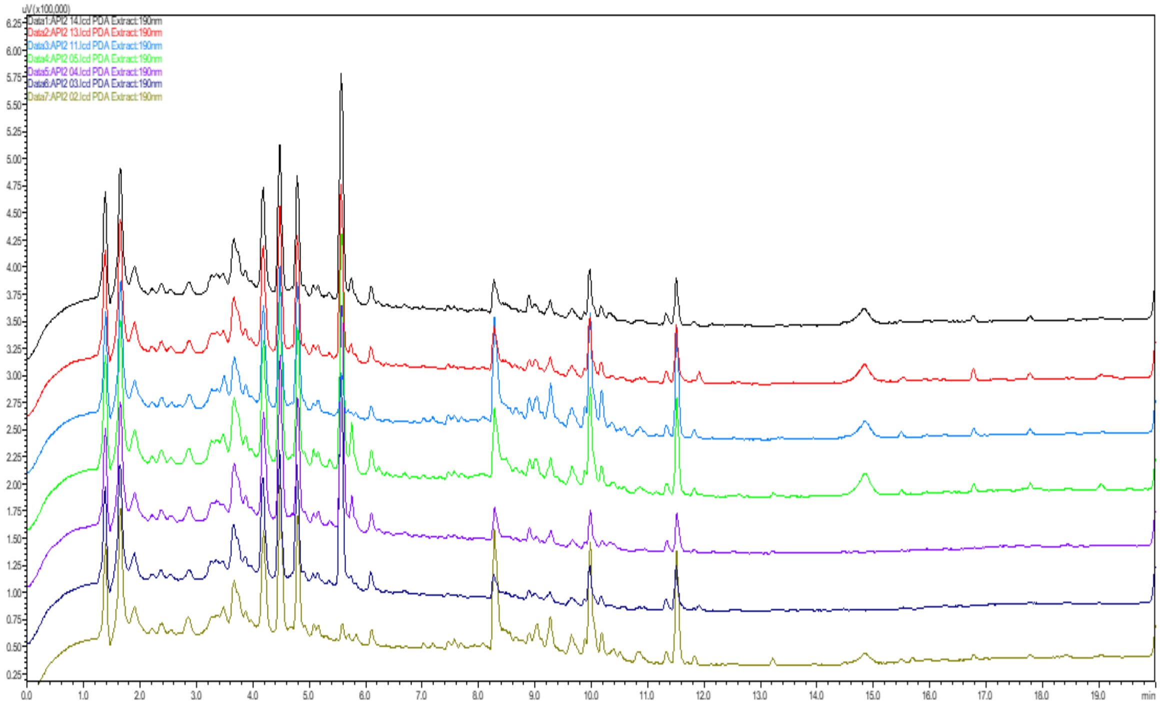


**Figure S39**: LCMS traces of *M. grisea* Δ*pyiD* + CcsD (revised sequence) transformants, demonstrating no new metabolites produced or restoration of pyrichalasin H production. (Note: Retention time shift of ~0.6 mins compared to Fig. S38 is due to working with a new column).

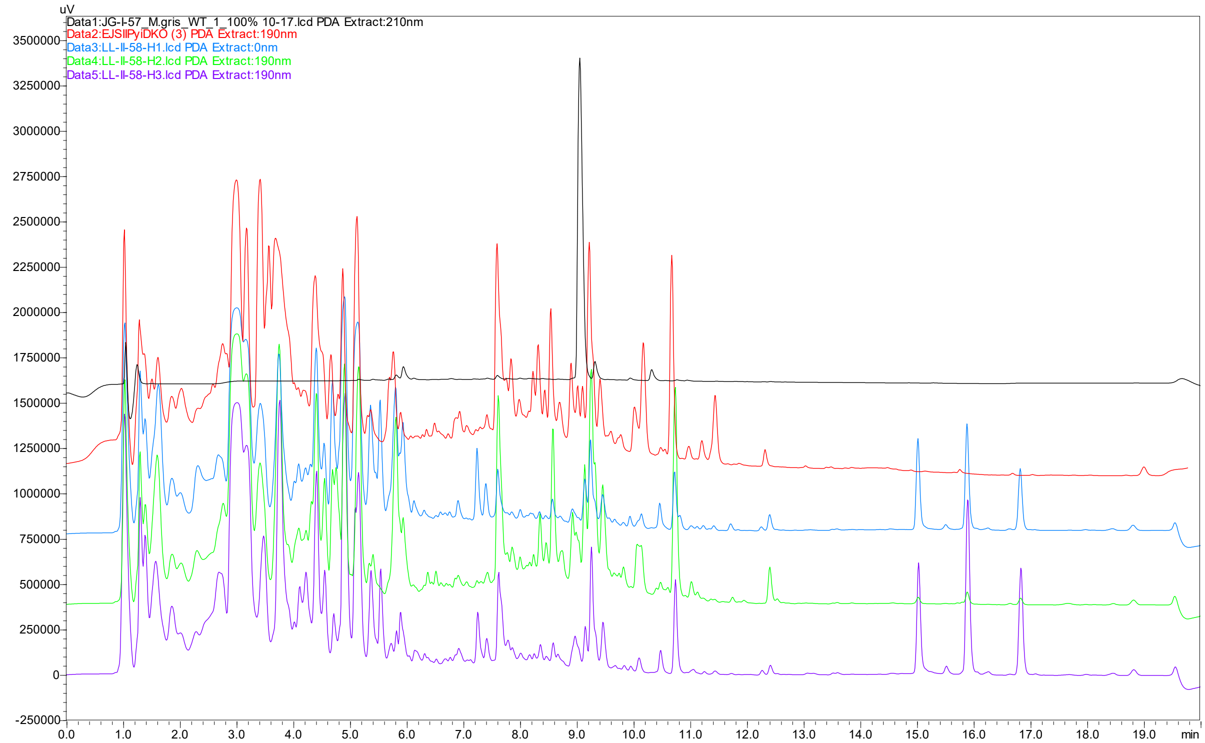


**Figure S40**: LCMS traces of *M. grisea* Δ*pyiD* + XsCYP2 transformants, showing no evidence of new compounds compared to the control (EJSIIPyiDKO) or production of pyrichalasin H based on comparison to a standard (Rt = 9.1 mins). Note: compounds eluting between 15 – 17 mins have mass ranges between 250 and 700 and are not related to cytochalasans.

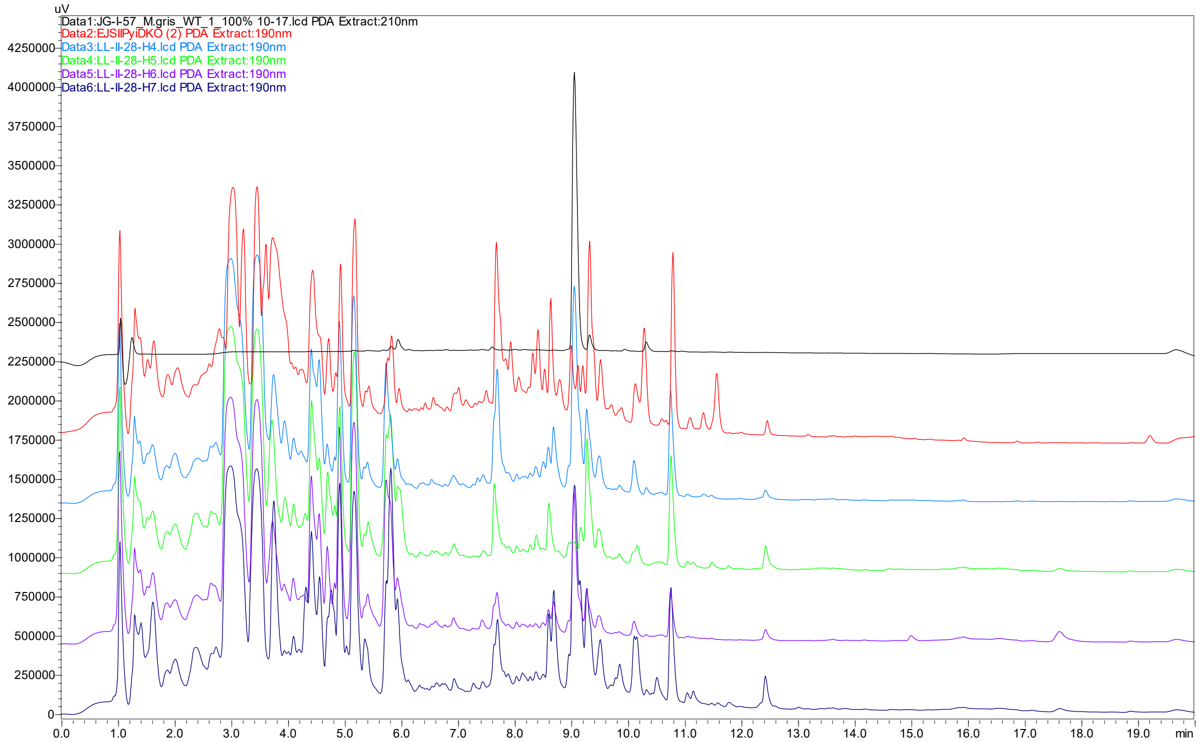


**Figure S41**: LCMS traces of *M. grisea* Δ*pyiD* + AhCYP2 transformants, showing production of pyrichalasin H based on comparison to a standard (Rt = 9.1 mins). Compared to the control strain (EJSIIPyiDKO) no new compounds are detected.

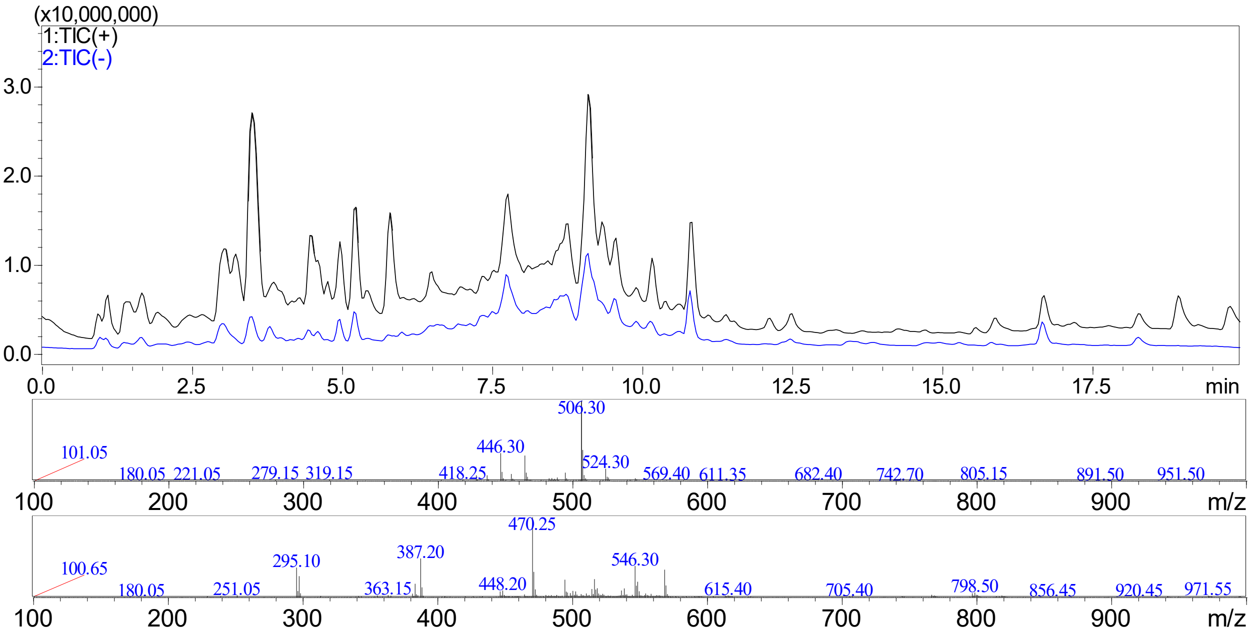


**Figure S42**: LCMS trace of *M. grisea* Δ*pyiD* + AhCYP2 transformant LL-II-28-H4, showing production of pyrichalasin H at Rt = 9.1 mins (top trace) and the corresponding mass spectra in positive ion mode (middle) and negative ion mode (bottom).

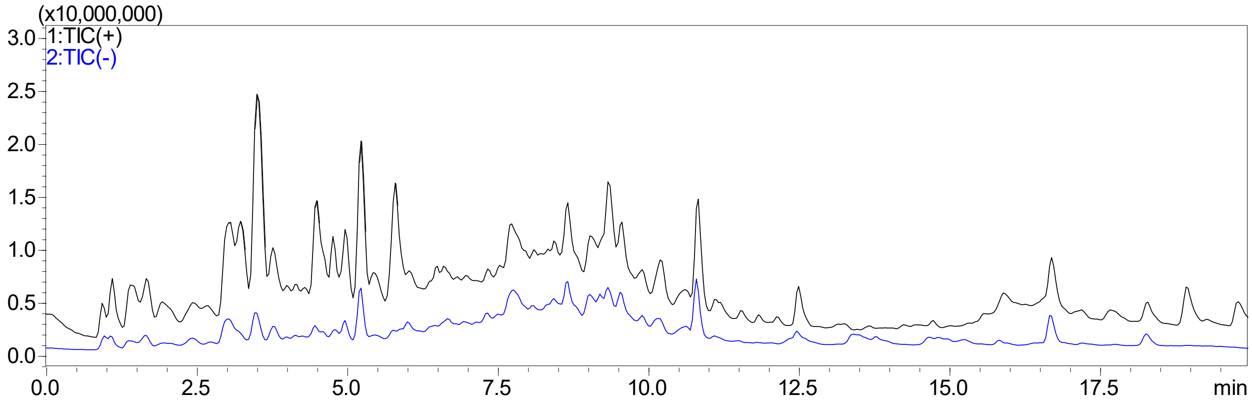


**Figure S43**: LCMS trace of *M. grisea* Δ*pyiD* + AhCYP2 transformant LL-II-28-H5, showing no production of pyrichalasin H at Rt = 9.1 mins.

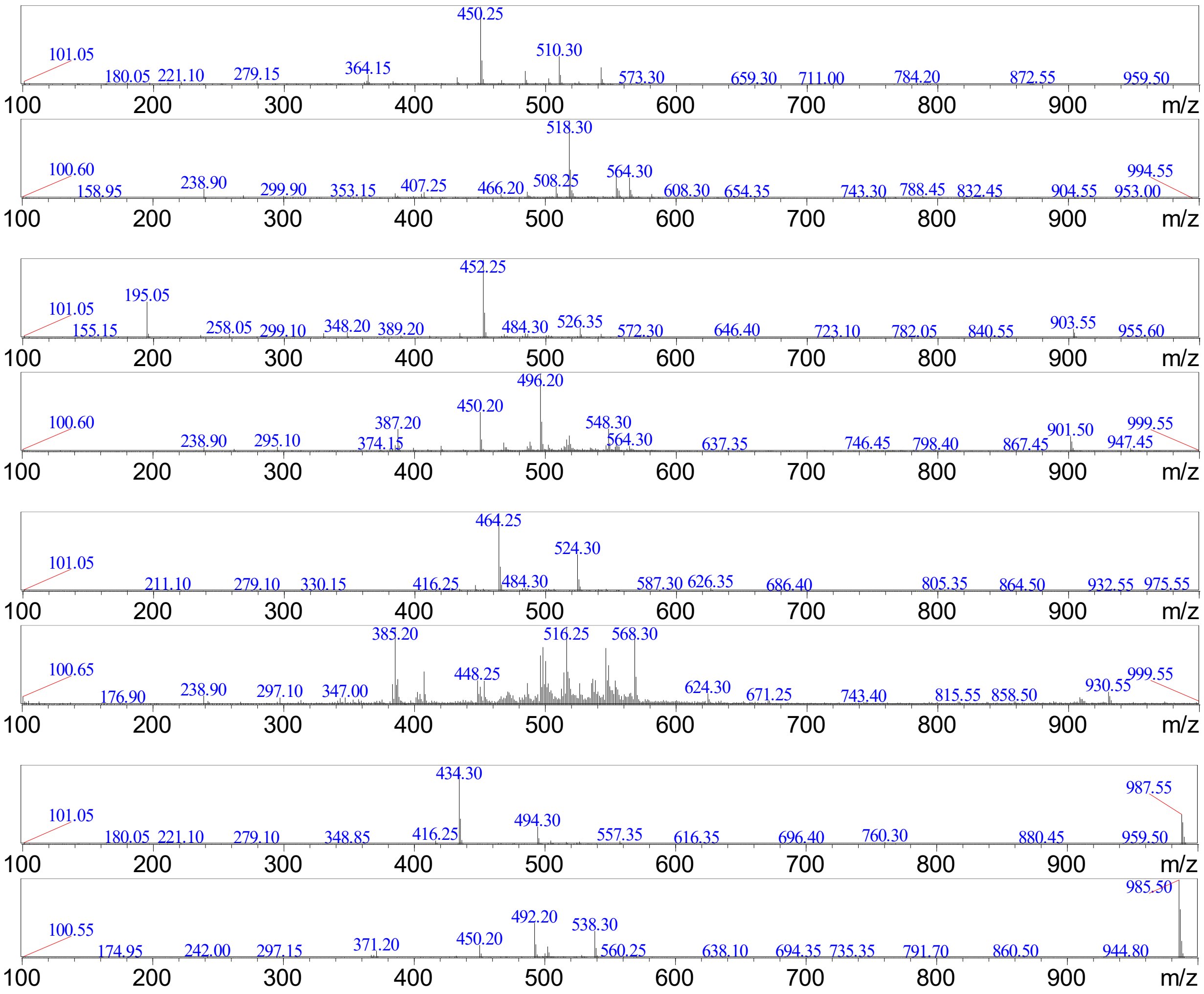


**Figure S44**: Mass spectra of *M. grisea* Δ*pyiD* + AhCYP2 transformant LL-II-28-H5, showing production of cytochalasans '509', '451', '523' and '493' at 7.8, 8.7, 9.4, and 10.9 mins respectively. Mass spectra are shown in pairs (positive then negative ion mode).

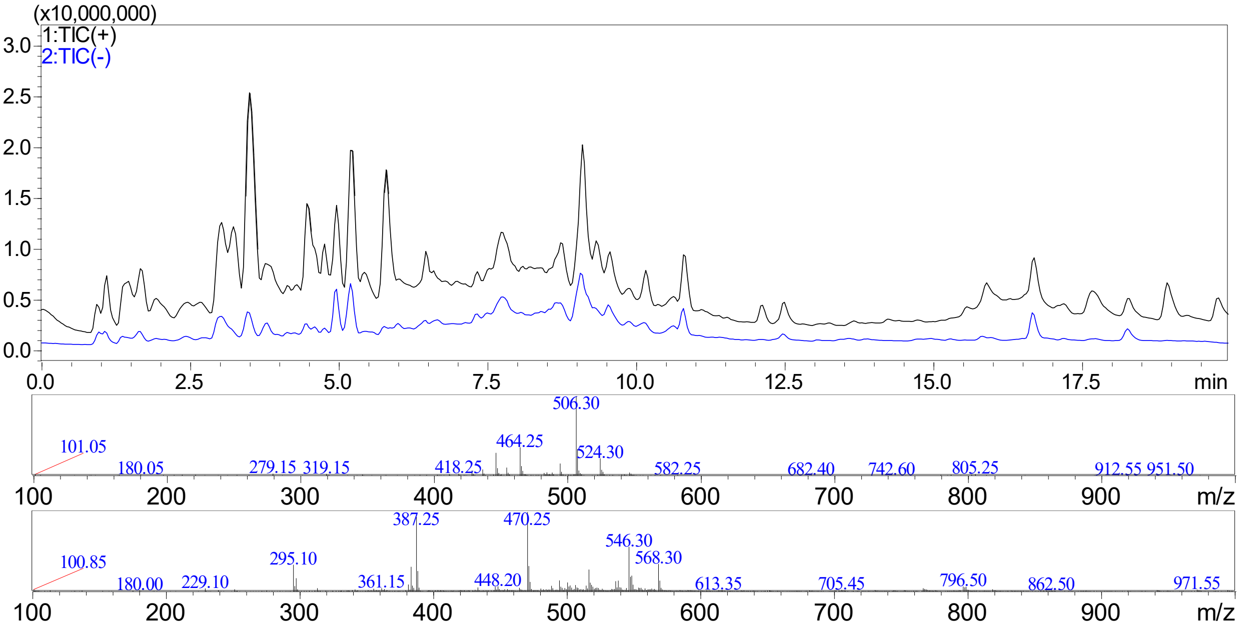


**Figure S45**: LCMS trace of *M. grisea* Δ*pyiD* + AhCYP2 transformant LL-II-28-H6, showing production of pyrichalasin H at Rt = 9.1 mins (top trace) and the corresponding mass spectra in positive ion mode (middle) and negative ion mode (bottom).

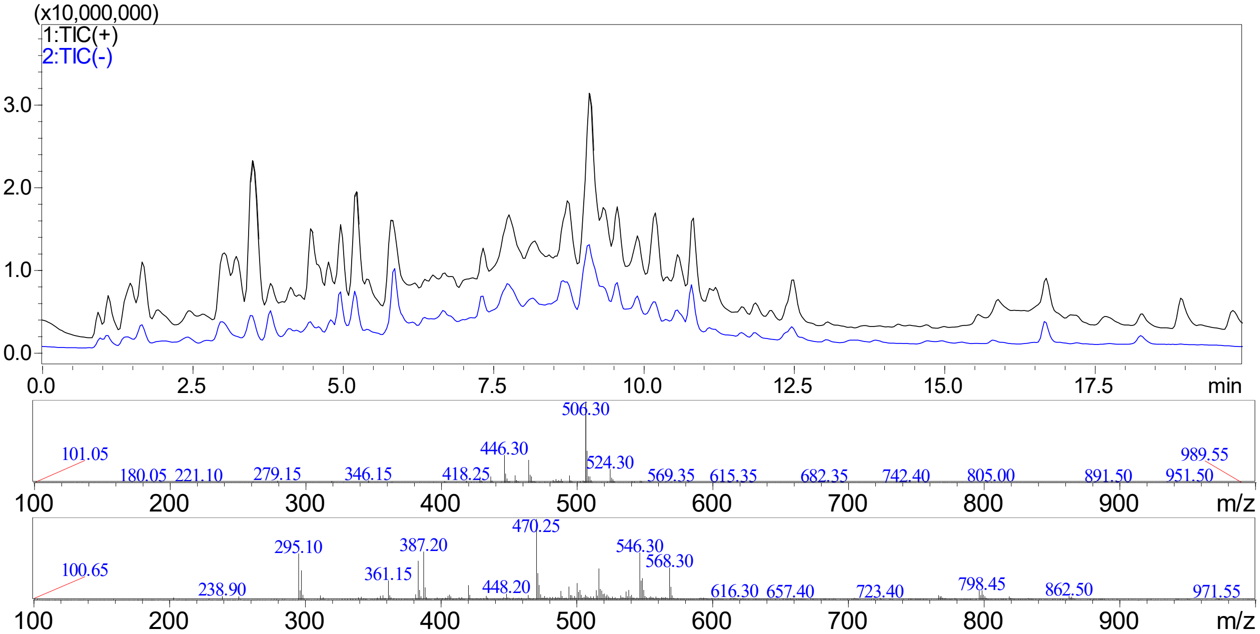


**Figure S46**: LCMS trace of *M. grisea* Δ*pyiD* + AhCYP2 transformant LL-II-28-H7, showing production of pyrichalasin H at Rt = 9.1 mins (top trace) and the corresponding mass spectra in positive ion mode (middle) and negative ion mode (bottom).

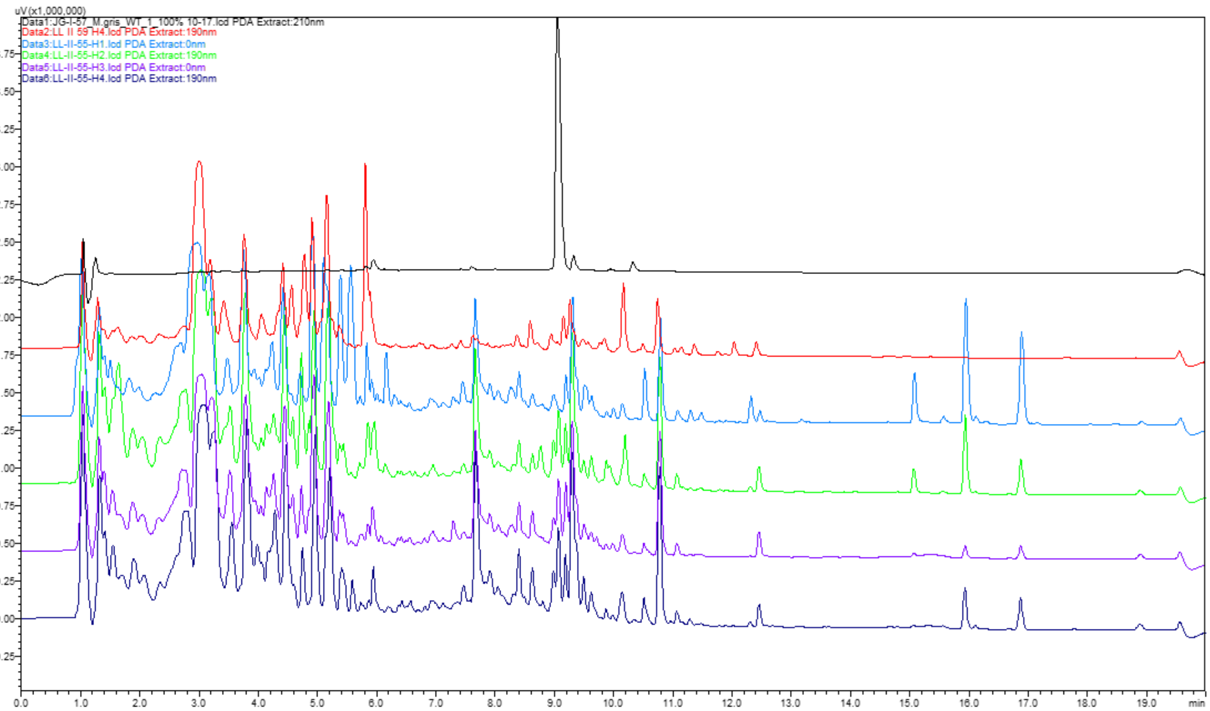


**Figure S47**: LCMS traces of *M. grisea* Δ*pyiD* + MrCYP1 transformants, showing no evidence of new compounds compared to control (LL II 59 H4) and no production of pyrichalasin H based on comparison to a standard (Rt = 9.1 mins). Note: compounds eluting between 15 – 17 mins have mass ranges between 250 and 700 and are not related to cytochalasans.

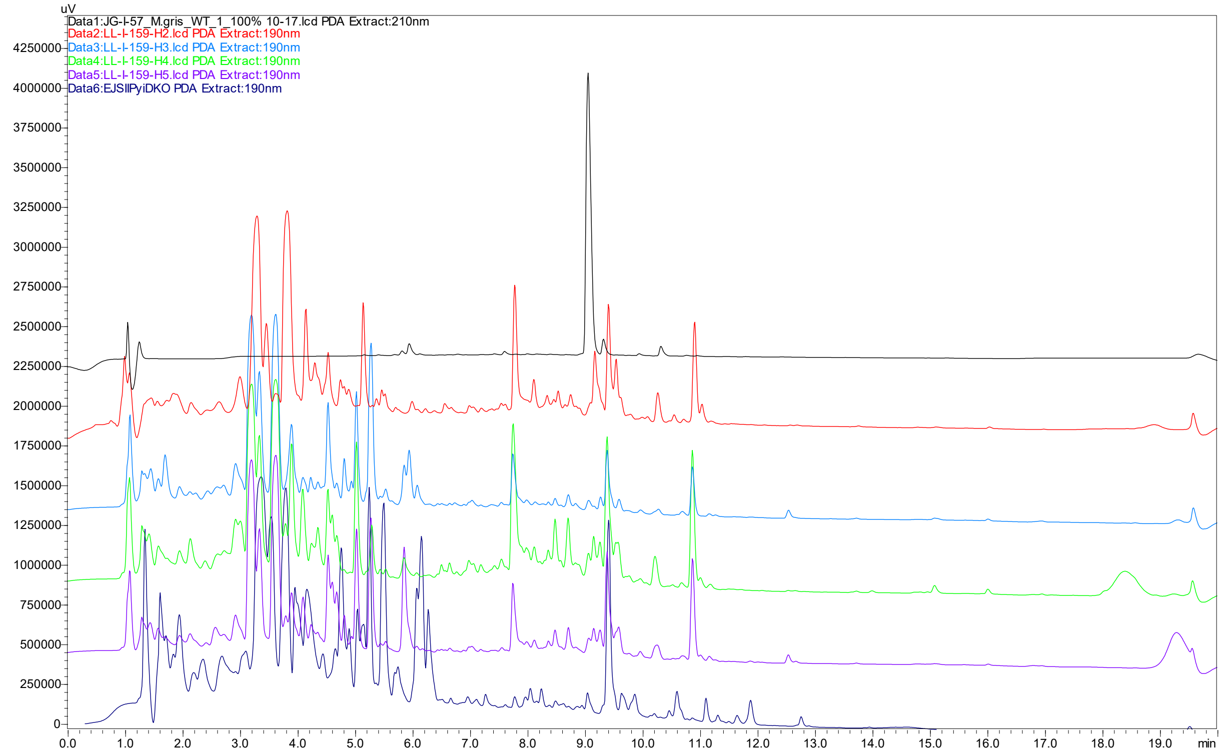


**Figure S48**: LCMS traces of *M. grisea* Δ*pyiD* + CsCYP1 transformants, showing no production of pyrichalasin H based on comparison to a standard (Rt = 9.1 mins). Compared to the control strain (EJSIIPyiDKO) no new compounds are detected.

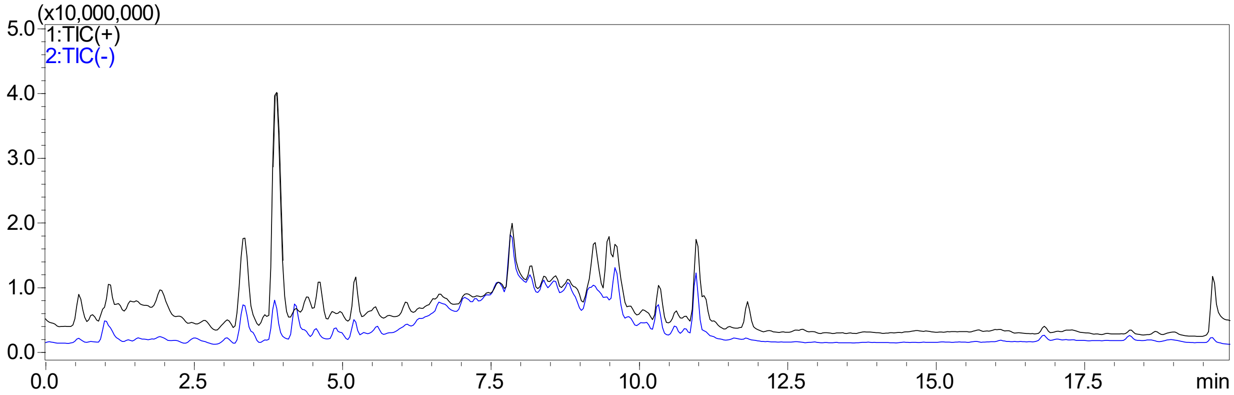


**Figure S49**: LCMS trace of *M. grisea* Δ*pyiD* + CsCYP1 transformant LL-I-159-H2, showing no production of pyrichalasin H at Rt = 9.1 mins.

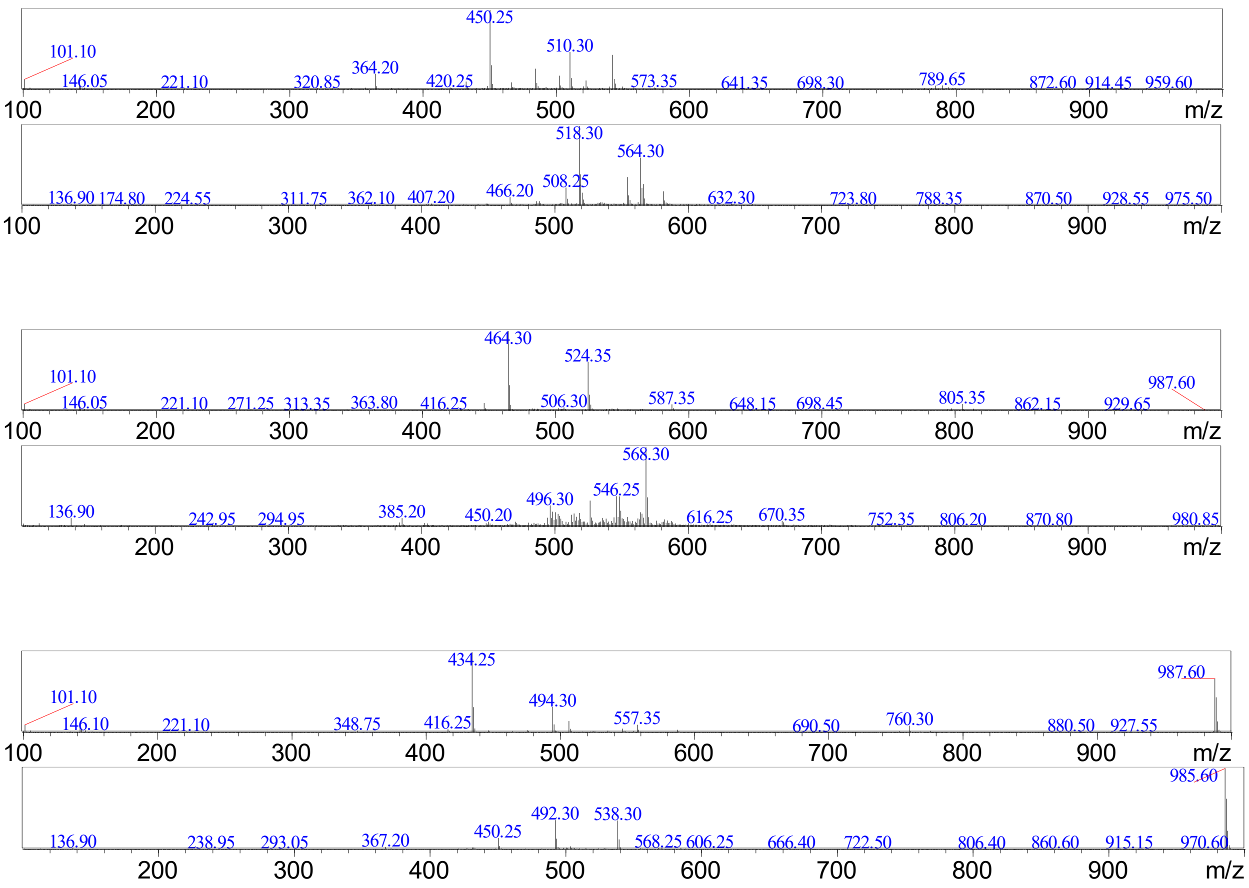


**Figure S50**: Mass spectra of *M. grisea* Δ*pyiD* + CsCYP1 transformant LL-I-159-H2, showing production of cytochalasans '509', '523' and '493' at 7.8, 9.4, and 10.9 mins respectively. Mass spectra are shown in pairs (positive then negative ion mode).

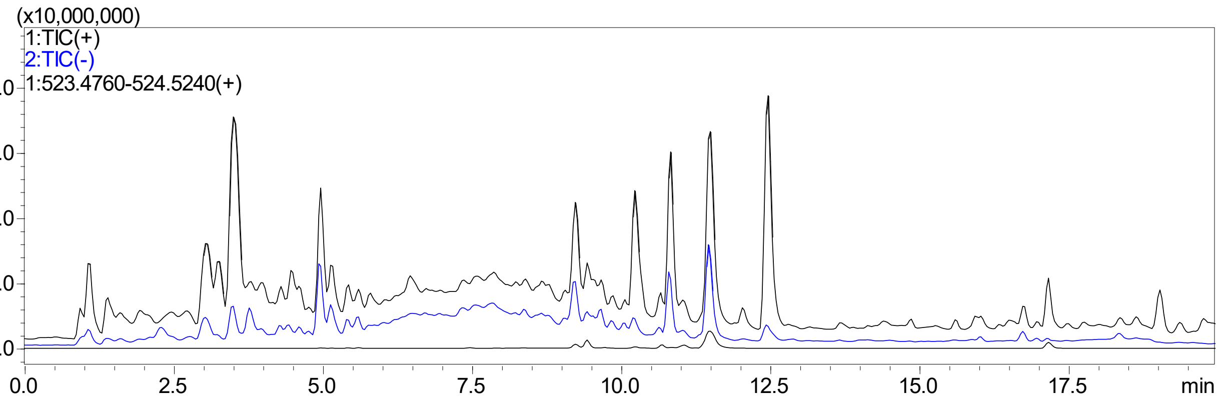


**Figure S51**: LCMS trace of *M. grisea* Δ*pyiG* host strain (LL-II-4-H8) showing no production of pyrichalasin H at Rt = 9.1. Instead cytochalasans with *m/z* = 539, 523 / 493, 507, 523, and 493 are observed at Rt = 9.2 – 12.5 mins respectively.

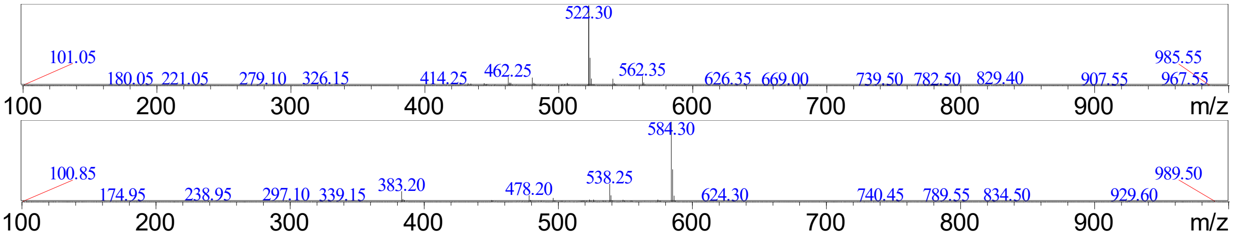


**Figure S52**: Mass spectra of *M. grisea* Δ*pyiG* host strain showing no production of pyrichalasin H at Rt = 9.1. Instead cytochalasan with *m/z* = 539 is observed at 9.2 mins in positive ion mode (top) and negative ion mode (bottom).

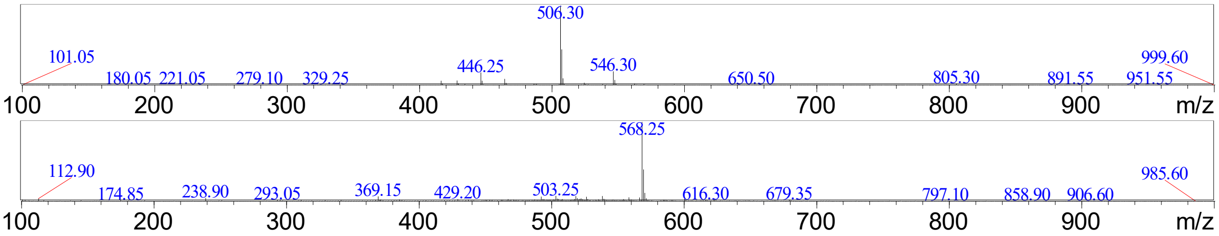


**Figure S53**: Mass spectra of *M. grisea* Δ*pyiG* host strain showing no production of pyrichalasin H at Rt = 9.1. Instead cytochalasan with *m/z* = 523 is observed at 10.2 mins in positive ion mode (top) and negative ion mode (bottom).

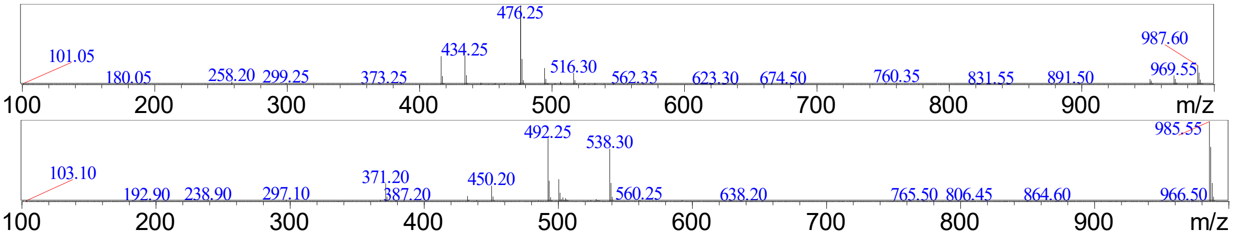


**Figure S54**: Mass spectra of *M. grisea* Δ*pyiG* host strain showing no production of pyrichalasin H at Rt = 9.1. Instead cytochalasan with *m/z* = 507 is observed at 10.8 mins in positive ion mode (top) and negative ion mode (bottom).

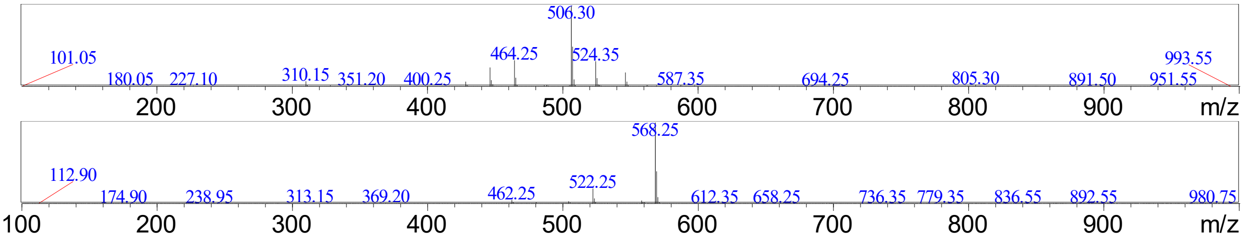


**Figure S55**: Mass spectra of *M. grisea* Δ*pyiG* host strain showing no production of pyrichalasin H at Rt = 9.1. Instead cytochalasan with *m/z* = 523 is observed at 11.5 mins in positive ion mode (top) and negative ion mode (bottom).

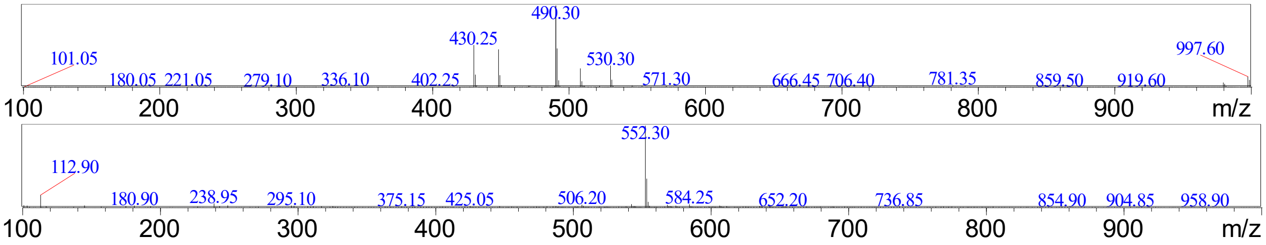


**Figure S56**: Mass spectra of *M. grisea* Δ*pyiG* host strain showing no production of pyrichalasin H at Rt = 9.1. Instead cytochalasan with *m/z* = 493 is observed at 12.5 mins in positive ion mode (top) and negative ion mode (bottom).

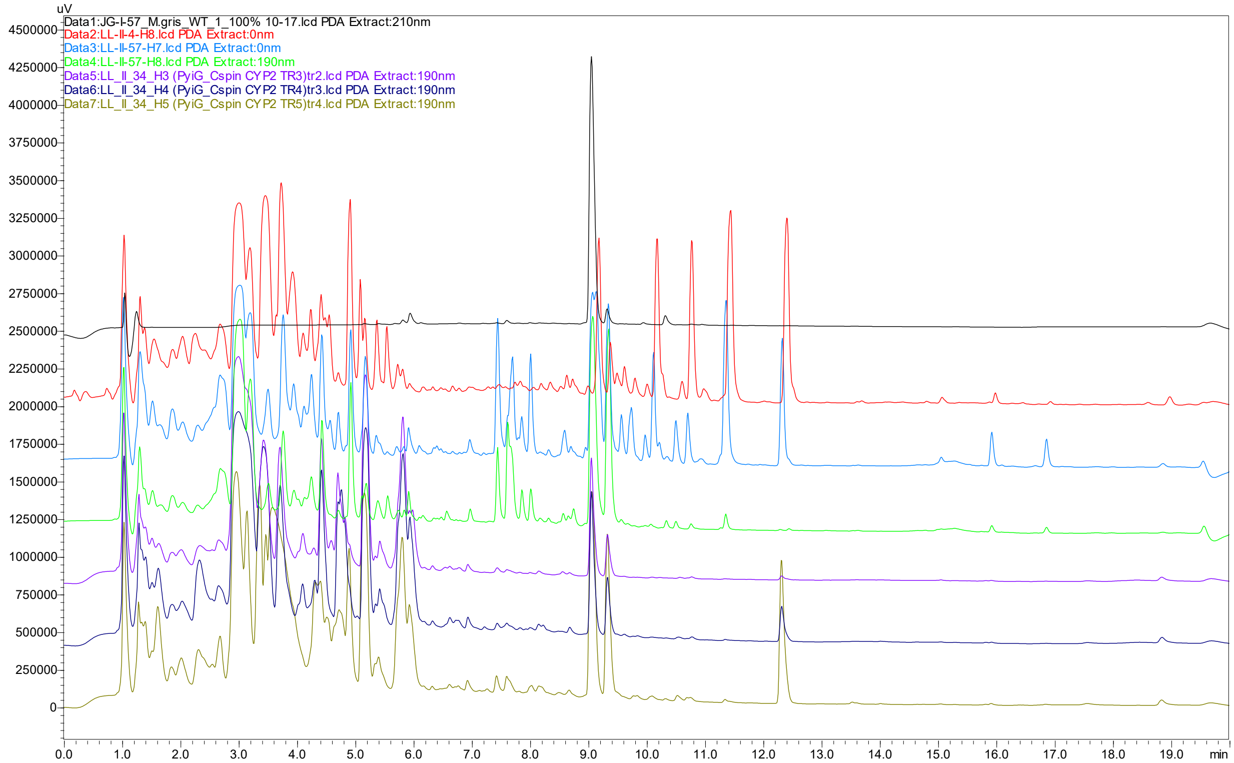


**Figure S57**: LCMS traces of *M. grisea* Δ*pyiG* + CsCYP2 transformants, showing production of pyrichalasin H based on comparison to a standard (Rt = 9.1 mins). Compared to the control strain (LL-II-4-H6) no new compounds are detected.

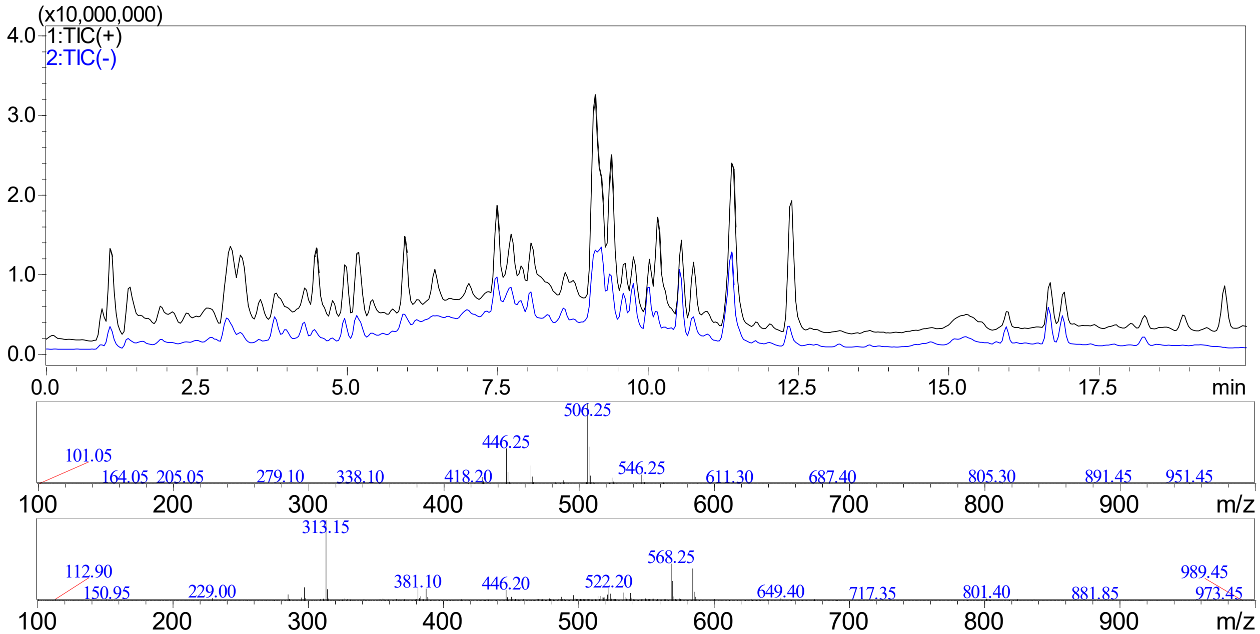


**Figure S58**: LCMS trace of *M. grisea* Δ*pyiG* + CsCYP2 transformant LL-II-57-H7, showing production of pyrichalasin H at Rt = 9.1 mins (top chromatogram) and the corresponding mass spectra in positive ion mode (middle) and negative ion mode (bottom).

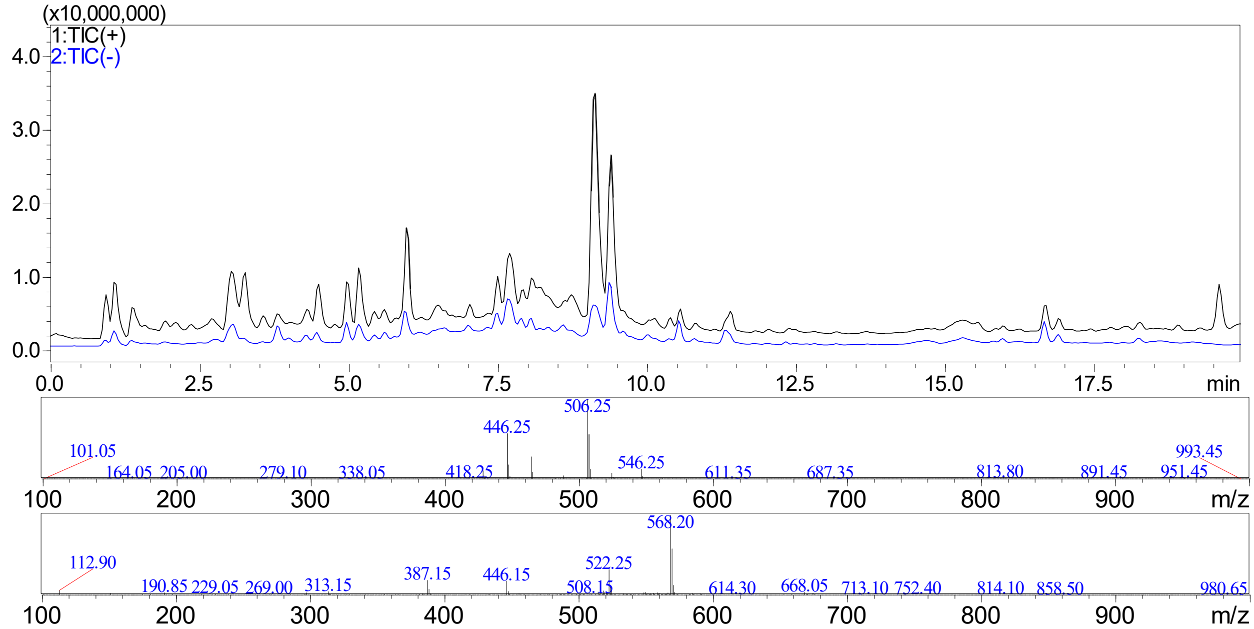


**Figure S59**: LCMS trace of *M. grisea* Δ*pyiG* + CsCYP2 transformant LL-II-57-H8, showing production of pyrichalasin H at Rt = 9.1 mins (top trace) and the corresponding mass spectra in positive ion mode (middle) and negative ion mode (bottom).

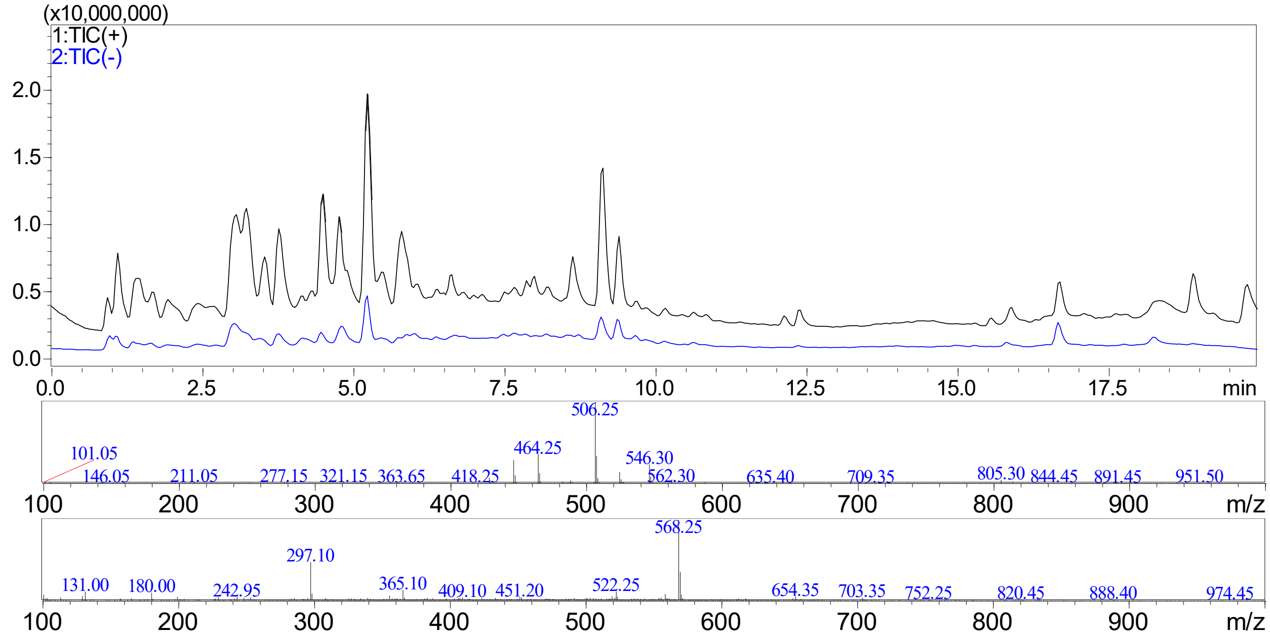


**Figure S60**: LCMS trace of *M. grisea* Δ*pyiG* + CsCYP2 transformant LL-II-34-H3, showing production of pyrichalasin H at Rt = 9.1 mins (top chromatogram) and the corresponding mass spectra in positive ion mode (middle) and negative ion mode (bottom).

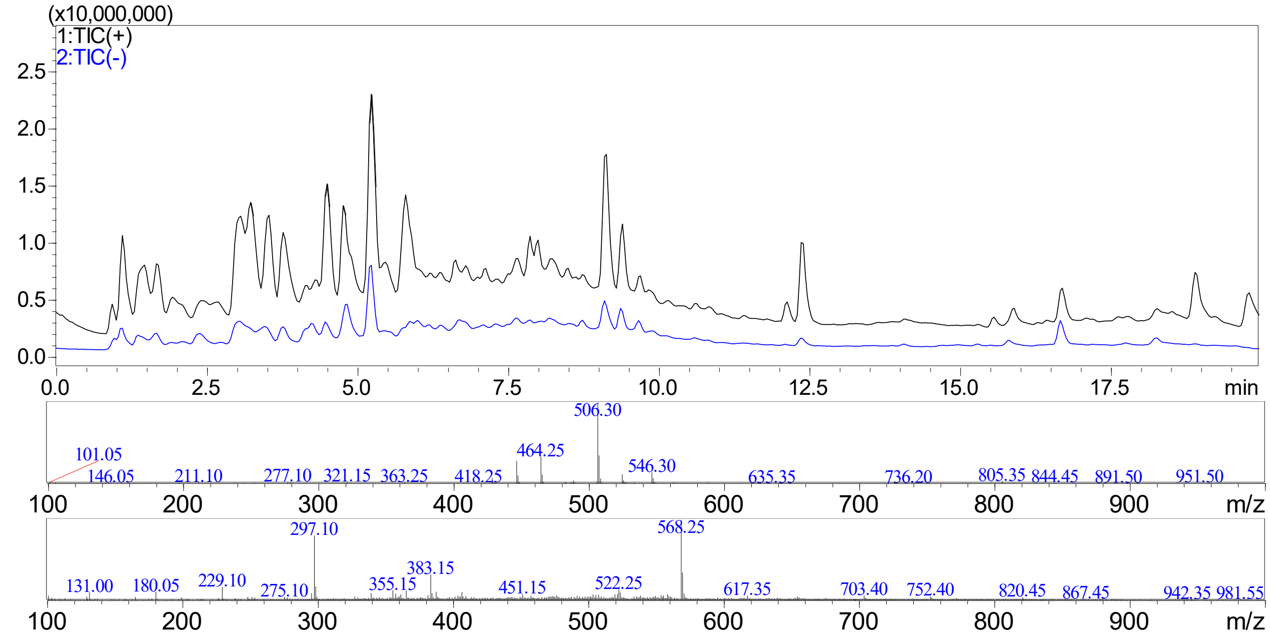


**Figure S61**: LCMS trace of *M. grisea* Δ*pyiG* + CsCYP2 transformant LL-II-34-H4, showing production of pyrichalasin H at Rt = 9.1 mins (top chromatogram) and the corresponding mass spectra in positive ion mode (middle) and negative ion mode (bottom).

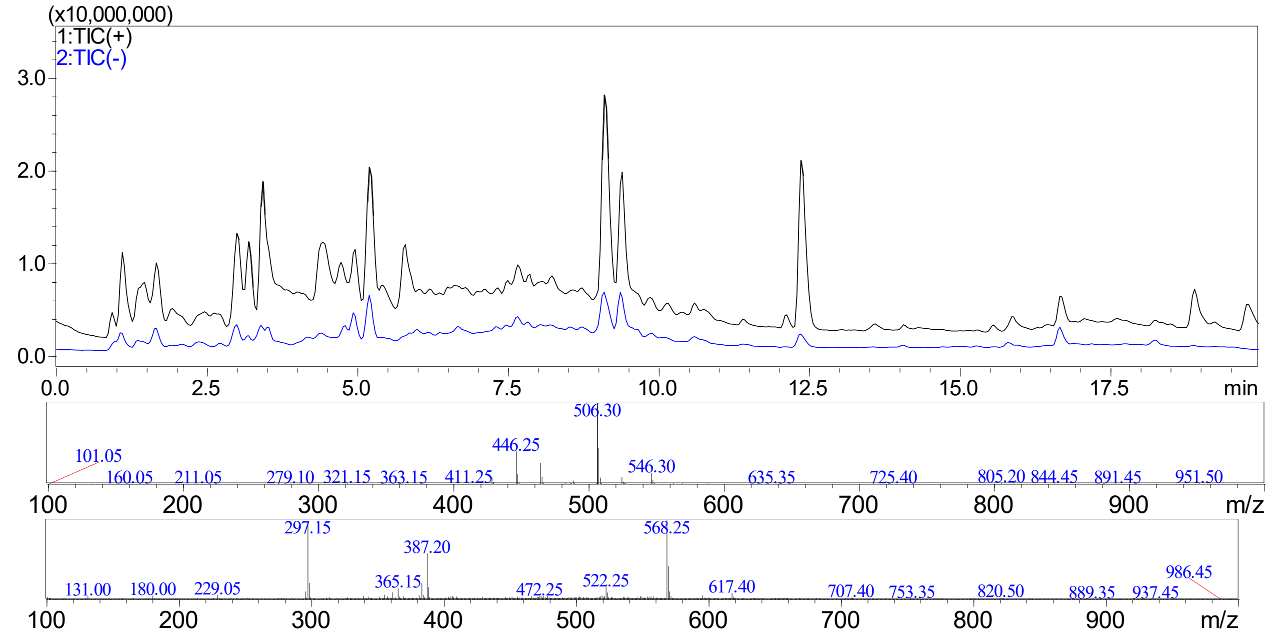


**Figure S62**: LCMS trace of *M. grisea* Δ*pyiG* + CsCYP2 transformant LL-II-34-H5, showing production of pyrichalasin H at Rt = 9.1 mins (top chromatogram) and the corresponding mass spectra in positive ion mode (middle) and negative ion mode (bottom).

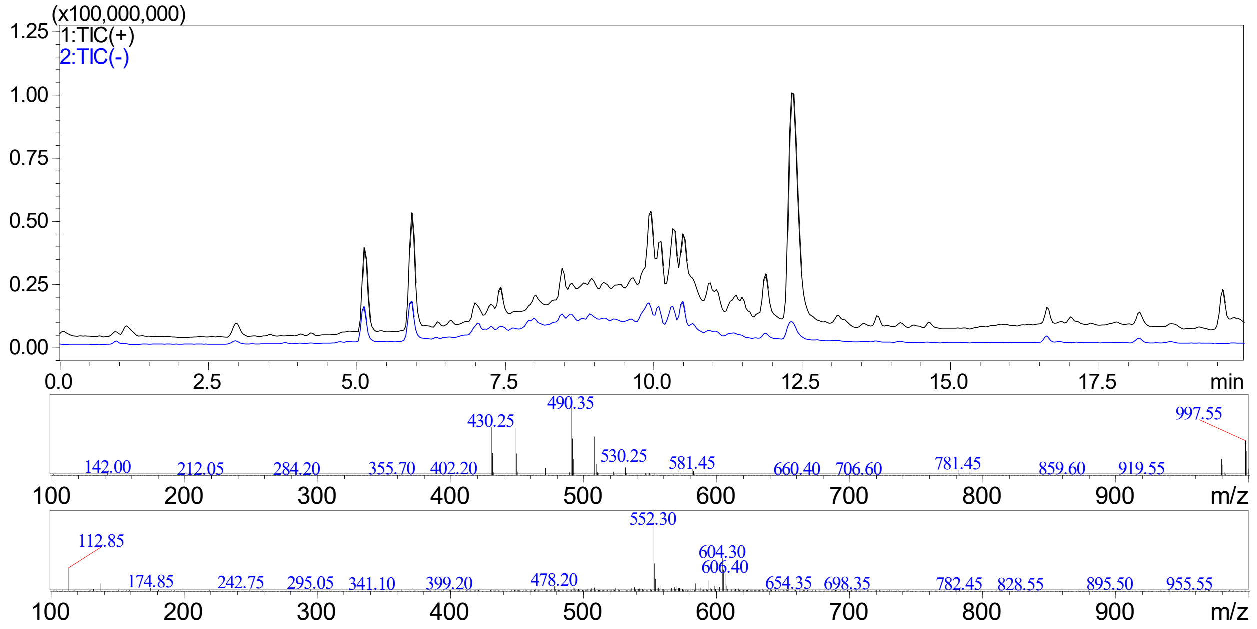


**Figure S63**: LCMS trace of semi-pure 7-desmethylpyrichalasin H at Rt = 12.5 mins (top chromatogram) and the corresponding mass spectra in positive ion mode (middle) and negative ion mode (bottom).

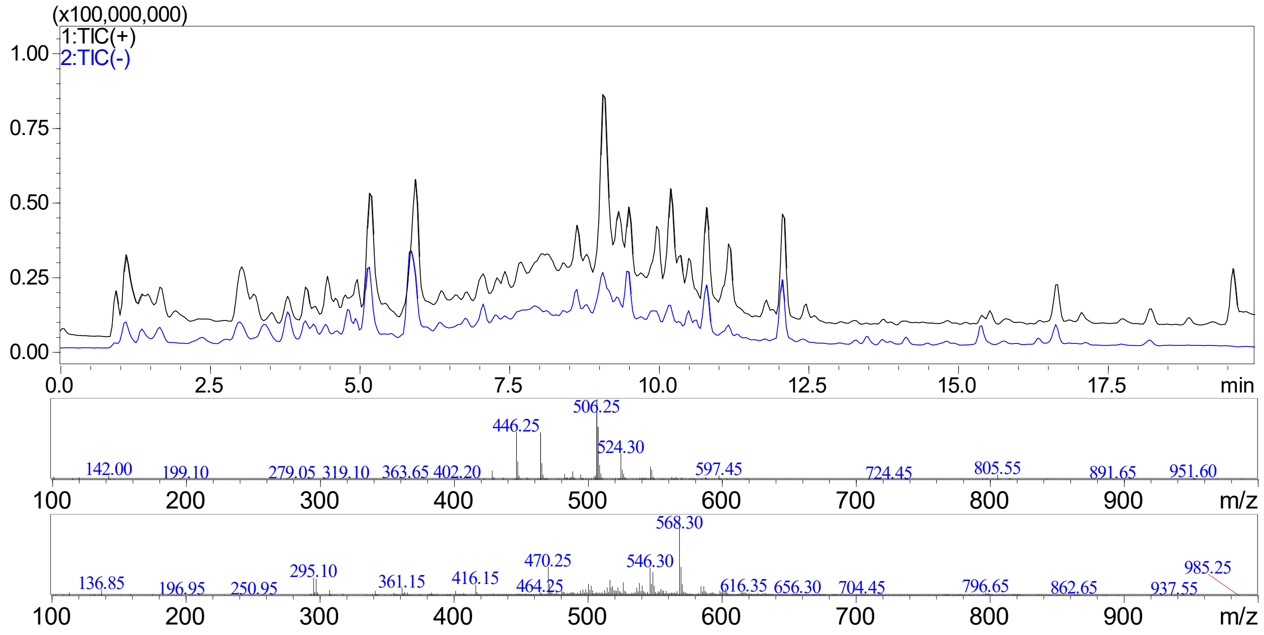


**Figure S64**: LCMS trace of *M. grisea* Δ*pyiD* + 7-desmethylpyrichalasin H (LL-II-59-H2), showing production of pyrichalasin H at Rt = 9.1 mins (top chromatogram) and the corresponding mass spectra in positive ion mode (middle) and negative ion mode (bottom).

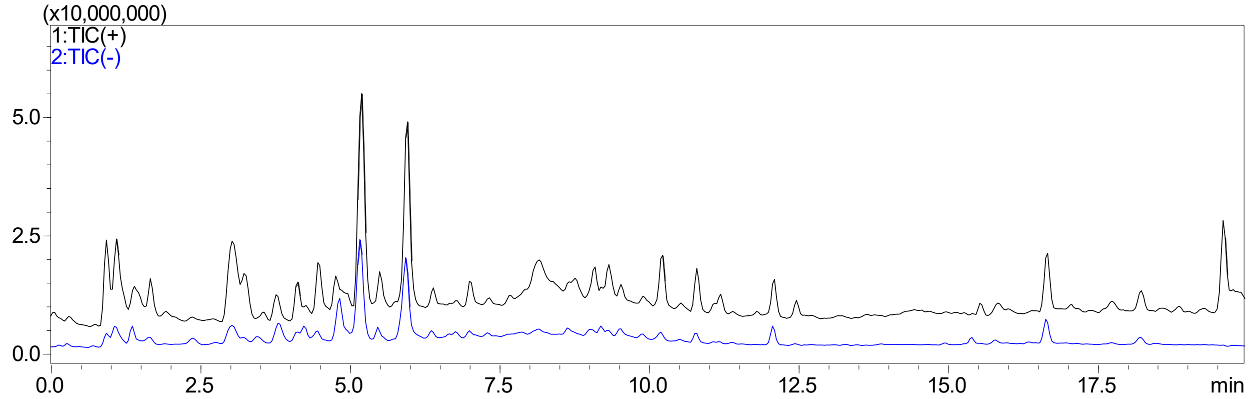


**Figure S65**: LCMS trace of *M. grisea* Δ*pyiD* + DMSO control (LL-II-59-H3), showing no production of pyrichalasin H at Rt = 9.1 mins.

#### 3. NMR Data

| Position | $\delta_H$ (ppm) | Multiplicity |
|---|---|---|
| 1 | - | - |
| 2 | 5.55 | s |
| 3 | 3.21 | ddd |
| 4 | 2.11 | dd |
| 5 | 2.79 | m |
| 6 | - | - |
| 7 | 3.83 | dd |
| 8 | 2.95 | dd |
| 9 | - | - |
| 10a/b | 2.59 / 2.80 | dd / dd |
| 11 | 0.99 | d |
| 12a/b | 5.12 / 5.35 | brs / brs |
| 13 | 5.77 | ddd |
| 14 | 5.40 | ddd |
| 15a/b | 1.82 / 2.05 | m / m |
| 16 | 1.83 | m |
| 17a/b | 1.57 / 1.87 | dd / m |
| 18 | - | - |
| 19 | 5.54 | dd |
| 20 | 5.90 | dd |
| 21 | 5.54 | dd |
| 22 | 1.05 | d |
| 23 | 1.35 | s |
| 1' | - | - |
| 2',6' | 7.05 | d |
| 3',5' | 6.90 | d |
| 4' | - | - |
| 24 | - | - |
| 25 | 2.24 | s |
| 26 | 3.81 | s |

**Table S9**: 1H NMR data of pyrichalasin H recorded at 400 MHz in $CDCl_3$. Values are in agreement with published data. [8]

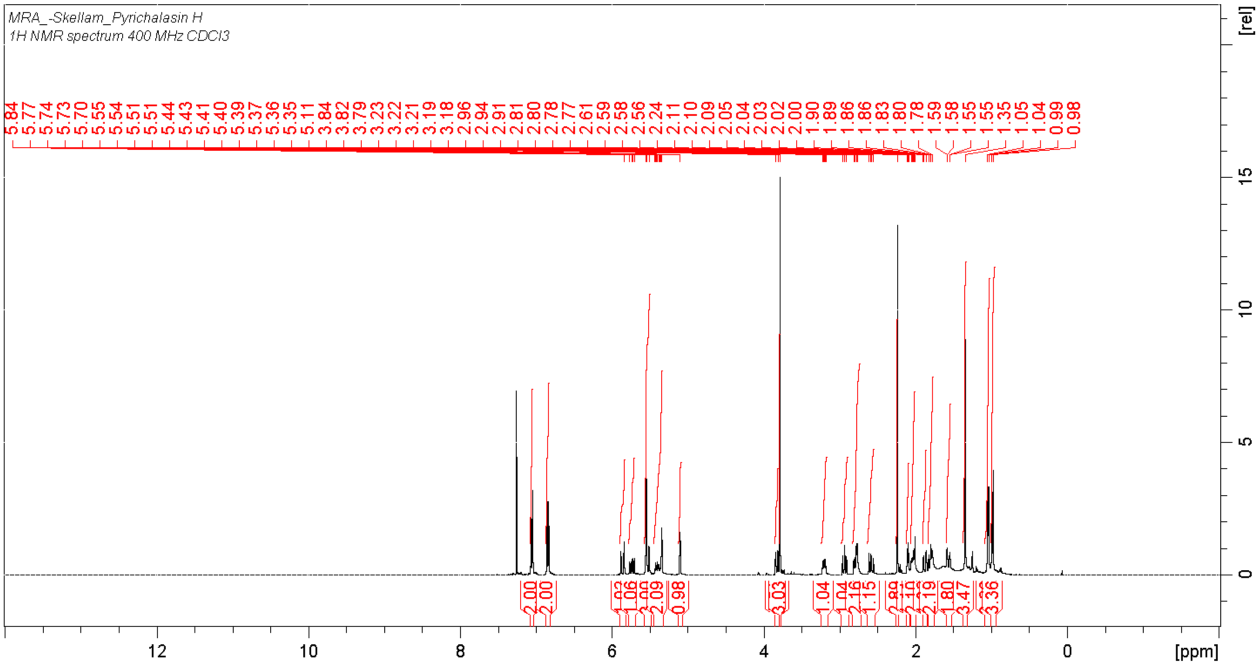

**Figure S66**. [1]H NMR spectrum (400 MHz, $CDCl_3$) of pyrichalasin H.

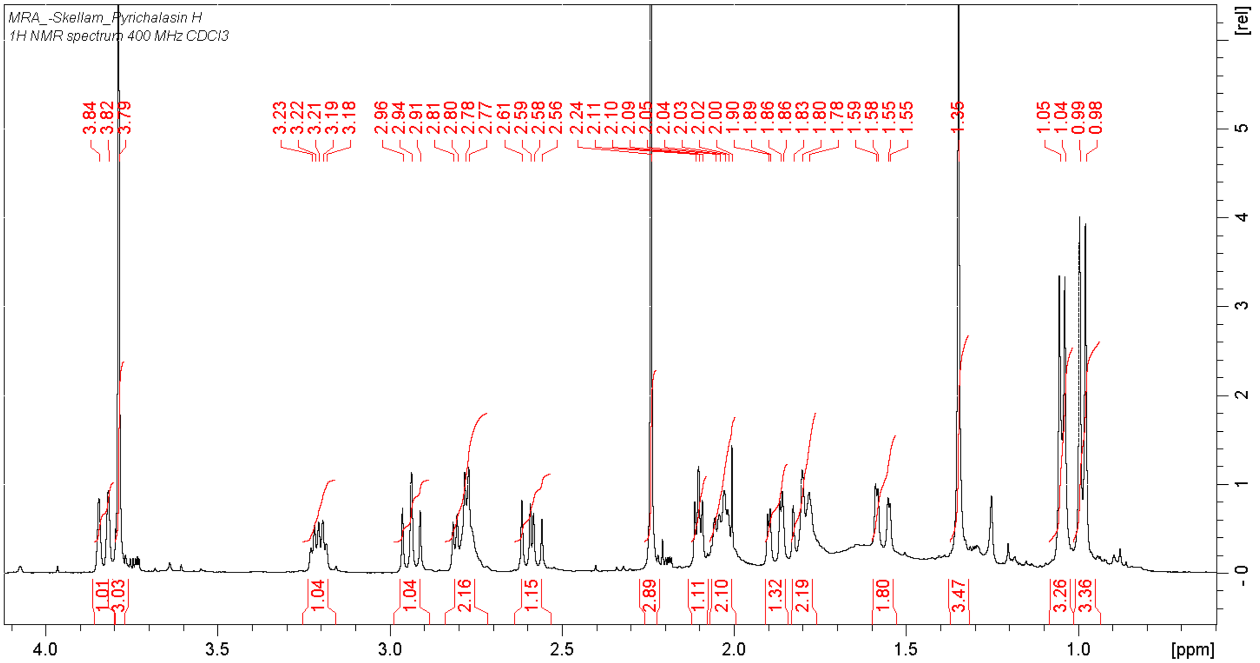

**Figure S67**. Expansion of [1]H NMR spectrum (400 MHz, $CDCl_3$) of pyrichalasin H.

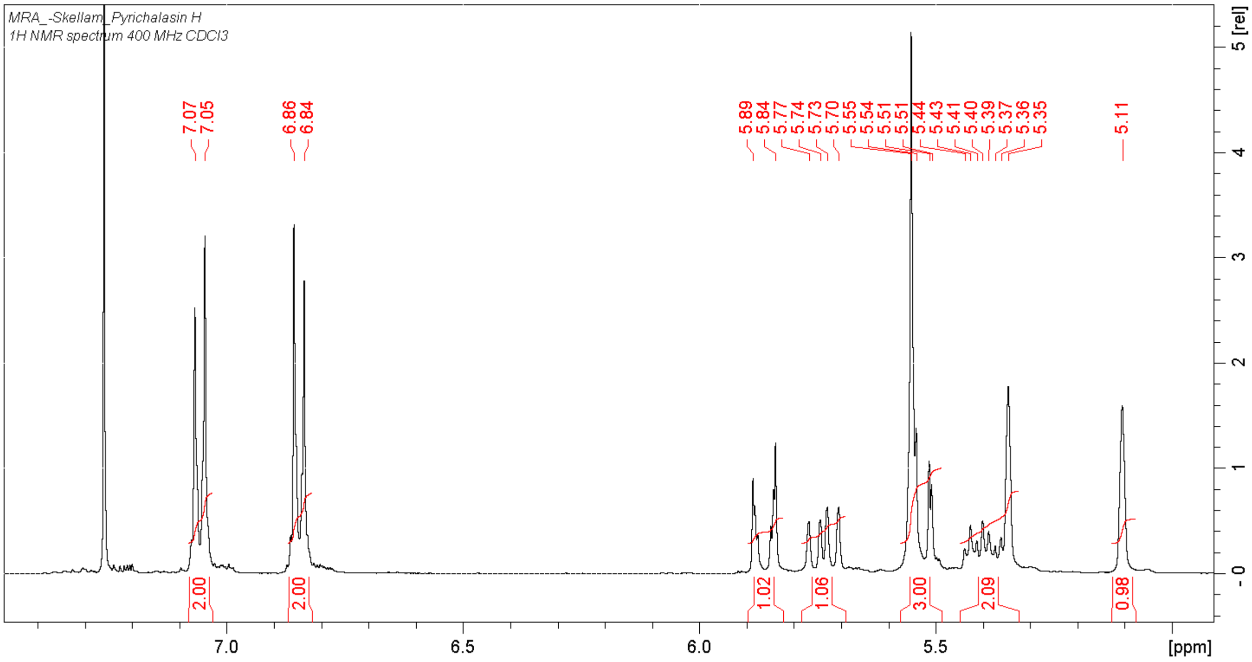


**Figure S68.** Expansion of [1]H NMR spectrum (400 MHz, $CDCl_3$) of pyrichalasin H.

| Position | $\delta_H$ (ppm) | Multiplicity | $\delta_C$ (ppm) |
|---|---|---|---|
| 1 | - | - | 175 |
| 2 | 5.55 | s | - |
| 3 | 3.10 | ddd | 56 |
| 4 | 2.13 | dd | 54 |
| 5 | 2.52 | m | 35 |
| 6 | - | - | 138 |
| 7 | 5.40 | brs | 128 |
| 8 | 3.2 | dd | 43 |
| 9 | - | - | 56 |
| 10a/b | 2.52 / 2.82 | dd / dd | 45 |
| 11 | 1.11 | d | 14 |
| 12 | 1.67 | brs | 20 |
| 13 | 5.80 | ddd | 129 |
| 14 | 5.20 | ddd | 135 |
| 15a/b | 1.77 / 1.99 | m / m | 43 |
| 16 | 1.83 | m | 29 |
| 17a/b | 1.57 / 1.85 | dd / m | 53 |
| 18 | - | - | 76 |
| 19 | 5.50 | dd | 137 |
| 20 | 5.90 | dd | 127 |
| 21 | 5.62 | dd | 77 |
| 22 | 1.02 | d | 26 |
| 23 | 1.32 | s | 32 |
| 1’ | - | - | 129 |
| 2’,6’ | 7.03 | d | 129 |
| 3’,5’ | 6.82 | d | 114 |
| 4’ | - | - | 159 |
| 24 | - | - | 170 |
| 25 | 2.22 | s | 55 |
| 26 | 3.79 | s | 55 |

**Table S10**: NMR data of 7-deshydroxypyrichalasin H recorded at 400 MHz in $CDCl_3$. Values are in agreement with published data. [8]

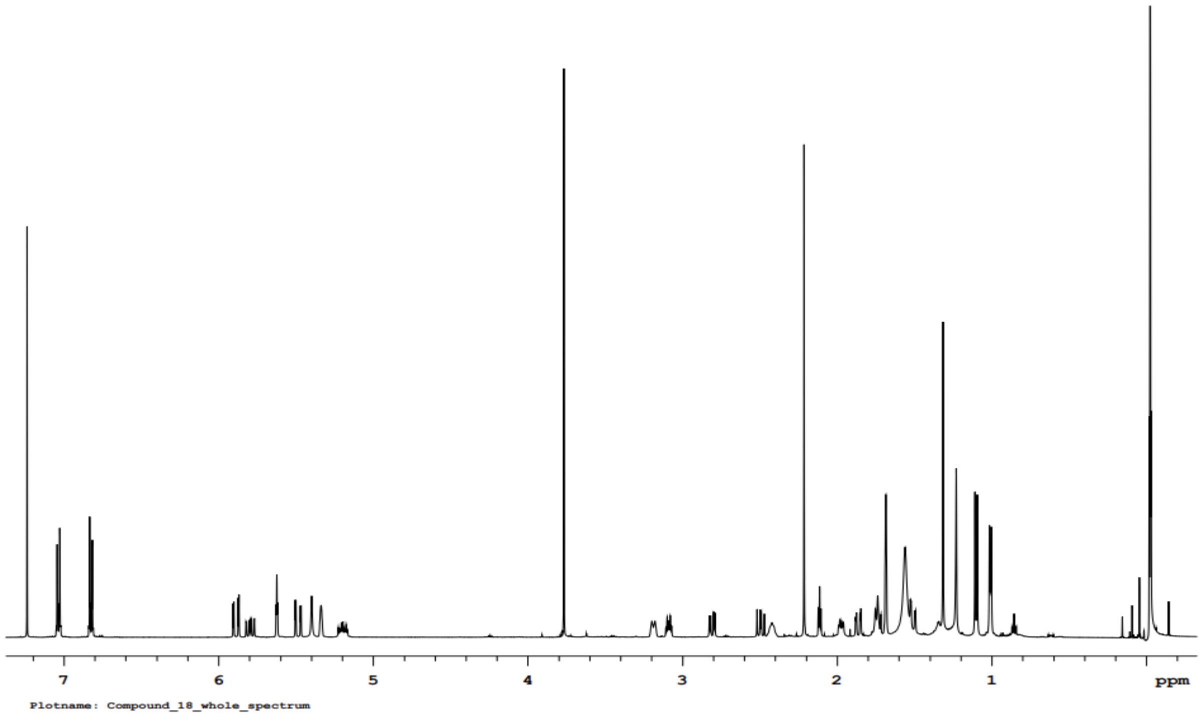


**Figure S69**. $^{1}H$ NMR spectrum (400 MHz, $CDCl_3$) of 7-deshydroxypyrichalasin H.

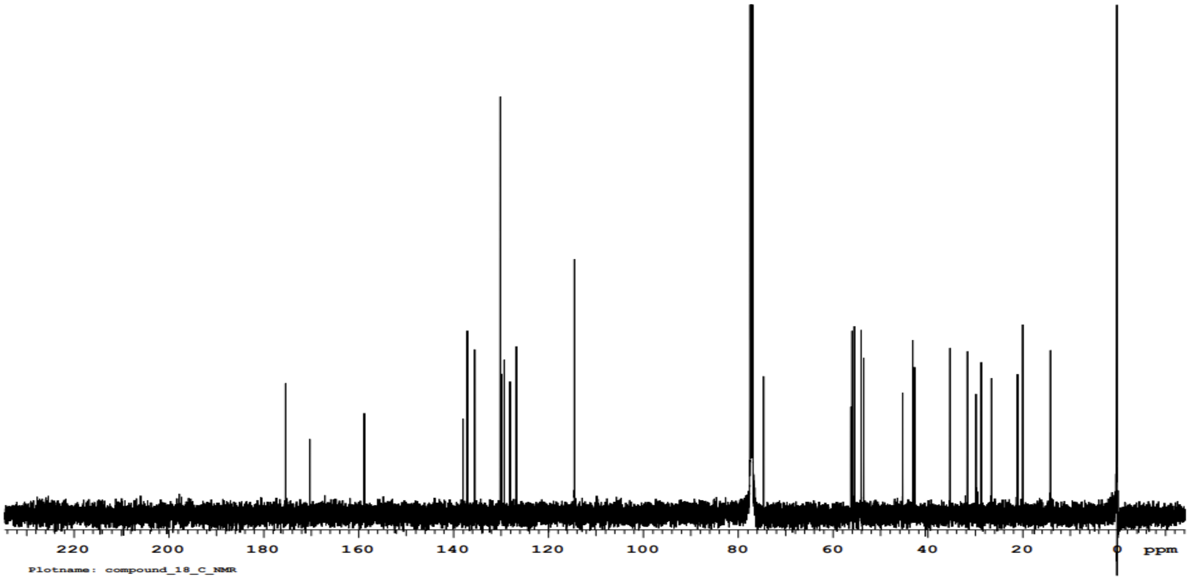


**Figure S70**. $^{13}C$ NMR spectrum (400 MHz, $CDCl_3$) of 7-deshydroxypyrichalasin H.
